# Imputation of ancient canid genomes reveals inbreeding history over the past 10,000 years

**DOI:** 10.1101/2024.03.15.585179

**Authors:** Katia Bougiouri, Sabhrina Gita Aninta, Sophy Charlton, Alex Harris, Alberto Carmagnini, Giedrė Piličiauskienė, Tatiana R. Feuerborn, Lachie Scarsbrook, Kristina Tabadda, Povilas Blaževičius, Heidi G. Parker, Shyam Gopalakrishnan, Greger Larson, Elaine A. Ostrander, Evan K. Irving-Pease, Laurent A.F. Frantz, Fernando Racimo

## Abstract

The multi-millenia long history between dogs and humans has placed them at the forefront of archeological and genomic research. Despite ongoing efforts including the analysis of ancient dog and wolf genomes, many questions remain regarding their geographic and temporal origins, and the microevolutionary processes that led to the diversity of breeds today. Although ancient genomes provide valuable information, their use is hindered by low depth of coverage and post-mortem damage, which inhibits confident genotype calling. In the present study, we assess how genotype imputation of ancient dog and wolf genomes, utilising a large reference panel, can improve the resolution provided by ancient datasets. Imputation accuracy was evaluated by down-sampling high coverage dog and wolf genomes to 0.05-2x coverage and comparing concordance between imputed and high coverage genotypes. We measured the impact of imputation on principal component analyses and runs of homozygosity. Our findings show high (R^2^>0.9) imputation accuracy for dogs with coverage as low as 0.5x and for wolves as low as 1.0x. We then imputed a dataset of 90 ancient dog and wolf genomes, to assess changes in inbreeding during the last 10,000 years of dog evolution. Ancient dog and wolf populations generally exhibited lower inbreeding levels than present-day individuals. Interestingly, regions with low ROH density maintained across ancient and present-day samples were significantly associated with genes related to olfaction and immune response. Our study indicates that imputing ancient canine genomes is a viable strategy that allows for the use of analytical methods previously limited to high-quality genetic data.

## Introduction

Among all domesticated species, dogs (*Canis familiaris*) are of unique public and scientific interest due to their extensive history with humans. Analyses of ancient dog and wolf genomes have advanced our understanding of their evolutionary history (Thalmann et al. 2013; Skoglund et al. 2015; Frantz et al. 2016; Botigué et al. 2017; Ní Leathlobhair et al. 2018; Ollivier et al. 2018; Bergström et al. 2020, 2022; Loog et al. 2020; Sinding et al. 2020, 2020; Da Silva Coelho et al. 2021; Feuerborn et al. 2021; Ramos-Madrigal et al. 2021). However, these insights have been limited by the typically low coverage and degraded nature of ancient DNA (aDNA), which leads to elevated uncertainty in genotype calling and restricts the type of questions that can be confidently addressed (Axelsson et al. 2008; Dabney et al. 2013*b*; Günther and Jakobsson 2019; Günther and Nettelblad 2019). Common approaches to dealing with low-coverage aDNA data include ‘pseudohaploidisation’—the random sampling of an allele at a given site, and genotype likelihoods, which incorporate genotype uncertainty due to read-depth and base quality. However, both these approaches have substantial limitations, as many common methods used in population genomics were designed for high confidence diploid genotypes with low error rates and low missing data.

A method that remains largely unused in canine aDNA studies is imputation—i.e., the statistical reconstruction of missing genetic variants based on haplotype similarity, using high-quality samples available from a large reference database (Das et al. 2018). Unlike pseudohaplodization, which reduces the information content of modern genomes to match the low coverage of ancient DNA, imputation allows for improving the quality of ancient genomes by leveraging information from other genomes. Imputation is being widely used in current analyses including genome-wide association studies (GWAS) using single nucleotide polymorphism (SNP) arrays (Li et al. 2009; Marchini and Howie 2010; Porcu et al. 2013; Quick et al. 2020) and population studies based on low depth genome sequences (Spiliopoulou et al. 2017; Gilly et al. 2018; Hui et al. 2020; Lou et al. 2021; Rubinacci et al. 2021).

Imputation of non-human animals has largely focused on model organisms and livestock, for which large reference panels are most abundant (Yang et al. 2020). Recent advances in computational algorithms have substantially improved imputation quality from low-coverage shotgun genomes (Hui et al. 2020; Rubinacci et al. 2021; Ausmees and Nettelblad 2023). Such methods have produced highly accurate results in ancient samples from species for which large reference panels exist—e.g., humans (Sousa Da Mota et al. 2023) and cattle (Erven et al. 2024). However, species with reference panels lacking ancestral diversity show reduced accuracy (e.g., pigs) (Erven et al. 2022). Imputation of modern dogs has shown promising results as a method to increase SNP density (Hayward et al. 2016, 2019; Jenkins et al. 2021; Buckley et al. 2022; Morrill et al. 2022; Meadows et al. 2023), but the accuracy of imputation has not been previously investigated for ancient canids, nor have results from such been applied to questions of canine migration or domestication.

In this study, we developed an imputation pipeline for ancient dog and wolf genomes using a large reference panel consisting of 1,519 modern canids. We benchmarked its accuracy using ten high-coverage (>10x) ancient and present-day dog and wolf samples representing different ancestries from Europe, Asia, Africa and North America, which we downsampled to lower coverages. We further assessed the impact of imputation on principal component analysis (PCA) and runs of homozygosity (ROH). Our results demonstrate that high accuracy is achieved for coverages as low as 0.5x for ancient dogs and 1.0x for Pleistocene wolves. Based on these results, we imputed a worldwide dataset of 50 ancient dogs and 40 ancient wolves, spanning the last 100,000 years of canine evolutionary history. We observed generally stable levels of inbreeding in dogs over the course of the last 10,000 years, which were notably lower compared to the levels seen in present day samples. We also assessed genomic regions with low ROH density (i.e., ROH deserts) across ancient and present-day samples and observed a significant enrichment for gene ontology terms related to olfactory reception and immunity.

## Methods

### Ancient data curation and assembly

We compiled a set of 82 publicly available ancient dog and wolf genomes from across Eurasia (Skoglund et al. 2015; Frantz et al. 2016; Botigué et al. 2017; Ní Leathlobhair et al. 2018; Bergström et al. 2020, 2022; Sinding et al. 2020; Feuerborn et al. 2021; Ramos-Madrigal et al. 2021), along with nine newly sequenced medieval and early modern period dog genomes from Lithuania and Latvia (Table S1) (total ancient samples=90). The ancient dog samples (n=50) range in date from 100 years BP to more than 10,000 years BP, and the ancient wolf samples (n=40) date from 3,000 BP to more than 100,000 BP (Fig.S1). All genomes had a depth of coverage of at least 0.5x for ancient dogs and 1.0x for ancient wolves, following the results of imputation benchmarking (see section “Imputation Benchmarking”). The median depth of coverage for the ancient dogs was 3.7x (min 0.57x, max 33.3x) and for the ancient wolves it was 2.34x (min 1.0x, max 15.9x) (Fig. S1).

### Archeological samples and context

#### Vilnius Lower castle, Lithuania (KT0033, KT0037, KT0039, KT0041, KT0043, KT0049, KT0052, KT0056)

Vilnius Lower castle was the central residence of the Grand Duke in the capital of the Grand Duchy of Lithuania from the early 14th to the middle of the 17th C AD. The zooarchaeological finds dating back from the 13th to the middle of the 14th C AD reflected the construction stages of the castle, and those of the late 14th to the 15th C AD represent the period of its prosperity. In the early 16th C AD, on the site of the castle, a new palace of the Grand Dukes of Lithuania was built, and this complex survived until the late 17th C AD. The castle was abandoned after a Muscovian attack in the middle of the 17th C AD, and completely demolished in the beginning of the 19th C AD. Canines analysed in this study were found during the archaeological excavations of 1988–2014, in the cultural layers dated to the 13th to 17th C AD. In Vilnius Lower Castle, an abundant zooarchaeological collection (NISP ca 80 000) with numerous dog remains (NISP 590, MNI 51) was collected and analysed. As historical records indicate, hunting was the main function of elite dogs in the Middle Ages and the early Modern Period. Therefore, dogs found in Vilnius Lower Castle and other elite residential environments were most likely used for hunting (Blaževičius et al. 2018; Piličiauskienė et al. 2023).

#### Riga city, Latvia (KT0094)

An almost complete dog skeleton was found in 2006, during archeological excavations (Lūsēns 2008) at the site of a 14th-17th century AD cemetery near the St. Gertruda church at Brivibas Street 42/4 in Riga. Nonetheless, it appears that the dog is not associated with the cemetery. It exhibits a notable pathology - knuckling, also known as carpal laxity syndrome.

### Ancient DNA extraction, library preparation and sequencing

All aDNA laboratory work for the medieval and early modern period dog genomes from Lithuania and Latvia was undertaken in the dedicated ancient DNA laboratory within the PalePalaeogenomics & Bio-Archaeology Research Network (PalaeoBARN), School of Archaeology, University of Oxford. Between 47.7-68.5mg of bone powder was finely drilled from each specimen using a rotary dental drill at low speed or pulverised using a Retsch MM400 dismembrator at low speed. DNA was extracted using a modified version of the (Dabney et al. 2013*a*) protocol, designed specifically for short DNA fragments, but replaced the Zymo-Spin V column binding apparatus with a high pure extender assembly from the High Pure Viral Nucleic Acid Large Volume Kit (Roche 05114403001). Double-stranded Illumina libraries were prepared using the Blunt-End Single Tube (BEST) protocol outlined in (Carøe et al. 2018), and quantitative PCR (qPCR) was used to assess the number of cycles necessary to amplify libraries to the concentration needed for sequencing by amplifying 1 uL of library with LabTAQ Green Hi Rox master mix (Labtech) and adapter-targeted primers on a StepOnePlus Real-Time PCR system (Thermofisher Applied Biosystems). Indexing PCR involved double indexing (Kircher et al. 2012) and used AccuPrime I supermix (ThermoFisher) and the primers described by (Carøe et al. 2018). PCR reactions were purified using AMPure XP beads (Beckman Coulter); fragment distribution was checked on a TapeStation 2200 (Agilent) with D1000 High Sensitivity screentapes and concentration was measured using a Qubit 3.0 (Thermofisher) fluorometer.

Initial screening was performed at the LMU Genzentrum, Munich, Germany on a NextSeq 1000 P2 flowcell (100 bp Single End run). Deeper sequencing was then undertaken at the National Institutes of Health USA on the NovaSeq 6000 Sequencing System with paired end sequencing and 150 bp reads. The data generated for this study have been deposited to the European Nucleotide Archive (ENA) under project number PRJEB73844.

### Ancient genome data preparation

Paired-end data reads were trimmed of adaptors and collapsed using adapterRemoval v2 (Schubert et al. 2016) and mapped with BWA aln v0.7.17 (Li and Durbin 2009; Li 2013) to the CanFam3.1 dog reference genome (Lindblad-Toh et al. 2005) using the following parameters: -l 16500 -n 0.01 -o 2. We used FilterUniqueSAMCons (Kircher 2012) to remove duplicate reads with the same orientation and same start and end coordinates.

### Imputation pipeline

To account for the genotype uncertainty in low-coverage ancient sequences, we phased and imputed the ancient dog and wolf dataset using GLIMPSE v1.1.1 (Rubinacci et al. 2021), which has been shown to produce highly accurate phased haplotypes from ancient DNA, when used with a large and representative reference panel (Sousa Da Mota et al. 2023). The imputation pipeline can be found at https://github.com/katiabou/dog_imputation_pipeline.

### Reference panel

We compiled a large and globally diverse canine reference panel, consisting of 139,268,526 variants and 1,701 whole-genome samples, including modern breed dogs (n=1,395) representing 237 dog breeds, village and indigenous dogs (n=111), New Guinea singing dogs (n=15), dingoes (n=32) and wild canids (n=148). These included grey wolves (n=116), African golden wolves (n=6), African wild dogs (n=3), jackals (n=5), coyotes (n=9), a dhole (n=1), an Ethiopian wolf (n=1), a grey fox (n=1) and red wolves (n=6) (Table S2).

We used BWA mem v0.7.17 (Li 2013) to perform the FASTQ alignment, which was then sorted using samtools v1.12 (Danecek et al. 2021). The GATK v4.1.8.0 MarkDuplicates tool (Van der Auwera and Brian D. O’Connor 2020) was then used to tag duplicate reads. GATK BaseRecalibrator was used to generate BQSR recalibration tables using CF31_dbSNP_v151.vcf as the known sites, followed by the GATK ApplyBQSR tool to apply the recalibrations to the samples. The GATK Haplotypecaller (Poplin et al. 2018) was used to emit all active sites in GVCF mode and generated GVCF files from the BQSR bam file in preparation for cohort calling. The GATK GenomicsDBImport tool was used to collate the GVCFs together. For parallelising purposes, the importation was done in approximately 5 MB intervals using natural gaps in the CanFam3.1 genome. GATK GenotyopeGVCFs was then used on these shards to generate region based VCFs which were then merged using the GATK GatherVcfsCloud tool. The resulting VCF had VQSR recalibration as described in (Plassais et al. 2019).

We filtered for samples with a minimum depth of coverage (DoC) of 8x (Fig. S2), and excluded all boxer breed samples (to avoid reference bias from alignment to the CanFam3.1 assembly). If duplicates of the same sample were present, the lowest coverage member of the pair was removed. This resulted in a final dataset of 1,519 high-quality samples which was used as the imputation reference panel. This included 1,277 breed dogs represented by 228 breeds, 80 village dogs and indigenous dogs, 29 dingoes, 14 New Guinea singing dogs, and 119 wild canids which included 101 grey wolves, 1 dhole, 3 jackals, 1 grey fox, 6 coyotes, 4 African golden wolves, 2 African wild dogs, 1 red wolf and 1 Ethiopian wolf.

We filtered sites using bcftools v1.15.1 (Danecek et al. 2021) to retain only biallelic SNPs which passed variant quality score recalibration with GATK, and removed sites with a fraction of missing genotypes greater than 5%; resulting in 29,480,023 sites in the autosomes. We subsequently phased the reference panel using shapeit v5.0.1(Hofmeister et al. 2023).

### Imputation of ancient dog and wolf dataset

We imputed the ancient dog and wolf dataset per chromosome following the recommended GLIMPSE workflow (Fig. S3) by: i) computing genotype likelihoods for each sample, restricting to the sites and alleles ascertained in the filtered reference panel, using bcftools v1.15.1 (Danecek et al. 2021) ‘mpileup’ function with the flags ‘-I -E -a “FORMAT/DP”’ and the ‘call’ function with the flags ‘-Aim -C alleles’; ii) splitting each chromosome into chunks using a window size of 2 Mb and a buffer size of 200 Kb using GLIMPSE_chunk; iii) imputing each chunk using the genotype likelihoods of each sample, the reference panel haplotypes and the CanFam3.1 genetic map (Campbell et al. 2016) using GLIMPSE_phase; and iv) ligating the chunks of each chromosome using GLIMPSE_ligate. We also carried out phasing of haplotypes with the ‘–solve’ flag using GLIMPSE_sample. We subsequently applied post-imputation filtering based on the imputation accuracy assessment results (see below), removing sites below an INFO score of 0.8 and a minor allele frequency (MAF) cutoff of 0.01 in the reference panel.

### Imputation benchmarking

We benchmarked GLIMPSE to test how accurately it can impute low coverage ancient dog and wolf samples using our reference panel, and to determine the best empirical cutoffs for post-imputation filtering. We chose 10 high coverage (>10x) targets representing different ancestries and time periods; including two late Neolithic European dogs (4,800 BP and 4,900 BP), one North American pre-contact dog (4,157 BP), one historical (60 BP) and one Iron Age (2,000 BP) Siberian dog as well as two present-day village dogs from Nigeria and China (since no ancient representatives of African and Asian ancestry are currently available), and three Pleistocene wolves (16,800 BP, 32,000 BP and 50,000 BP) (Table S3).

We downsampled each high-coverage genome to six lower coverage levels (0.05x, 0.1x, 0.2x, 0.5x, 1x and 2x) using samtools v1.15.1 (Danecek et al. 2021). We then followed the same GLIMPSE workflow as above, imputing each downsampled target individual separately. Modern samples (i.e. those included in the original reference panel) were removed from the reference panel for the benchmarking.

We subsequently used the GLIMPSE concordance tool (GLIMPSE_concordance) to test for concordance between the downsampled imputed genotypes and the high coverage validation genotypes (see validation dataset section below). We assessed how MAF and INFO cutoff scores (0.8, 0.9, 0.95) affected concordance values. INFO scores indicate the level of uncertainty in the posterior genotypes probabilities of each imputed site. We computed concordance across MAF and INFO scores using both all sites and transversions only. We ran the GLIMPSE_concordance tool using the following flags ‘-minDP 8 -minPROB 0.9 –af-tag AF –bins 0.00000 0.00100 0.00200 0.00500 0.01000 0.05000 0.10000 0.20000 0.50000’, as suggested in the GLIMPSE manual (https://odelaneau.github.io/GLIMPSE/glimpse1/).

The GLIMPSE concordance tool also provides metrics of genotyping errors for homozygous alternative, heterozygous and homozygous reference alleles. It also outputs the non-reference discordance (NRD) metric, which only takes into consideration imputation errors at alternative alleles by excluding confidently imputed homozygous reference alleles. This is equal to:

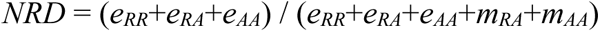

where *e_RR_*, *e_RA_*, *e_RR_* are the mismatches at homozygous reference, heterozygous and homozygous alternative alleles respectively, whereas *m_RA_* and *m_AA_* are the matches at heterozygous and homozygous alternative alleles. We also tested how the amount of canid haplotype diversity present in the reference panel influenced imputation accuracy by using a dog-only reference panel in a separate imputation analysis (n=1,399 and 18,497,052 sites).

### Validation dataset filtering

To limit the impact of genotyping errors in our benchmarking pipeline, we applied the following filters on the 10 high coverage samples used for benchmarking while using the bcftools ‘mpileup’ and ‘call’ functions, following (Sousa Da Mota et al. 2023): i) reads with mapping and base quality below 30 (-q 30, -Q 30) were removed and the ‘-C 50’ option was used to downgrade mapping quality for reads containing excessive mismatches; ii) sites with QUAL lower than 30 were excluded; iii) sites with extreme values of depth of coverage (i.e., sites with a depth of coverage greater than twice the mean genome-wide depth, and sites with a depth below either 8x or one third of the mean depth of coverage (i.e., max(DoC/3, 8)), whichever is greater were also excluded; and iv) heterozygous sites at which the one of the two alleles was found in less than 15% or more than 85% of the reads using bcftools v1.15.1 ‘view’ and the flags ‘–exclude ’GT=“het” && ((INFO/AD[1] / INFO/DP < 0.15) || (INFO/AD[1] / INFO/DP > 0.85))’.

### PCA of imputed samples

#### Downsampled target samples

We next assessed how imputation of low coverage samples would affect their placement in PCA space in comparison to pseudohaploid data. To do this, we called pseudohaploid genotypes in the downsampled target samples using the -doHaploCall function in angsd v0.94 (Korneliussen et al. 2014), the -doCount 1 option, filtering for a minimum base and map quality of 30 (-minMapQ 30, -minQ 30), trimming five base pairs at the beginning and end of each read (-trim 5) and restricting to transversion sites (-noTrans 1). We applied a minimum MAF (0.01) and INFO score (0.8) cutoffs in the imputed samples (based on the benchmarking, see results) to assess how this compares to unfiltered imputed genotypes. After filtering the pseudo-haploid dataset for sites present in the filtered reference panel, we merged it with the high-coverage genotyped validation samples, the imputed samples (high-coverage and downsampled) and the filtered reference panel.

We subsequently carried out PCA using smartpca eigensoft v8.0 (Patterson et al. 2006). For our ancient dog PCA we used a reference panel of 502 present-day dogs, and for our ancient wolf PCA we used a reference panel of 95 present-day wolves (Table S2). We projected the imputed and pseudohaploid replicate of each target sample along with its genotyped high coverage version onto the PCA using the lsproject option. We subsequently estimated the sum of weighted PC distances between each downsampled target (imputed and pseudohaploid) and the genotyped high-coverage counterpart (used as the ground truth) across the first 10 principal components.

#### Imputed ancient dog and wolf dataset

Prior to the PCA of the full imputed ancient dataset, we merged all imputed ancient samples into the same VCF and re-calibrated the INFO scores in order to maintain a consistent filtering of sites across individuals. We applied MAF≥0.01 and INFO≥0.8 cutoffs based on the benchmarking results. We again applied the smartpca tool of eigensoft, this time using the imputed and present-day reference panel samples to create the first 10 principal components. The present-day reference panel was filtered only for sites present in the merged imputed dataset. We ran a PCA for dog and wolf samples separately, using either present-day dogs or present-day wolves respectively.

### Runs of homozygosity of imputed samples

Prior to estimating ROH, we applied MAF (0.01) and INFO score (0.8) cutoffs on each of the imputed samples (i.e., prior to INFO score recalibration in the merged callset). We used the PLINK v1.9 (Chang et al. 2015) (www.cog-genomics.org/plink/1.9/) --homozyg tool to estimate ROH, carrying out two runs: i) only including transversions and ii) including both transversions and transitions. In both runs, the following parameters were set: --homozyg-density 50, --homozyg_gap 500, --homozyg_kb 500, -- homozyg_snp 50, --homozyg_window_het 1, --homozyg_window_missing 5, --homozyg_window_snp 50, --homozyg_window_threshold 0.05. We chose these parameters following published recommendations for ancient samples (Ceballos et al. 2021; Sousa Da Mota et al. 2023). We chose the PLINK parameter -- homozyg-window-het 1, consistent with the ancient DNA literature (Schroeder et al. 2018; Ceballos et al. 2021; Sousa Da Mota et al. 2023) and with some present-day studies (Clark et al. 2019; Aramburu et al. 2020; Lavanchy and Goudet 2023). However, we note that this configuration allows an unlimited number of heterozygous SNPs across a putative ROH block, as long as no more than one heterozygous SNP appears in a sliding window of size --homozyg-window-snp 50. As such, the biological interpretation of these loci should be that they are regions of low diversity, rather than strictly uninterrupted runs of homozygosity.

This applies to all published literature where no upper bound is specifically set with the flag --homozyg-het. The same set of parameters was used for the downsampled imputed target samples, the high coverage genotyped target samples, the full imputed ancient dataset and the reference panel. For the ROH analysis, a MAF 0.01 filter was applied to the reference panel.

### ROH estimates using ROHan

As part of our benchmarking approach, we compared our results to ROHan v1.0 (Renaud et al. 2019) - a method designed to infer ROHs on ancient medium-coverage data (at least 7X) that has not been imputed. We used ROHan to infer ROHs on the non-imputed downsampled and HC targets to compare against the inferred ROH on the imputed ones from our pipeline. For the ancient genomes, we first ran the ‘bam2prof’ utility of ROHan to obtain the deamination pattern from the first 5 base pairs of the 5’ and 3’ prime end at each downsampled coverage and consider a minimum base quality of 20 (-minq 20 -minl 5). The resulting deamination profile of each sample at each coverage was then run along with the BAM file in ROHan (via option –-deam5p and --deam3p) using the default parameters, except for the number of heterozygous sites (--rohmu) which was set to 4 x 10^-5^ and the sliding window (--size) which we ran on the default 1Mbp (we also tried a smaller window size to match the window size of 500Kbp of our imputation pipeline, but this resulted in lower accuracy estimates). The modern genomes were run similarly without the deamination profile option.

### ROH accuracy assessment

To assess the accuracy of inferred ROH blocks in the downsampled imputed samples, we estimated the ROH overlap with the high-coverage samples with respect to the total number of segments and total length of overlapping bases, using the GenomicRanges v1.50.2 R package (Lawrence et al. 2013). For both approaches, we calculated true positives (TP), false positives (FP), and false negatives (FN). Additionally, for the length-based approach, we calculated true negatives (TN). For the segment-based approach, we used the F1-score metric (F1= 2*(precision*sensitivity)/(precision+sensitivity)) which is calculated based on sensitivity (correct positive predictions relative to total actual positives - TP/(TP+FN)) and precision (correct positive predictions relative to total positive predictions - TP/(TP+FP)), where 0 indicates no ROH overlap and 1 shows perfect overlap.

For the length-based approach we used the F1-score and the Mathews correlation coefficient (MCC), which takes into consideration all four confusion matrix categories (FN, FP, TN, TP), allowing equal contribution of positives and negatives. This is considered to be more reliable than the F1-score (Chicco and Jurman 2020), as a high score is obtained when all four confusion matrix categories obtained good results (TP, TN, FP, FN), in comparison to the F1-score which primarily weighs the correct positive predictions. To calculate the MCC, we used the mcc function of mltools (Ben Gorman 2018) in R. We then estimated the normalised Mathews correlation coefficient (nMCC=(MCC+1)/2), where a value equal to 0.5 indicates a random prediction and a value closer to 1 represents complete overlap. Finally, using the length-based overlap estimates, we calculated specificity (TN/(TN+FP)), sensitivity (TP/(TP+FN)) and false discovery rate (FDR=FP/(TP+FP)) on the total outcome.

### ROH estimates across space and time

We estimated the total number and total length of ROH for each imputed individual, as well as the inbreeding coefficient (*F_ROH_*), which is equal to the total length of ROH, divided by the total genome length. We estimated these metrics for both long (>=1.6Mb) and short (<1.6Mb) ROH blocks separately, since they can be indicative of different demographic events (Ceballos et al. 2018). We chose these cutoffs based on the distribution of ROH lengths calculated for ancient and modern dogs.

In order to visualise fluctuations in inbreeding patterns through space and time, we grouped the ancient dog samples into three geographic regions: Europe, the Arctic and the Near East (Table S1). ROH estimates of present-day dogs (breed dogs and village dogs) from these three regions were included for comparison. Within the Near Eastern cluster, we also included African and Indian village dogs, as well as African modern breeds. We also grouped the imputed ancient wolves into three populations: Pleistocene, Holocene Eastern Eurasia and Holocene Western Eurasia. This grouping was based on previous work showing that Pleistocene wolves were a panmictic population and that population structure and differentiation increased during the Holocene (Bergström et al. 2022). Present-day wolf samples from east and west Eurasia as well as from North America were included for comparison.

We subsequently carried out per population Mann–Whitney U tests using the wilcox.test function in R to test for significant differences between dog populations and time periods. For this we used the three geographic groupings (Europe, Arctic and Near East) and then carried out tests between each pairwise combination of: i) ancient dog populations, ii) present-day and ancient dog populations and iii) present-day dog populations. We carried out these tests on all, short and long ROH.

### Prevalence of ROH in ancient and modern samples

To further characterise patterns of ROH presence and absence in ancient and present-day populations, we used the windowscanr v0.1 R package (https://github.com/tavareshugo/WindowScanR) to estimate the prevalence of ROH across the genome, following the published approach (Stoffel et al. 2021, https://github.com/mastoffel/sheep_ID). We split the genome into 500 Kb windows and estimated the percentage of samples which contained a ROH in each window. This was carried out separately for the 50 ancient dogs, the 40 ancient wolves, a subset of present-day dogs (n=502) and a subset of present-day wolves (n=95) from the reference panel. We excluded windows with extremely high or low read depth, as they may be enriched for structural variants (e.g., copy number variants or segmental duplications) or mapping errors. We identified these outlier windows by estimating the average depth of coverage per window (n=4,385 total windows) using all ancient dog or wolf samples, and excluded all windows with a depth of coverage outside of two standard deviations from the mean. We also excluded windows which did not have any sites present in the imputed dataset. Based on these cutoffs, we retained 98% of the total windows for dogs (n=4,300) and 98.5% for wolves (n=4,323).

We defined ROH deserts as windows in which <5% of the ancient samples and <5% of the present-day samples had an ROH. We carried out gene enrichment analysis on these ROH deserts using the GOfuncR package (Grote 2023) to test for an over-representation of genes related to specific biological categories among the genes that fell within ROH deserts. To this end, we applied the hypergeometric test for GO enrichment, correcting for gene length. We removed the 84 dog and 61 wolf outlier windows with high or low read depths from the background genomic regions. We used the ’org.Cf.eg.db’ OrgDb package for GO-annotations and the ’TxDb.Cfamiliaris.UCSC.canFam3.refGene’ TxDb package for gene-coordinates. To correct for multiple testing and test interdependency, we computed the family-wise error rate (FWER) for each GO-category, using 1000 randomised sets of the data. In each randomised set, the background and candidate genes are permuted, and new p-values are computed. For a given GO-category, the FWER is then the fraction of the randomised sets whose lowest p-value is lower than or equal to the original p-value of the GO-category. For example, a FWER of 0.1 for a GO-category “X” means that, in 10 out of 1000 randomised sets of the data, the set’s minimum p-value is smaller than or equal to the original p-value of “X” (see GOfuncR’s online manual for an extended explanation).

## Results

### A new pipeline for ancient dog genome imputation

We implemented a fully reproducible imputation pipeline using the GLIMPSE software and a reference panel of over 1,500 modern canids, available at https://github.com/katiabou/dog_imputation_pipeline. We applied this pipeline to the largest dataset of ancient dog and wolf genomes analysed to date, including nine new dog genome sequences. We tested the accuracy of our imputation pipeline by downsampling 7 high-coverage ancient and present-day dog samples and 3 Pleistocene wolf samples, and assessed the concordance of the imputed genotypes against the original high-coverage genotypes. Based on the benchmarking results, we subsequently imputed 50 ancient dog and 40 ancient wolf genomes (total=90).

### Imputation accuracy assessment

Our analysis showed high concordance when imputing dogs as low as 0.5x (r^2^ > 0.9) and wolves as low as 1x coverage (r^2^ > 0.8) (Fig.1, Fig. S4-S13). As expected, reduced accuracy (r^2^ < 0.8) was observed at lower levels of coverage and at sites with a lower MAF (<0.01) in the reference panel. Applying an INFO score cutoff of 0.8 removes low confidence imputed sites, which increases concordance, although no further improvement was noticed under higher INFO score thresholds (0.9 and 0.95). All dog samples with ≥0.5x coverage and all wolf samples with ≥1x coverage reached a r^2^ plateau for sites with a MAF of ≥0.05, and in many cases as low as 0.01, demonstrating the accuracy with which GLIMPSE can impute common variants (Fig. S4-S13).

Among the various dog ancestries tested, ancient and historical Siberian and modern African and Asian village dogs showed high accuracy levels (r^2^ > 0.9), even at 0.2x coverage with a MAF cutoff of 0.01 (r^2^ > 0.9). One of the Pleistocene wolves (CGG33) showed similar accuracy levels (r^2^ > 0.9) for coverages from 1x and above.

**Fig. 1:**
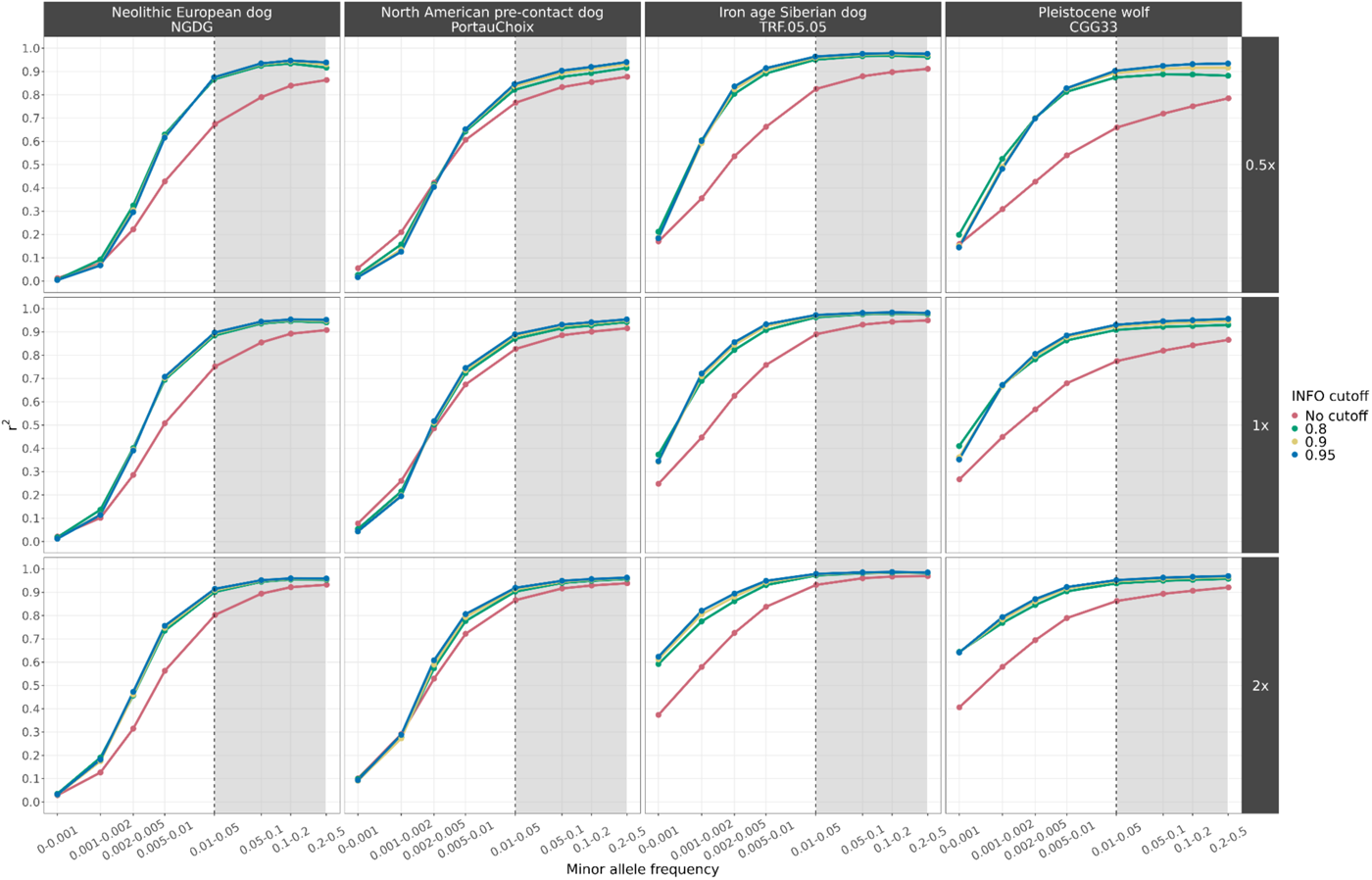
Squared correlation (r^2^) between imputed genotypes by GLIMPSE and highly confident called genotypes for four high coverage samples (three ancient dogs and one Pleistocene wolf), at three downsampled coverage values (0.5x, 1x, 2x) and across different MAF bins. Each colour depicts the accuracy for a given INFO score cutoff. Red: no cut-off, Green: 0.8, Yellow: 0.9 and Blue: 0.95. Sites belonging to the MAF bins within the grey shaded area were retained after post-imputation filtering.

We observed low genotyping error rates (<10%) for most dog samples with coverage ≥0.5x when applying an INFO score cutoff of ≥0.8 (Fig. S14). Overall, the error rate for homozygous reference and heterozygous genotypes was lower than 5% in both 0.5x dogs and 1x Pleistocene wolves. The 0.5x Port au Choix individual, a North American pre-contact dog, possessed the highest level of errors amongst heterozygous genotypes (12.1%).

Genotyping errors for homozygous alternative genotypes were higher than error rates for homozygous reference and heterozygous genotypes in all samples, with estimates ranging from 5.8% in a 0.5x dog downsampled genome from Iron Age Siberia to 12.4% in the 0.5x downsampled Port au Choix individual. Genotyping error rates for homozygous alternative genotypes were higher in Pleistocene wolves than in dogs with error values between 12.1% and 28.3% for 1x Pleistocene wolves (Fig. S15).

We also looked at another measure of error, the non-reference discordance (NRD) rate, which gives weight to the incorrectly imputed alternative allele sites (homozygous or heterozygous) and not the homozygous reference sites, which represent the majority of sites. When applying an INFO score cutoff of 0.8, all 0.5x imputed dog samples showed NRD rates <10%, apart from the Port au Choix individual (NRD=18.4%). The NRD rates for 1x Pleistocene wolves ranged from 7.9% to 15.9%.

For the majority of ancestries tested, dogs with ≥0.5x coverage and wolves with ≥1x coverage exhibited less than a 1% difference in genotyping errors associated with transversions only versus all sites (Fig. S16-S20). The Port au Choix North American pre-contact dog was the only sample that showed a decreased genotyping error rate when restricting to transversions compared to including all sites across all tested coverages (Fig. S17, S19). Specifically, at 0.5x coverage and 0.8 INFO score cutoff, genotyping rates decreased from 12.1% to 0.8% for heterozygous alternative sites, 12.3% to 11.4% for homozygous alternative sites and from 18.4% to 12.3% for the NRD rate. These results suggest that above our coverage cutoffs for all tested ancestries, apart from North America pre-contact dogs, there is no benefit to restricting imputed sites to transversions only.

In order to test whether the diversity of canid species within the reference panel influences imputation accuracy, we also ran our pipeline using a reference panel consisting only of dogs. Imputation accuracy of the 1x Pleistocene wolves decreased going from r^2^ > 0.8 to r^2^ < 0.74 for sites within a MAF bin of 0.01-0.05 (Fig.S21). Furthermore, imputed sites associated with the lower MAF bins increased in accuracy for dogs, e.g. from r^2^=0.63 to r^2^=0.75 for sites within a MAF bin of 0.005-0.01 for a Neolithic European dog. Despite this, the overall number of sites retained after applying INFO score and MAF cutoffs was lower compared to using the all-canid reference panel. For example, for a Neolithic European dog, the number of sites reduced from 8,003,059 to 6,954,516, and for a Pleistocene wolf from 6,156,410 to 5,142,128 when using a dog only reference panel (Fig. S22). Using the dog only reference panel, all 0.5x dog and 1x wolf samples with an INFO score cutoff of ≥0.8 showed <5% error rates for homozygous reference and heterozygous sites, similar to the results obtained using the full reference panel (Fig. S23, S24). The Port au Choix dog showed the highest errors for heterozygous genotypes (9.1%), which was, however, lower than the errors observed when utilising the all canid reference panel (12.1%). Genotyping errors for homozygous alternative sites increased when using the dog reference panel, with the lowest error rate increasing from 8.3% to 8.4% (Historical Siberia), and the highest error rate increasing from 12.3% to 14.7% (Port au Choix) for 0.5x dog samples. For 1x Pleistocene wolves, the lowest genotyping error rate for alternative sites increased from 12.1% to 40.1%, and the highest increased from 28.3% to 56%. NRD rates remained <10% for all dog samples, apart from the Port au Choix dog which increased from 18.4%, when using the full panel, to 19% when using the dog only panel. The lowest NRD rates for 1x Pleistocene wolves increased from 7.9% to 20.7%, whereas the highest rates increased from 15.9% to 30.1%. Given these results all subsequent analyses were based on imputation using the full reference panel.

Based on our results we decided to include imputed samples with at least 0.5x coverage for ancient dogs and 1x for ancient wolves in subsequent analyses, while filtering for sites with INFO scores of at least 0.8 and MAF above 0.01. Considering the potential loss of informative sites when filtering only for transversions (30.76% of all sites), we chose to keep all sites within the imputed dataset. Finally, due to the elevated genotyping error that the imputed North American pre-contact sample showed, we did not impute any dogs assigned to this population (n=1, Table S1).

### PCA of downsampled imputed and non-imputed samples

To further assess the accuracy of the imputed genotypes, we carried out a PCA using each high coverage sample as the ground truth. We then calculated the sum of weighted PC distances between each projected sample and their corresponding high coverage samples across 10 PCs (Fig. 2, Fig. S25-S34). We tested this on the pseudohaploid samples, the imputed filtered (MAF ≥0.01 and INFO score ≥0.8), and imputed non-filtered samples.

**Fig. 2:**
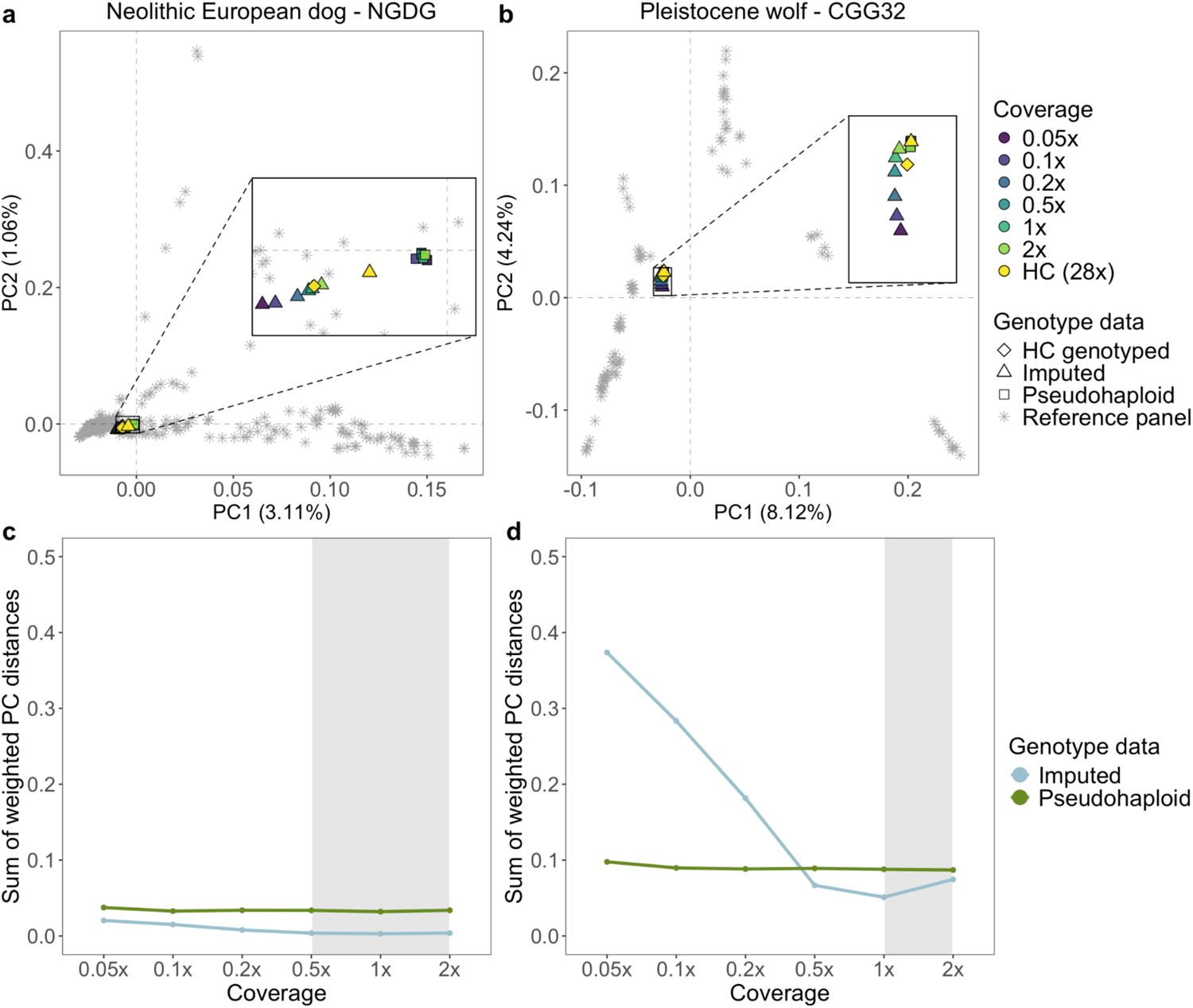
a, b) Principal component analysis demonstrating the placement of the non-filtered imputed Newgrange Neolithic European dog and the CGG32 Pleistocene wolf against their corresponding pseudohaploid counterpart in PCA space across all tested downsampled coverages. PCs were created using modern dog or wolf samples from the reference panel, and all versions of the ancient target sample were projected onto them. c, d) Sum of weighted PC distances for each imputed (blue line) and pseudohaploid (green line) sample relative to the high coverage ground truth sample across all tested coverages. The grey shaded area corresponds to the coverage cutoffs for dogs (0.5x) and wolves (1x) HC: high coverage.

For dogs, we noticed a better placement of the imputed samples (both filtered and unfiltered) in PCA space, compared to the pseudohaploid versions, for the majority of samples with coverages ≥0.5x. The PC distance between the 0.5x pseudohaploid dog samples and the ground truth ranged from being 1.2 times greater (for Iron Age Siberia, TRF.05.05) to 4.9 times greater (for Neolithic Europe, NGDG) compared to the distance between the 0.5x filtered imputed samples and the ground truth. Three imputed ancient samples showed better placement than the pseudohaploid genotypes for all tested coverages (Fig.2, Fig. S25-S27): a North American pre-contact dog (Port au Choix) and two Neolithic European dogs (NGDG and SOTN01). When applying post-imputation filters, the placement of the imputed Pleistocene wolves performed worse across all coverages compared to their pseudo-haploid counterparts. In turn, imputed 1x and 2x samples without any post-imputation filtering were on average 2 and 1.2 times closer to the ground truth compared to their corresponding pseudo-haploid calls (Fig. 2, Fig. S32-S34).

### ROH in downsampled imputed and non-imputed samples

Finally, we compared estimated ROH between the imputed downsampled and high-coverage genotypes for all autosomes. Overall, overlapping ROH estimates varied among samples depending on the metric used (F1-score or nMCC), the reference used to estimate overlap (segment based or total length) and the sites included (transversions or transversions+transitions) (Fig. 3, Fig. S35-S44). Higher nMCC and F1-scores were observed when restricting the analysis to transversions only, with the highest difference observed for Port au Choix, going from values ≤0.75 to >0.75 for coverages ≥0.5x (Fig. S35b-S44b). Both nMCC and F1-score estimates followed similar trajectories across coverages.

**Fig. 3:**
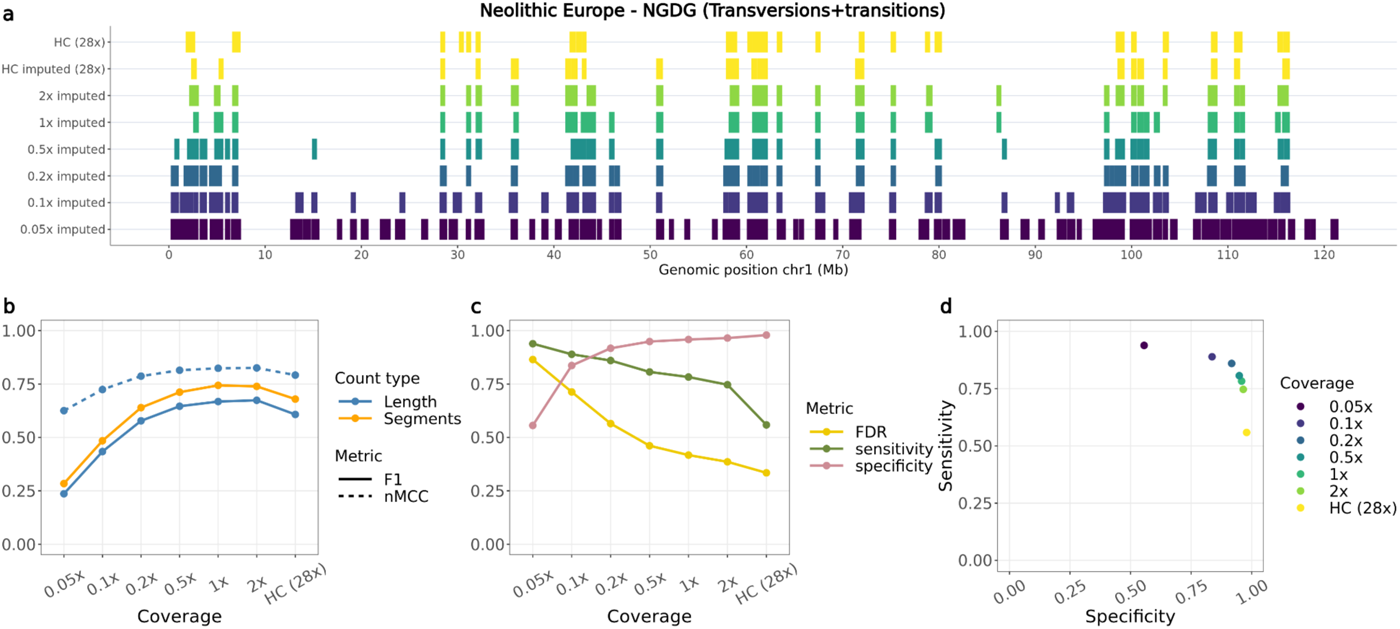
Overlap of ROH called from the Newgrange Neolithic European dog for each imputed downsampled replicate using ROH estimates from the ground truth. a) ROH called across the six tested coverages and the high coverage imputed and genotyped (ground truth) sample on chromosome one, including transversions and transitions. b) Accuracy of recovering ROH across all tested coverages based on total length in bp (blue lines) and total number of segments (orange line) using the F1-score (solid line) and normalised Matthew correlation coefficient (nMCC) (dotted line). c) FDR, sensitivity and specificity measurements based on the total length of recovered ROH per coverage. d) Sensitivity plotted against specificity estimated based on the total length of recovered ROH across all tested coverages. HC: High coverage.

Specificity scores (true negative rates) showed consistently high values (>0.8) across all tested individuals at coverages >0.2x. Sensitivity scores (true positive rates) typically showed a decreasing pattern with increasing coverage, starting from >0.8 at 0.05x coverage and decreasing to <0.1 at 2x coverage in the most severe scenario (Fig. 3c,d, Fig.S35c,d-S44c,d). Increased sensitivity at lower coverages seemed to be due to decreased false negatives at the cost of increased false positives (therefore lower specificity).

When compared to the results from ROHan on non-imputed data, the ROH inferred from the imputed samples presented consistently higher nMCC scores, specificity estimates and lower false discovery rates (Fig. S45-54). For cases below the recommended coverage (7x) at which ROHan is supposed to be used for ancient samples, ROHan would highly overestimate the total length of ROH, as reflected by the high sensitivity and false discovery rates and low specificity estimates observed when looking at samples between 0.5x and 2x (Fig. S45-S49, S52-S54). An increase in nMCC scores was observed in the high coverage samples (11.2x-28x). However, this was due to an underestimation of ROH as shown in the high specificity and low sensitivity rates. Even though the high coverage samples were above the recommended coverage threshold for ROHan, ROH inferred from the high coverage imputed samples consistently showed higher nMCC scores and sensitivity estimates.

### Imputed ancient dog and wolf dataset

Based on our assessment of imputation accuracy, we imputed 50 ancient dog genomes with at least 0.5x coverage and 40 ancient wolves with at least 1x coverage (Fig. S1). After merging all samples, we recalibrated the INFO scores and filtered for sites with an INFO score above 0.8 and for sites with a MAF above 0.01 in the reference panel, leading to a dataset of 10,992,085 SNPs. We subsequently merged the imputed dataset with a subset of samples from the reference panel (n=502 dogs and n=95 wolves) for downstream analyses. Visualising our imputed dog samples in PCA space, we observed a geographic grouping of present-day and ancient samples, forming four main clusters: European, African-Near East-India, Arctic and East Asian (Fig. S55).

### ROH in ancient dogs and wolves

ROH were estimated for the ancient imputed and present-day samples using the same parameters in PLINK. Both transition and transversion sites were included. We estimated the total number and total length of ROH, as well as the ROH-based inbreeding coefficient for: i) all ROH, ii) short ROH (<1.6Mb) and iii) long ROH (≥1.6Mb) (Fig. 4, Fig.S56-S62).

Overall, we observed remarkable stability in inbreeding for dogs during the past 10,000 years, until the beginnings of modern breed formation, which led to a substantial increase in the total number and length of ROH segments (Fig. 4, Fig. S58, S59, S63, Table S5). Among ancient dogs, the highest inbreeding coefficients were calculated for Arctic and European individuals. Eight ancient dog samples from these regions had >10% of their genome located within an ROH (*F_ROH_*>0.1) (Fig. 4): An early modern period Lithuanian dog (153 BP, *F_ROH_*=0.24), an Iron Age and a historical dog from the Iamal-Nenets region (1,111 BP, *F_ROH_*=0.16 & 93 BP, *F_ROH_*=0.15), a Mesolithic dog from the Veretye site in Western Siberia (10,930 BP, *F_ROH_*=0.14), a Neolithic dog from Croatia (4,900 BP, *F_ROH_*=0.11), a historical dog from the Bulgunnyakhtakh site in Northeast Siberia (100 BP, *F_ROH_*=0.1), a Mesolithic dog from Zhokhov island in Eastern Siberia (9,515 BP, *F_ROH_*=0.1) and a Swedish Pitted Ware sample from the island of Gotland (4,800 BP, *F_ROH_*=0.1).

**Fig. 4:**
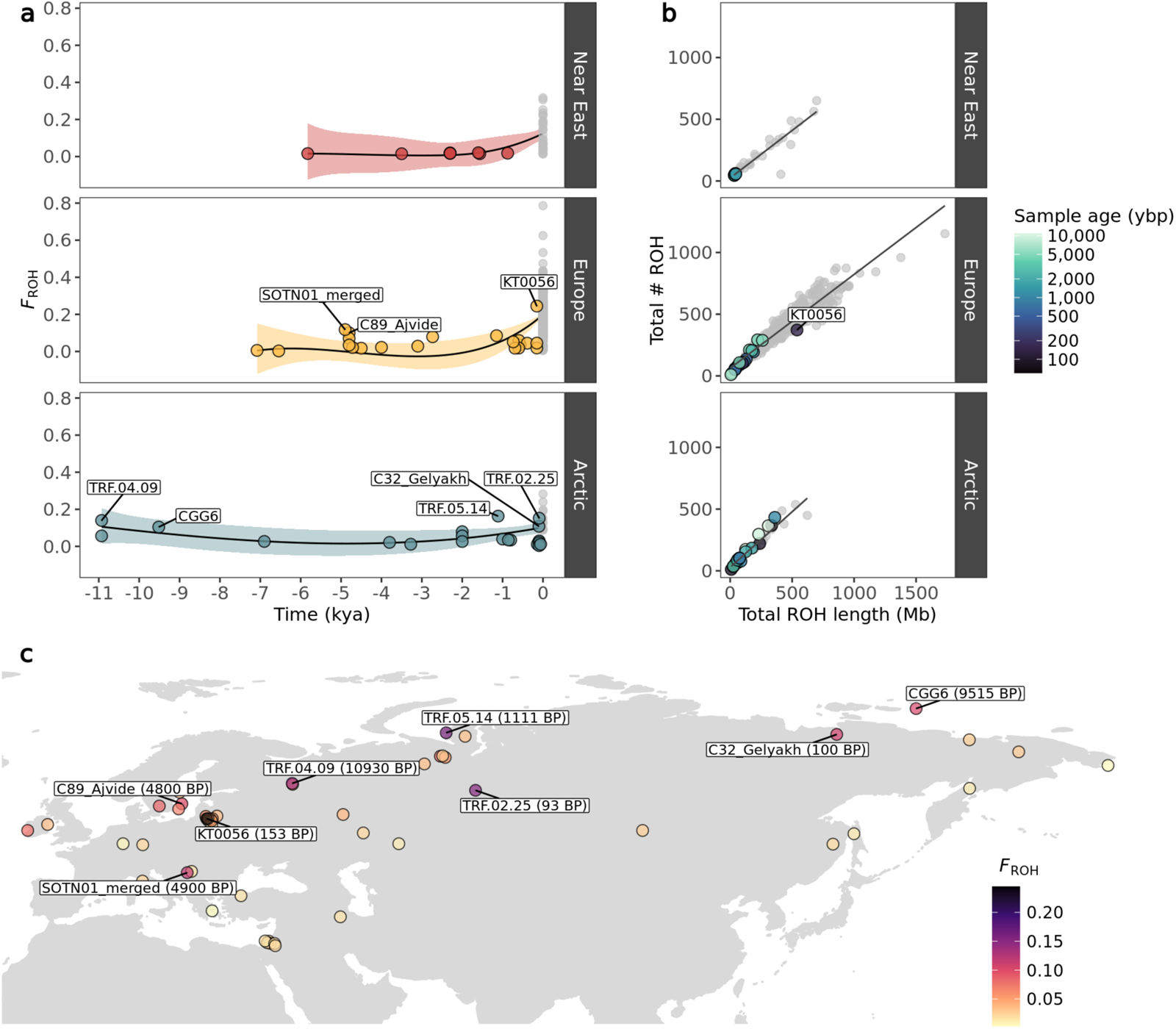
a) Genomic inbreeding coefficient (F_ROH_) of imputed and modern dogs plotted as a function of time, calculated based on ROH. Imputed samples are coloured based on their geographic grouping, while modern samples are coloured in grey. A lowess regression was applied with each coloured shaded area depicting the standard error. b) Total number of ROH segments plotted against total ROH length for the imputed dogs. Colours correspond to age of imputed samples in years before present, while modern samples belonging to each dog group are coloured in grey. c) Map of imputed dog samples coloured by their inbreeding coefficient (F_ROH_). Samples with F_ROH_ values above 0.1 are indicated.

In Europe, we observe a notable increase in inbreeding around 5,000 BP, with three dogs showing increased inbreeding coefficients, primarily due to a higher presence of short ROH segments (Fig. 4, Fig. S58): two individuals from the island of Gotland in Sweden dated to 4,800 BP, and a Croatian dog dated to 4,900 BP. A Lithuanian dog from 153 BP showed the highest *F_ROH_* among ancient dogs (0.24), with an inbreeding coefficient substantially higher compared to eight other dogs from the same region and time period. The high *F_ROH_* of this sample seems to be driven predominantly by the presence of long ROH (≥1.6Mb) (Fig. S58). The ancient Arctic dogs showing highest *F_ROH_* coefficients did not follow a specific temporal or geographic pattern. Ancient Near Eastern dogs showed the lowest *F_ROH_* coefficients with minimal fluctuations in inbreeding levels until the emergence of modern breeds.

Ancient Near Eastern *F_ROH_* estimates differed significantly from ancient European (Mann-Whitney W=28, p<0.05) and Arctic (Mann-Whitney W=33, p<0.05), whereas ancient Arctic and ancient European did not differ statistically from each other (Mann-Whitney W=216, p=0.92). Present-day European dog *F_ROH_* levels differed significantly from present-day Near Eastern (Mann-Whitney W=2735, p<0.05) but not from present-day Arctic (Mann-Whitney W=2637, p=0.055). Present-day Near Eastern and Arctic dogs did not show significant differences (Mann-Whitney W=121, p=0.48) (Table S4).

The imputed wolves showed substantially low *F_ROH_* (<0.04) compared to present-day wolf populations (>0.5), with minimal fluctuations until modern times (Fig. S60-S62, S64, Table S5). The *F_ROH_* levels in Pleistocene wolves remained low (<0.02), with only three samples showing *F_ROH_*>0.01, which is driven by short *F_ROH_* (Fig. S61, S62). It is worth noting that some present-day wolves (from Sweden, Norway and Mexico) showed higher *F_ROH_* levels than present-day breed dogs (Fig.S59, S62, Tables S5).

### Frequency of ROH across the genome of ancient and present-day dogs

We estimated the prevalence of ROH across the genome of ancient and present-day dogs and wolves in 500 Kb windows. Figures 5 and S65 show the percentage of ancient and present-day dogs and wolves with an ROH in each window throughout all autosomes. Yellow regions indicate windows with low ROH frequency (ROH deserts) and purple regions higher ROH frequencies. Grey coloured regions indicate windows with average depth of coverage above or below the mean ± 2*std (Fig. S66, S67), and were not included in the ROH frequency estimation. Ancient dogs showed a substantially lower prevalence of ROH across the genome than modern dog breeds. The majority of windows containing ROH were shared with less than 10% of all ancient dogs and approximately 20% of present-day dogs. When examining ROH deserts (i.e., windows for which <5% of the ancient and <5% of the present-day samples shared an ROH segment), we found 133 windows (3.1% of total windows included). The highest signals from the gene enrichment analysis showed an over-representation of genes related to olfaction and immunity (FWER≤0.2) (Table S6). When examining ancient wolves, few samples shared an ROH in the same window (Fig. S65), whereas 200 windows were identified as ROH deserts (4.6% of total windows included). Gene enrichment analysis also supported an over-representation of olfaction and immunity genes, though FWER values for the top categories were much higher in this case (FWER≤0.4) (Table S7).

**Fig. 5:**
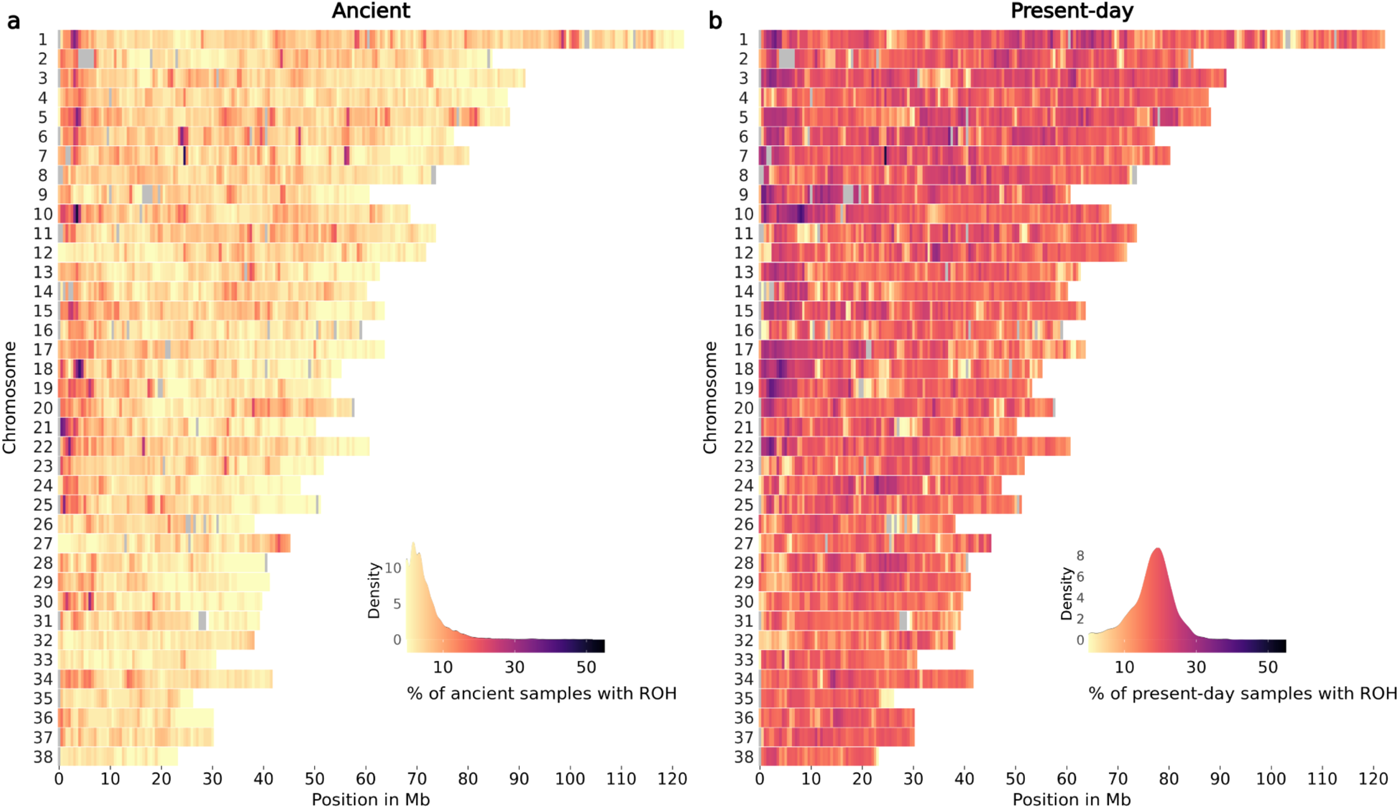
ROH across all chromosomes of a) ancient dogs and b) present-day dogs. The colour legend represents the % of samples which have a ROH at each genomic position, with more yellow regions representing ROH deserts and more purple regions representing ROH islands. Grey coloured regions indicate windows with an average depth of coverage estimated from all ancient dog samples above or below the mean ± 2*std.

## Discussion

### The first imputed ancient dog and wolf genomes

Our results show that it is both possible and beneficial to impute ancient dogs and ancient wolves based on present-day canid haplotypes. We can confidently impute data from ancient dogs with coverage as low as 0.5x, and ancient wolves as low as 1x, when applying the appropriate post-imputation MAF (≥0.01) and INFO score (≥0.8) filters. These results were consistent across the major dog lineages tested: Arctic, European, African/Near Eastern and Asian, for which the reference panel contained a sufficient number of present-day representatives such as modern breeds and village dogs.

An imputed North American pre-contact dog (Port au Choix) showed lower accuracy than other dogs. This lineage originated from Siberia and spread into the Americas 20,000 years ago, prior to European colonisation (Ní Leathlobhair et al. 2018; Sinding et al. 2020; Perri et al. 2021). The isolation of this lineage and its near disappearance after the arrival of Europeans means that today this ancestry is not well represented in our panel. Therefore, we suggest that our reference panel does not contain sufficient haplotypes to impute low-coverage North American pre-contact dogs with high accuracy. This may also explain the relatively greater improvement in accuracy observed when restricting our analysis to transversion sites only for this sample, as transition sites, which are affected by aDNA damage, cannot be properly corrected by imputation if there is poor haplotypic representation of the relevant ancestries. Previous studies imputing ancient humans (Sousa Da Mota et al. 2023), horses (Todd et al. 2023) and pigs (Erven et al. 2022) have shown how imputation accuracy can change depending on the ancestral composition of the reference panel. We suggest that ancestry-specific coverage cutoffs may be applied prior to imputation.

The inclusion of non-dog canid haplotype donors in the reference panel substantially improved the imputation accuracy in ancient wolves and ancient dogs, albeit to a lesser extent in dogs. Non-dog canids such as wolves and coyotes may have maintained ancestral variation which was also present in ancient dog lineages, and which has now been depleted from modern dog breeds due to multiple bottlenecks and human artificial selection. Thus, including closely related canid species in the reference panel can assist in imputing sites in other ancient canids such as dogs and wolves. To our knowledge, including high coverage ancient samples in reference panels has not yet been tested. Doing so may improve the accuracy of imputation in cases where that ancestry is poorly represented in the reference panel, however, care must be taken to avoid introducing bias from aDNA damage. Future work benchmarking this approach may provide further opportunities to impute ancestries which have scarce representation in present-day populations.

We note that additional post or pre-imputation filtering approaches as tested in (Hui et al. 2020) could potentially further improve the imputation of samples at lower coverages or with limited ancestral representation in the reference panel. This could be a focus of subsequent studies.

### Imputed vs pseudohaploid genotypes in PCA space

In most cases, the projection of imputed genotypes outperformed the projection of pseudohaploid genotypes in PCA space, particularly for dog genomes above 0.5x and ancient wolf genomes above 1x. Three imputed target ancient dogs (Port au Choix, Newgrange, SOTN01) performed better across all coverages compared to their pseudohaploid versions. These results suggest that imputing diploid genotype data from each sample retains more information and can correct potential biases introduced when calling one allele per site, as done during pseudohaploidisation.

For some samples with lower coverages (<0.2x), pseudohaploidisation surpassed imputation performance (e.g. for Historical Siberia dog, Chinese Village dog, Nigerian Village dog). We attribute this to the larger proportion of sites filtered out in low coverage samples after imputation. In those cases, higher uncertainty is expected at imputed sites with lower depth of coverage and/or among sites with less shared variation with the reference panel, which in turn means that more sites get filtered out when applying the INFO score cutoff. Effectively, this means that the pseudohaploid version of the sample ends up containing more sites than the imputed version, and so tends to be better placed in PCA space.

A notable difference was observed when comparing the placement of the imputed Pleistocene wolves. When applying post-imputation filters, imputation performed worse than pseudohaploidisation across all coverages, whereas applying no post-imputation filters led to better performance for coverages above and equal to 1x. This suggests a trade-off between retaining fewer but more accurate imputed sites, versus retaining more sites with higher uncertainty. Furthermore, the PCA made by present-day genetic variation may not provide the ideal space onto which to project ancestral genetic variation. It has been shown that Pleistocene wolves represent a basal lineage that branched off before the differentiation of present-day wolves and dogs (Ramos-Madrigal et al. 2021; Bergström et al. 2022). Therefore, projecting these ancestries onto a PCA space determined by modern variation may produce misleading placements. All Pleistocene wolves formed a distinct cluster close to present-day East Eurasian Wolves. These Pleistocene wolves are distributed across Eurasia (Germany, Russia, Belgium) and North America (USA and Canada), thus supporting the notion of a panmictic population without strong population structure throughout the Pleistocene (Bergström et al. 2022). The Holocene Eastern Eurasian sample clustered with present-day Eastern Eurasian wolves whereas the Holocene Western Eurasian wolf samples clustered with present-day Western Eurasian wolves (Fig. S55).

Notably, we observed that the PCA placement of the HC imputed samples varied from that of the HC genotyped samples in all dog samples. This may be due to residual genotype errors in the HC genotyped samples which are corrected by imputation, or it may be due to imputation bias from the modern haplotypes in the reference panel.

Overall, the downsampled imputed samples were projected further from the HC genotyped in PCA space compared to the pseudohaploid ones, when the PCs were constructed using genetic variation that is distantly related to the target sample (e.g. Pleistocene vs present-day wolves). For samples which belong to ancestries that are not well represented in the PCA space, we observed better performance when retaining more sites with higher genotype uncertainty (i.e., not applying any post-imputation filters). Considering that imputation corrects for genotyping errors, we suggest that the imputed samples should be incorporated in the making of the PC axis, rather than simply projected onto a pre-existing PCA space, in order to capture the full patterns of genetic variation.

### Accuracy of ROH estimation in imputed samples

The number and length of ROH across the genome can reveal past demographic processes such as recent or past bottlenecks (Palkopoulou et al. 2015; Ceballos et al. 2018, 2021; van der Valk et al. 2019). However, estimating ROH in ancient samples presents challenges, due to low coverage and post-mortem damage resulting in false heterozygous calls. The imputation of ancient samples can correct for genotyping errors and increase the density of diploid genotypes, thus facilitating more accurate ROH estimation.

We retrieved a high overall concordance of ROH segments that overlapped in the validation and imputed target samples. We found two specific scenarios where there were inconsistencies. First, we observed overestimation of ROH segments in the low coverage (<0.2x) samples, likely due to lower imputation accuracy and to fewer SNPs retained following post-imputation filtering, including many heterozygous sites. This leads to an overestimation of ROH throughout the genome, as shown in the high false discovery rates. Secondly, we observed underestimation of ROH segments in the higher coverage samples. We speculate there are two possible explanations for this. First, imputation may be correcting heterozygous sites, which were incorrectly called homozygous in the ground truth sample. Second, sites which were called heterozygous in the imputed samples may have been removed during the initial filtering of the ground truth samples (see ‘Validation dataset filtering,’ Methods section). We note that decreasing false discovery rates (FDR) were observed with increasing coverage across all target samples (Fig. 3c, S35c-S44c).

Our ROH estimates from the imputed samples resulted in higher accuracy scores than those from ROHan based on the non-imputed version of the data. Given that ROHan is intended to detect ROH in ancient genomes with coverage no lower than 7x and with moderate DNA damage levels, such results are not surprising. This highlights the importance of our approach, which now permits the estimation of ROH in ancient dog samples at coverages as low as 0.5x and for wolves as low as 1x, as long as an appropriate imputation reference panel is available.

Echoing our benchmarking results, a lack of ancestral populations in the reference panel that are good representatives of the target samples led to reductions in imputation accuracy, which subsequently led to less accurate inference of ROH segments. This was the case for a North American pre-contact sample (Port au Choix). Even though accuracy for this sample improved somewhat when restricting to transversions, we emphasise that the biological interpretation of ROH based only on transversion sites is unclear and results should be taken with caution.

### Assessing inbreeding levels of dogs and wolves through time

The evolutionary history of dogs has been tightly linked with human movements, leading to founder events, bottlenecks and admixture between populations (Freedman et al. 2014; Witt et al. 2015; Wang et al. 2016; Botigué et al. 2017; Ní Leathlobhair et al. 2018; Ollivier et al. 2018; Da Silva Coelho et al. 2021; Feuerborn et al. 2021). This, in combination with intensive human driven selective breeding to develop and maintain specific breed traits in more recent periods, have shaped dog genetic diversity through time. Previous studies on wolves have also found past and recent demographic events, including bottlenecks and within and between-species admixture (Pilot et al. 2014, 2019, 2021; Fan et al. 2016; Loog et al. 2020; Bergström et al. 2022; Lobo et al. 2023). Even though wolves have not undergone the same human selective breeding as dogs, they have been subjected to a high degree of human induced pressures via habitat loss and systematic persecution (Wayne et al. 1992; Fredrickson et al. 2007; Sastre et al. 2011; Pilot et al. 2014; Kuijper et al. 2016). Subsequently, inbreeding levels in dog and wolf populations may have changed through time as the result of different factors.

We assessed inbreeding patterns in ancient dogs and ancient wolves using phased and imputed genomes. We found that inbreeding in dogs has predominantly occurred in recent times, with modern breeds containing significantly more ROH than ancient individuals across Eurasia. Despite the overall low levels of inbreeding in ancient samples, some individuals showed relatively high inbreeding coefficients with no clear temporal pattern. These include two Neolithic dogs from Croatia and the island of Gotland in Sweden (∼4,800 BP), a Mesolithic dog from the Veretye site in North-Western Siberia (∼11,000 BP), a Mesolithic dog from Zhokhov island in North-Eastern Siberia (9,515 BP), a 1,111 BP dog and a ∼100 BP dog from the Iamal-Nenets region in North-Western Siberia, a 100 BP dog from the Bulgunnyakhtakh site in North-Eastern Siberia, and a ∼150 BP dog from Lithuania. We hypothesise that, in some of these cases, isolation by distance may be a main driver of these sporadic increases in inbreeding, as the locations of most of these samples seem to be in remote and inaccessible areas, such as the island of Gotland and Northern Siberia. The Lithuanian sample from the 19th century (KT0056), characterised by high levels of inbreeding, aligns with historical records that describe how noble owners of estates engaged in the selective breeding of unique hunting dog varieties. Frequently, these dogs were named after their noble breeders (e.g. Bialozar pointer, Kociol hound) (Dmitrij, V. 1876).

Differences were also observed between present-day populations. Ancient European and Arctic dogs did not differ significantly in estimated ROH levels through most of their history. However present-day European breeds display significantly higher inbreeding levels compared to Near Eastern breeds, likely due to specific and targeted breeding practices since the Victorian era, which led to the formation of European breeds.

High inbreeding levels were also observed for some present-day wolf populations, likely reflecting bottlenecks related to habitat fragmentation and recent population declines (Dufresnes et al. 2018; Kardos et al. 2018; Robinson et al. 2019). The generally low levels of ROH observed in ancient wolves confirms previous findings supporting high connectivity and low differentiation of wolf populations throughout the Pleistocene (Bergström et al. 2022). In their paper, (Bergström et al. 2022) found that despite the higher levels of differentiation in samples from the last 10,000 years, suggestive of population bottlenecks due to habitat fragmentation and human hunting, levels of individual heterozygosity remained the same. They attributed this to limited gene flow rather than a species-wide population decline. This would match our results, with low F*_ROH_* estimates maintained in the Holocene wolves.

Finally, our assessment of ROH frequency in present-day and ancient dog samples showed an enrichment for genes related to olfaction and immunity. Intriguingly, both of these functions are deemed crucial for dogs, which strongly depend on their sense of smell for survival (Miklósi 2014; Serpell 2016) and which have been subject to multiple pathogenic pressures throughout their history of cohabitation and migrations with humans (Liu et al. 2018; Ní Leathlobhair et al. 2018). This ROH pattern may be due to balancing selection for multiple variants associated with these functions, or to the presence of recessive deleterious variation within these regions, leading homozygous individuals to be at a disadvantage. It is possible that some of these signals may be driven by copy number variation, though we attempted to correct for this using strict coverage cutoffs and a gene length correction in the GO enrichment test. Further investigation into these regions via formal tests for selection, or via detailed functional characterisation of the variants within them may shed light on the causes for these patterns.

Imputed diploid genotypes of ancient samples can grant access to genomic tools mainly tailored for analysing high-quality genomic data. This, in turn, can enable researchers to address problems that often require high-quality phased haplotypes, such as detecting natural selection, inferring and dating past admixture events and estimating local ancestry tracts. The increase of sequenced ancient samples will inevitably fill spatiotemporal gaps in the evolutionary history of multiple species, including dogs and wolves. Testing and applying imputation methods on ancient genomes of sequenced species is a promising approach to maximise the genomic information retrieved from each sample, so as to better understand the evolutionary processes that shaped their past and present diversity.

## Data availability

Raw reads generated for this study have been deposited to the European Nucleotide Archive (ENA) under project number PRJEB73844. The code for all the analyses presented is available at https://github.com/katiabou/ancient_dog_imputation_paper. The imputation pipeline is available at https://github.com/katiabou/dog_imputation_pipeline.

## Acknowledgements

We are grateful to the members of the Racimo group for the useful discussions throughout the different parts of this study. F.R. and K.B. were supported by a Villum Young Investigator Grant (project no. 00025300). F.R. was also supported by a Novo Nordisk Fonden Data Science Ascending Investigator Award (NNF22OC0076816) and by the European Research Council (ERC) under the European Union’s Horizon Europe programme (grant agreements No. 101077592 and 951385). L.A.F..F. and G.L. were supported by European Research Council grants (ERC-2013-StG-337574-UNDEAD and ERC2019-StG-853272-PALAEOFARM) and Natural Environment Research Council grants (NE/K005243/1, NE/K003259/1, NE/S007067/1, and NE/S00078X/1). L.A.F.F. and A.C. were supported by the Wellcome Trust (210119/Z/18/Z). G.P. and P.B. were supported by a Research Council of Lithuania, grant number S-MIP-20-5. S.G was supported by a Danish National Research Foundation award - DNRF143. E.A.O. was supported by the Intramural Program of the National Human Genome Research Institute.

## Author contributions

K.B., E.K.I.P., L.A.F.F. and F.R. led the study. K.B., E.K.I.P., L.A.F.F. and F.R. conceptualised the study. E.K.I.P., L.A.F.F., E.A.O. and F.R. supervised the research. L.A.F.F., G.L., E.A.O. and F.R. acquired funding for research. G.P. and P.B. were involved in sample collection. S.C., T.R.F., S.G., A.H., E.A.O., H.G.P., G.P., L.S. and P.B. were involved in sample curation. S.C. and K.T. undertook laboratory work. K.B., S.G.A., A.C. and A.H. undertook formal analysis of the data. K.B., L.A.F.F., E.K.I.P. and F.R. drafted the main text. K.B., S.G.A., S.C., L.A.F.F., A.H. and G.P. drafted supplementary notes and materials. K.B., S.C., L.A.F.F., E.K.I.P., G.L., E.A.O., F.R. and L.S. were involved in reviewing drafts and editing.

