## Supplementary figures for "Imputation of ancient canid genomes reveals inbreeding history over the past 10,000 years"

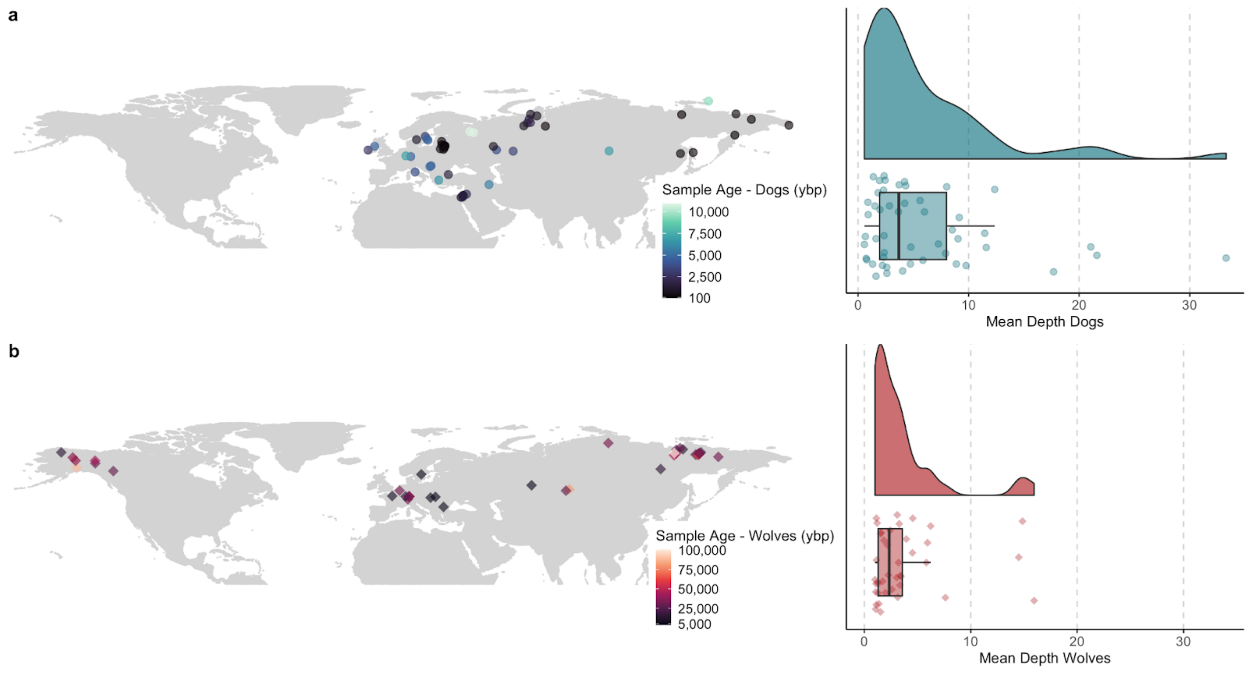


**Fig. S1:** *Map of ancient a) dog (>0.5x) and b) wolf samples (>1x) used in this study, along with the distributions of mean depth for each.*

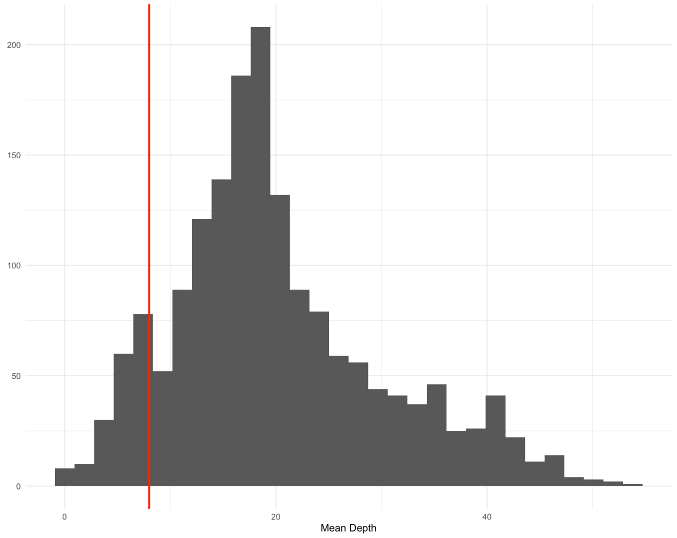


**Fig. S2:** *Histogram showing the mean depth for all genomes within the 1,715 sample VCF. The red line indicates the 8x cutoff we applied for including a sample in the reference panel. Two samples with very high coverage (>300x) were excluded from the plot for visualisation purposes.*

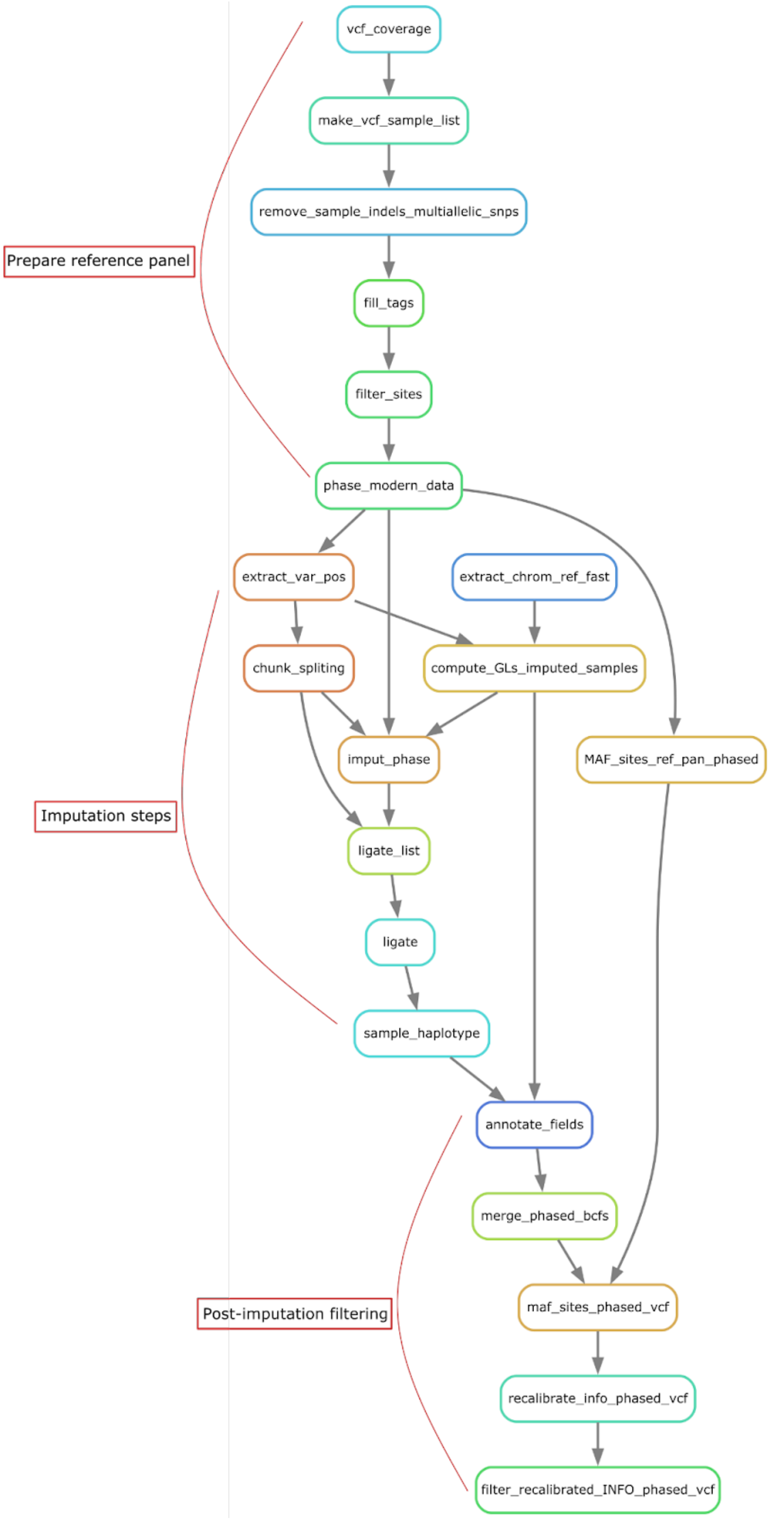

Prepare reference panel
vcf_coverage
make_vcf_sample_list
remove_sample_indels_multiallelic_snps
fill_tags
filter_sites
phase_modern_data
extract_var_pos
extract_chrom_ref_fast
chunk_spliting
compute_GLs_imputed_samples
imput_phase
MAF_sites_ref_pan_phased
Imputation steps
ligate_list
ligate
sample_haplotype
annotate_fields
merge_phased_bcfs
Post-imputation filtering
maf_sites_phased_vcf
recalibrate_info_phased_vcf
filter_recalibrated_INFO_phased_vcf

**Fig. S3:** *Snakemake rulegraph of the imputation pipeline used for this study partitioned into three sections: 1) Filtering and phasing of the reference panel, 2) imputation and phasing of ancient samples and 3) post imputation filtering by applying MAF and INFO score cutoffs.*

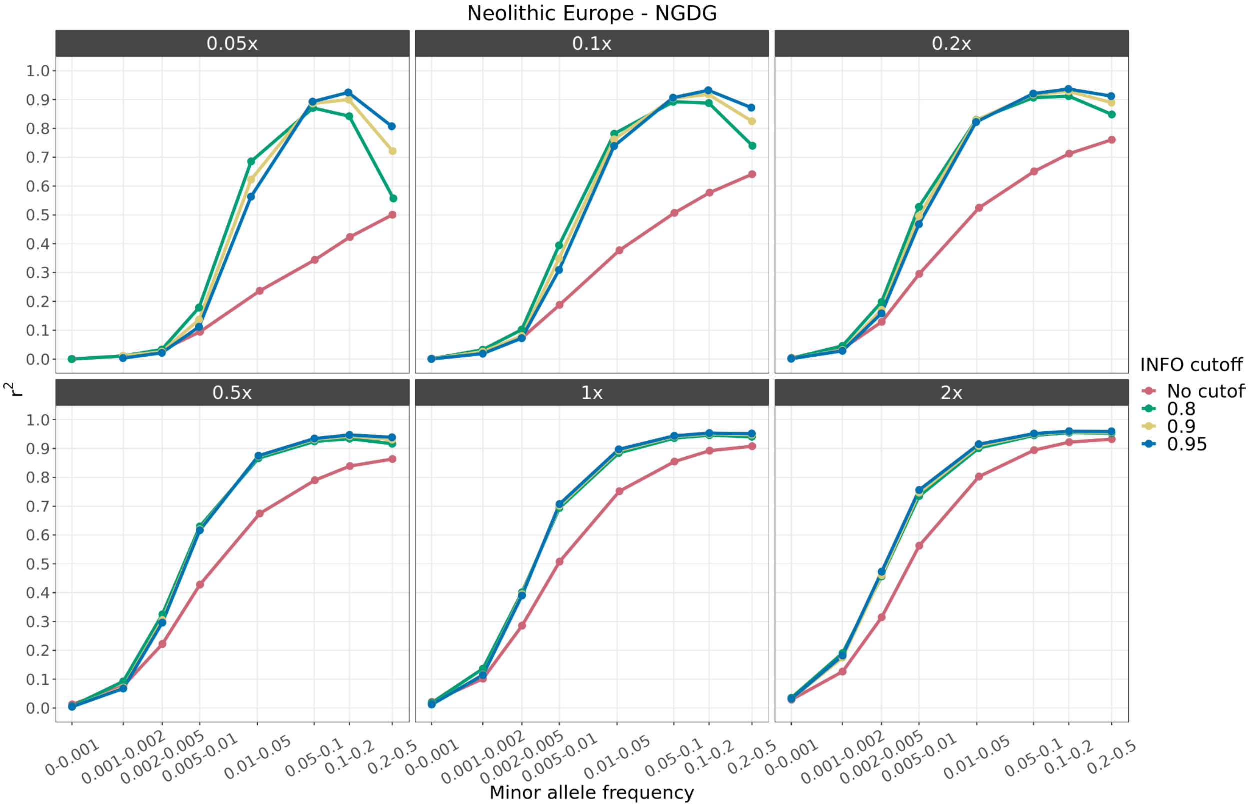


**Fig. S4:** *Squared correlation between imputed genotypes by GLIMPSE and highly confident called genotypes for the Newgrange Neolithic European dog, downsampled to six coverage values (0.01x, 0.05, 0.1x, 0.5x, 1x and 2x) and across different MAF bins. All target samples were imputed using the reference panel containing all canids. Each colour depicts the accuracy for a given INFO score cutoff. Red: no cut-off, Green: 0.8, Yellow: 0.9 and Blue: 0.95.*

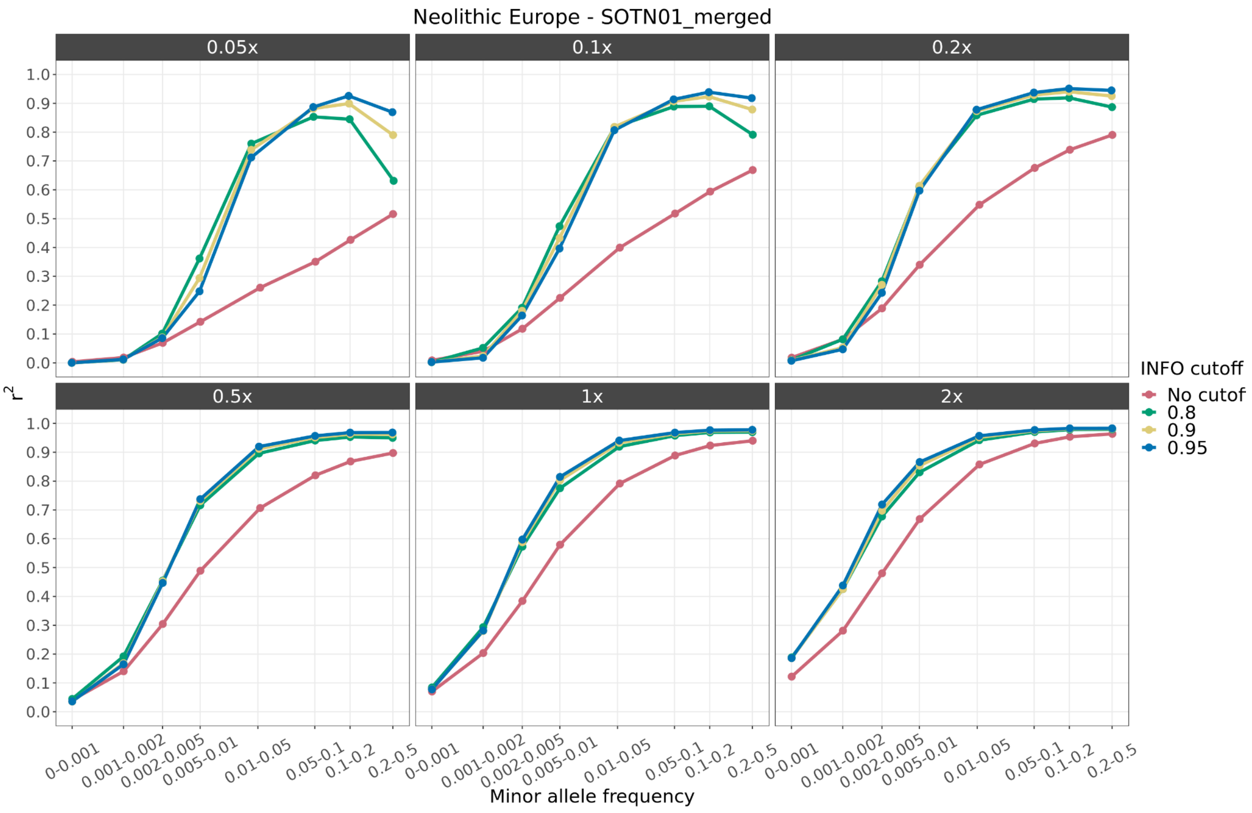


**Fig. S5:** *Squared correlation between imputed genotypes by GLIMPSE and highly confident called genotypes for the SOTN01 Neolithic European dog, downsampled to six coverage values (0.01x, 0.05, 0.1x, 0.5x, 1x and 2x) and across different MAF bins. All target samples were imputed using the reference panel containing all canids. Each colour depicts the accuracy for a given INFO score cutoff. Red: no cut-off, Green: 0.8, Yellow: 0.9 and Blue: 0.95.*

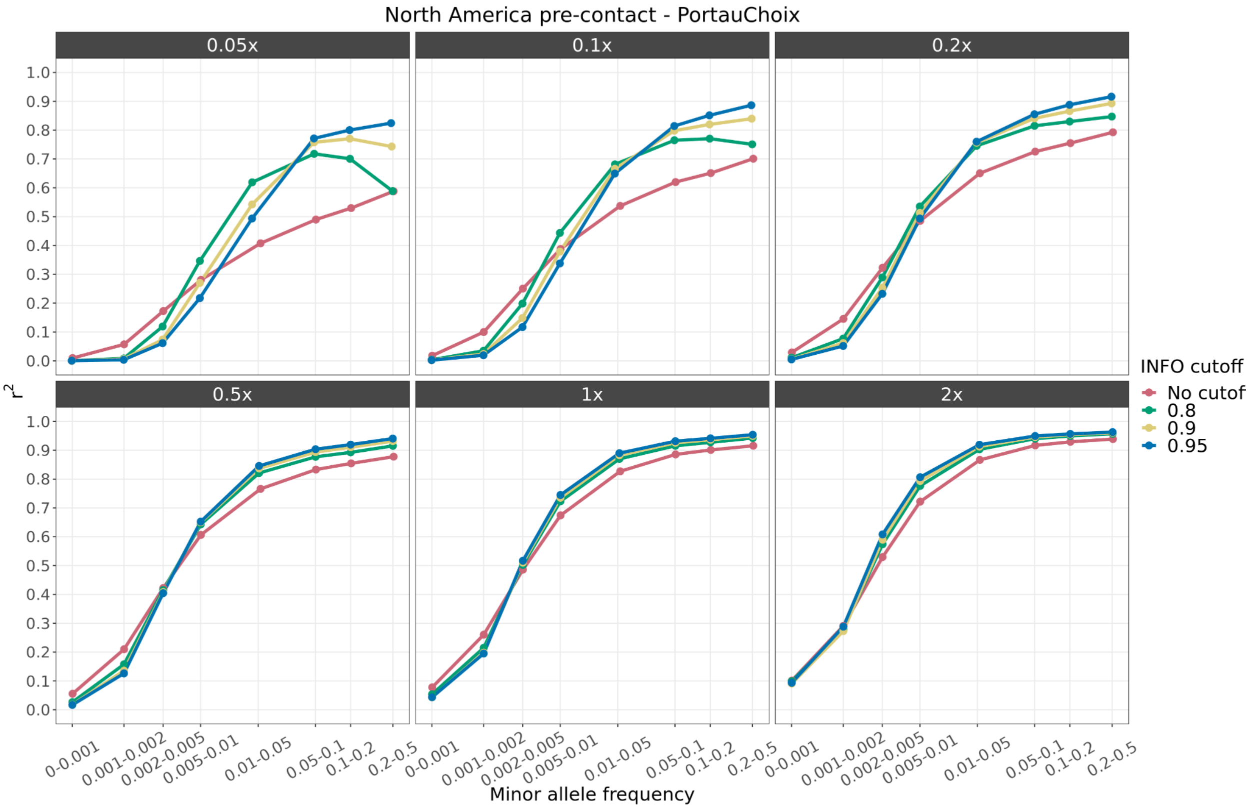


**Fig. S6:** *Squared correlation between imputed genotypes by GLIMPSE and highly confident called genotypes for the Port au Choix North American pre-contact dog, downsampled to six coverage values (0.01x, 0.05, 0.1x, 0.5x, 1x and 2x) and across different MAF bins. All target samples were imputed using the reference panel containing all canids. Each colour depicts the accuracy for a given INFO score cutoff. Red: no cut-off, Green: 0.8, Yellow: 0.9 and Blue: 0.95.*

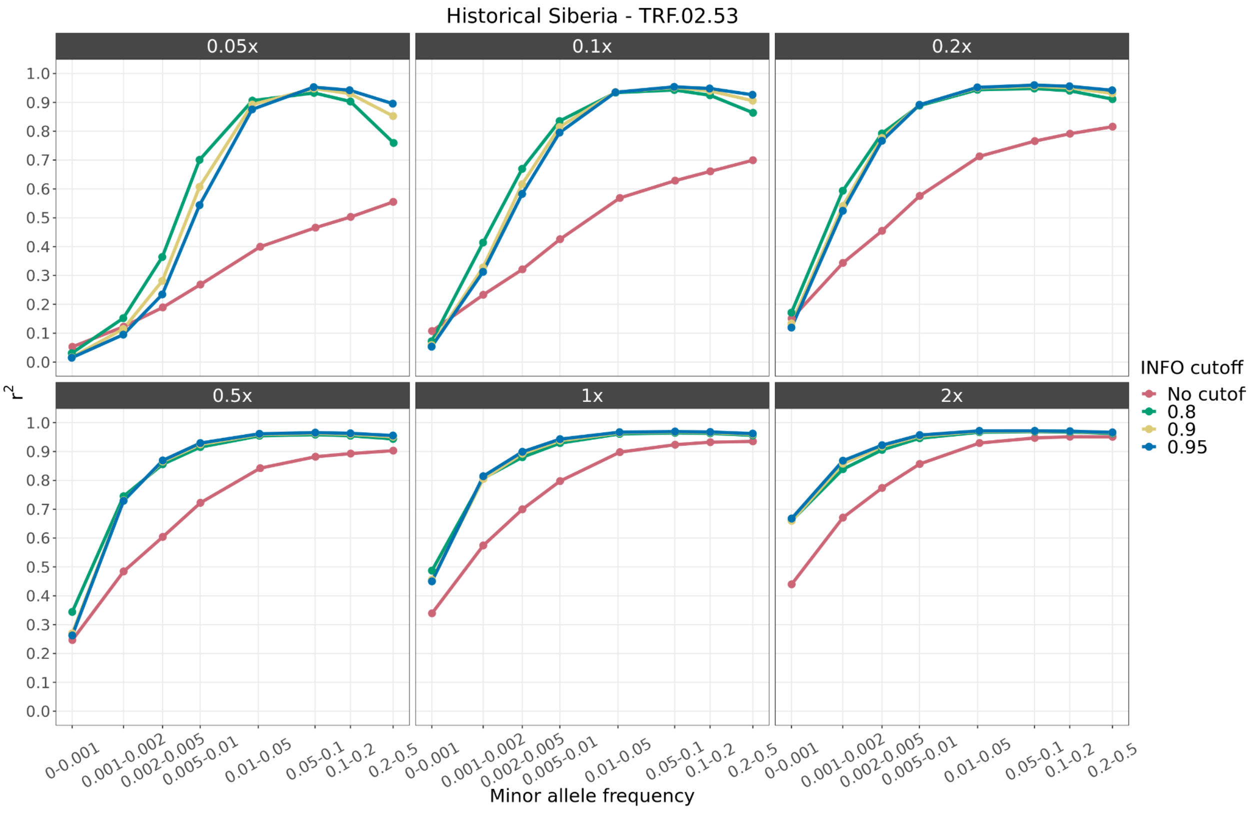


**Fig. S7:** *Squared correlation between imputed genotypes by GLIMPSE and highly confident called genotypes for the TRF.02.53 historical Siberian dog, downsampled to six coverage values (0.01x, 0.05, 0.1x, 0.5x, 1x and 2x) and across different MAF bins. All target samples were imputed using the reference panel containing all canids. Each colour depicts the accuracy for a given INFO score cutoff. Red: no cut-off, Green: 0.8, Yellow: 0.9 and Blue: 0.95.*

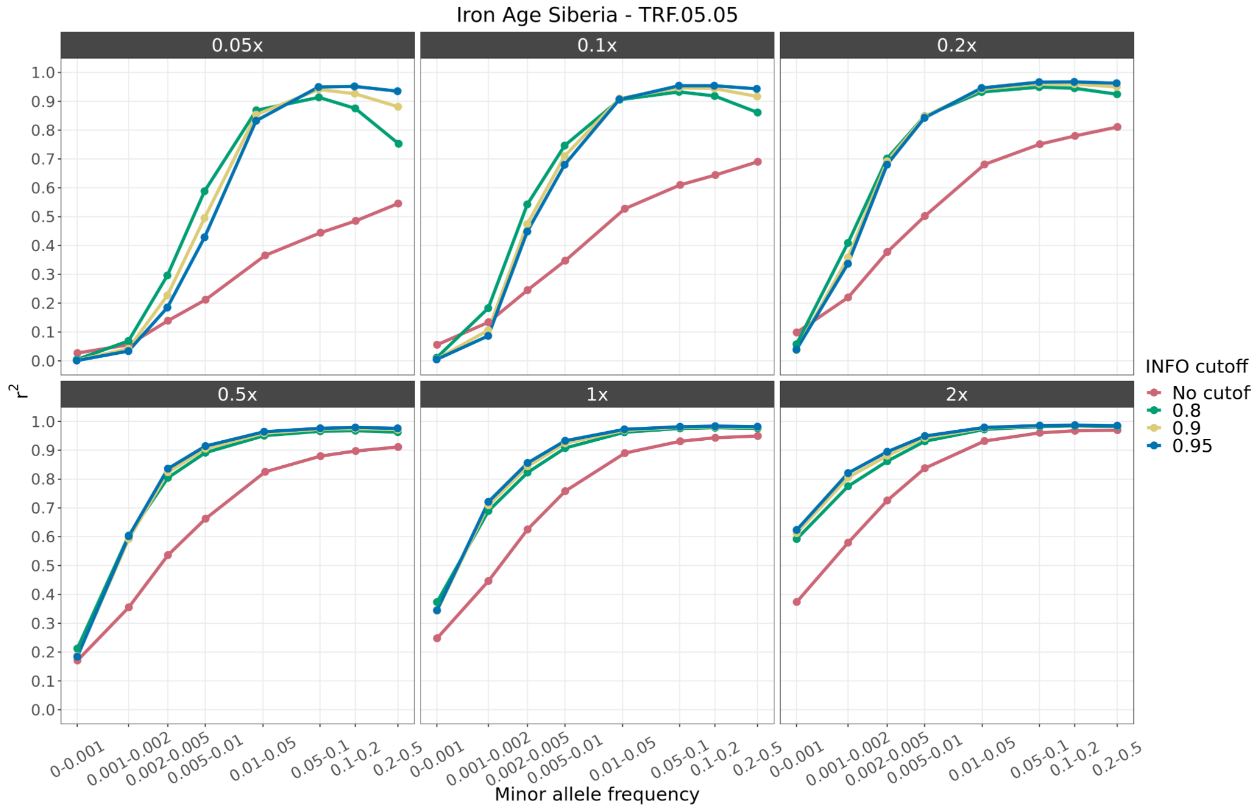


**Fig. S8:** *Squared correlation between imputed genotypes by GLIMPSE and highly confident called genotypes for the TRF.05.05 Iron Age Siberian dog, downsampled to six coverage values (0.01x, 0.05, 0.1x, 0.5x, 1x and 2x) and across different MAF bins. All target samples were imputed using the reference panel containing all canids. Each colour depicts the accuracy for a given INFO score cutoff. Red: no cut-off, Green: 0.8, Yellow: 0.9 and Blue: 0.95.*

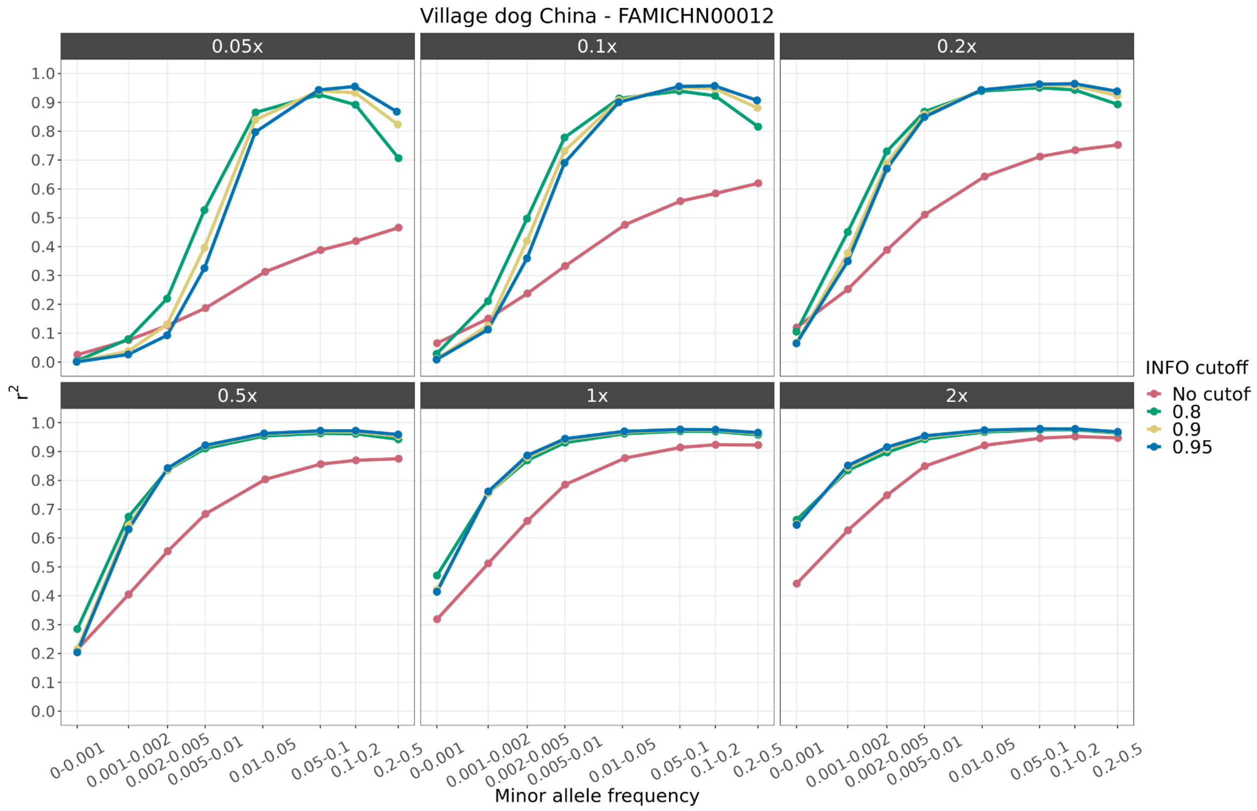


**Fig. S9:** *Squared correlation between imputed genotypes by GLIMPSE and highly confident called genotypes for the FAMICHN00012 Chinese Village dog, downsampled to six coverage values (0.01x, 0.05, 0.1x, 0.5x, 1x and 2x) and across different MAF bins. All target samples were imputed using the reference panel containing all canids. Each colour depicts the accuracy for a given INFO score cutoff. Red: no cut-off, Green: 0.8, Yellow: 0.9 and Blue: 0.95.*

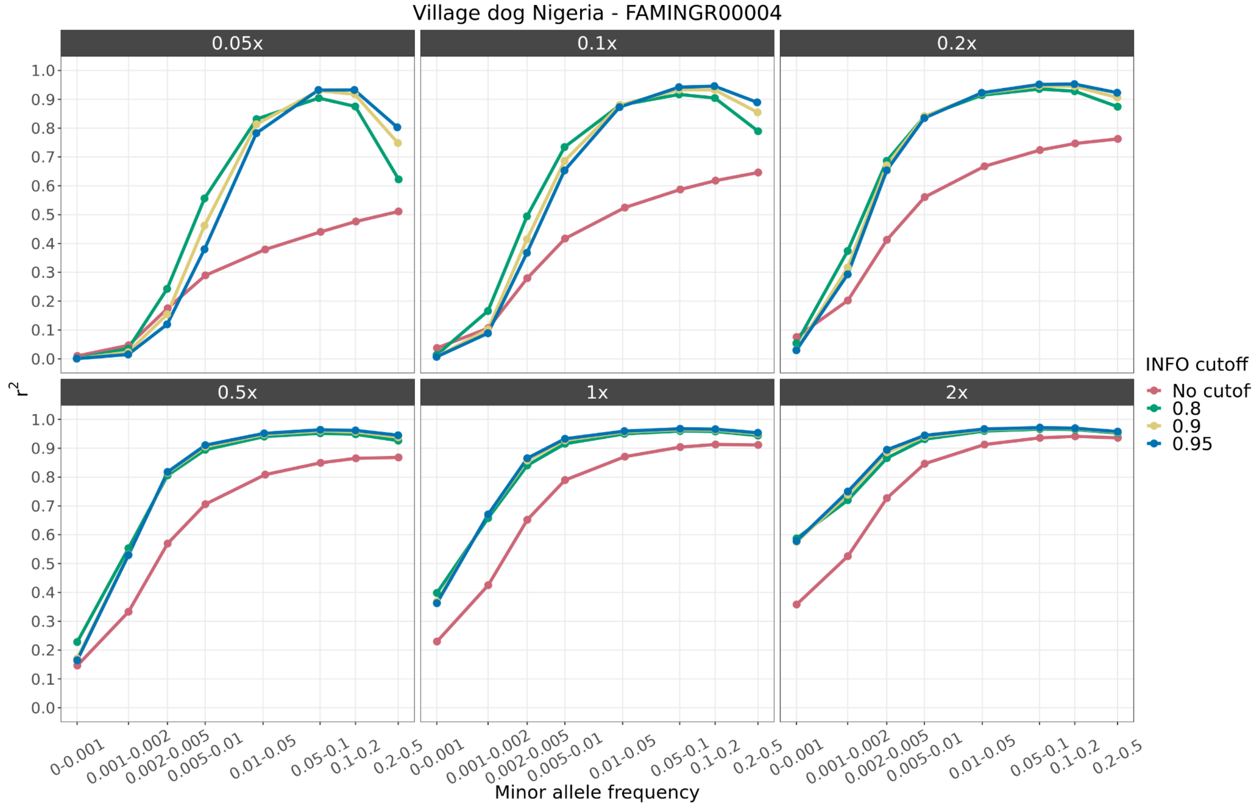


**Fig. S10:** *Squared correlation between imputed genotypes by GLIMPSE and highly confident called genotypes for the FAMINGR00004 Nigerian Village dog, downsampled to six coverage values (0.01x, 0.05, 0.1x, 0.5x, 1x and 2x) and across different MAF bins. All target samples were imputed using the reference panel containing all canids. Each colour depicts the accuracy for a given INFO score cutoff. Red: no cut-off, Green: 0.8, Yellow: 0.9 and Blue: 0.95.*

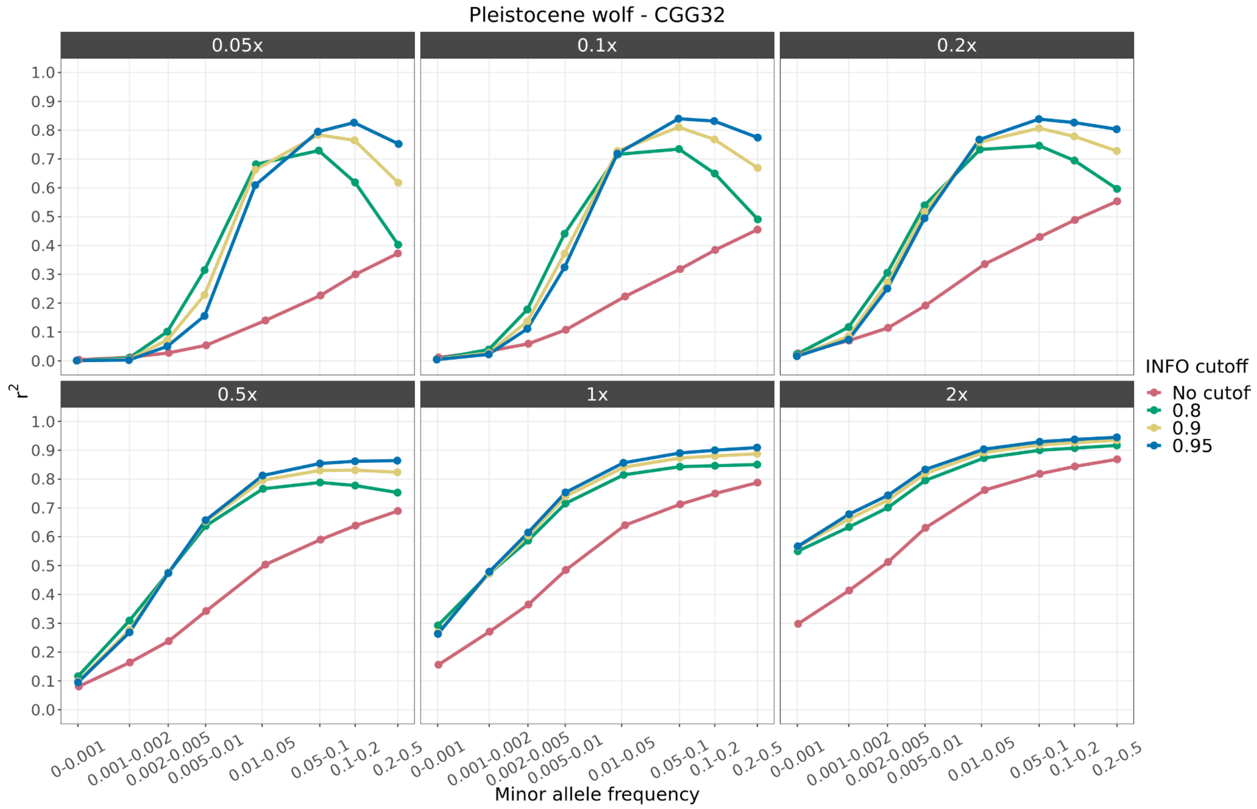


**Fig. S11:** *Squared correlation between imputed genotypes by GLIMPSE and highly confident called genotypes for the CGG32 Pleistocene wolf, downsampled to six coverage values (0.01x, 0.05, 0.1x, 0.5x, 1x and 2x) and across different MAF bins. All target samples were imputed using the reference panel containing all canids. Each colour depicts the accuracy for a given INFO score cutoff. Red: no cut-off, Green: 0.8, Yellow: 0.9 and Blue: 0.95.*

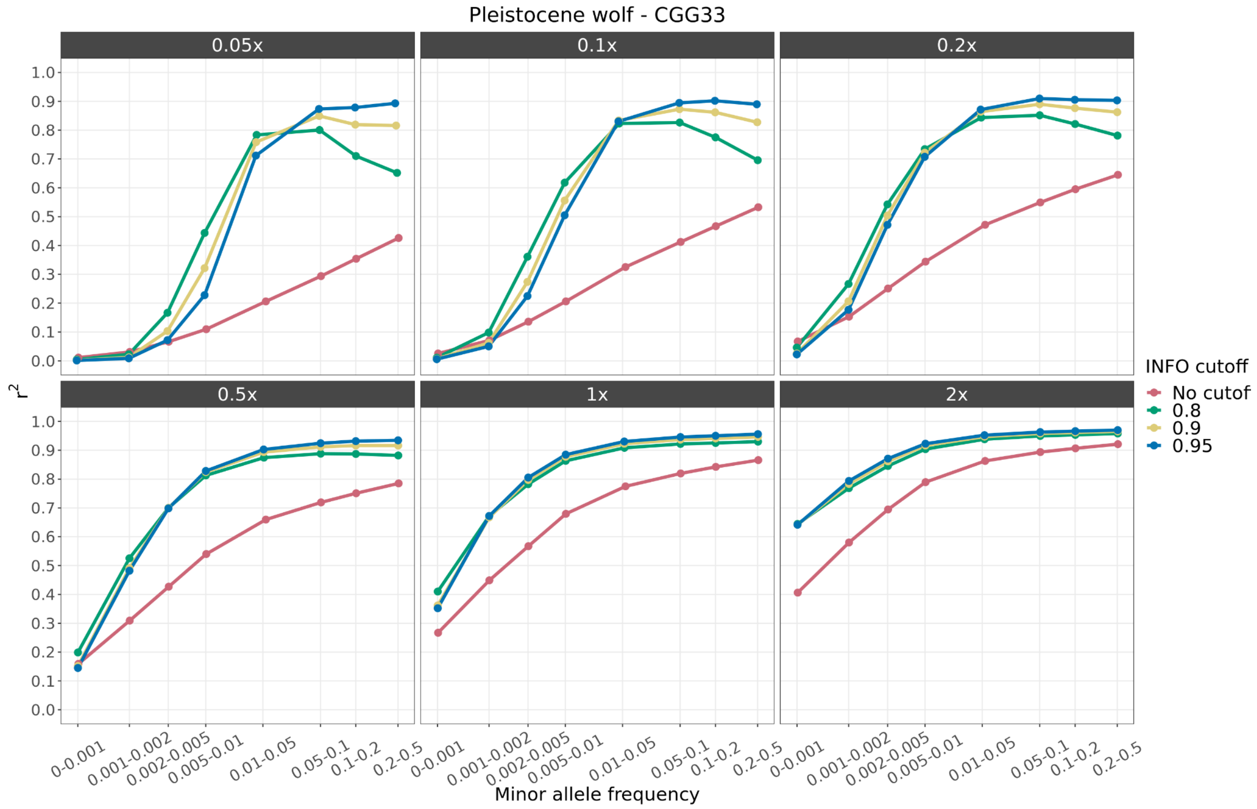


**Fig. S12:** *Squared correlation between imputed genotypes by GLIMPSE and highly confident called genotypes for the CGG33 Pleistocene wolf, downsampled to six coverage values (0.01x, 0.05, 0.1x, 0.5x, 1x and 2x) and across different MAF bins. All target samples were imputed using the reference panel containing all canids. Each colour depicts the accuracy for a given INFO score cutoff. Red: no cut-off, Green: 0.8, Yellow: 0.9 and Blue: 0.95.*

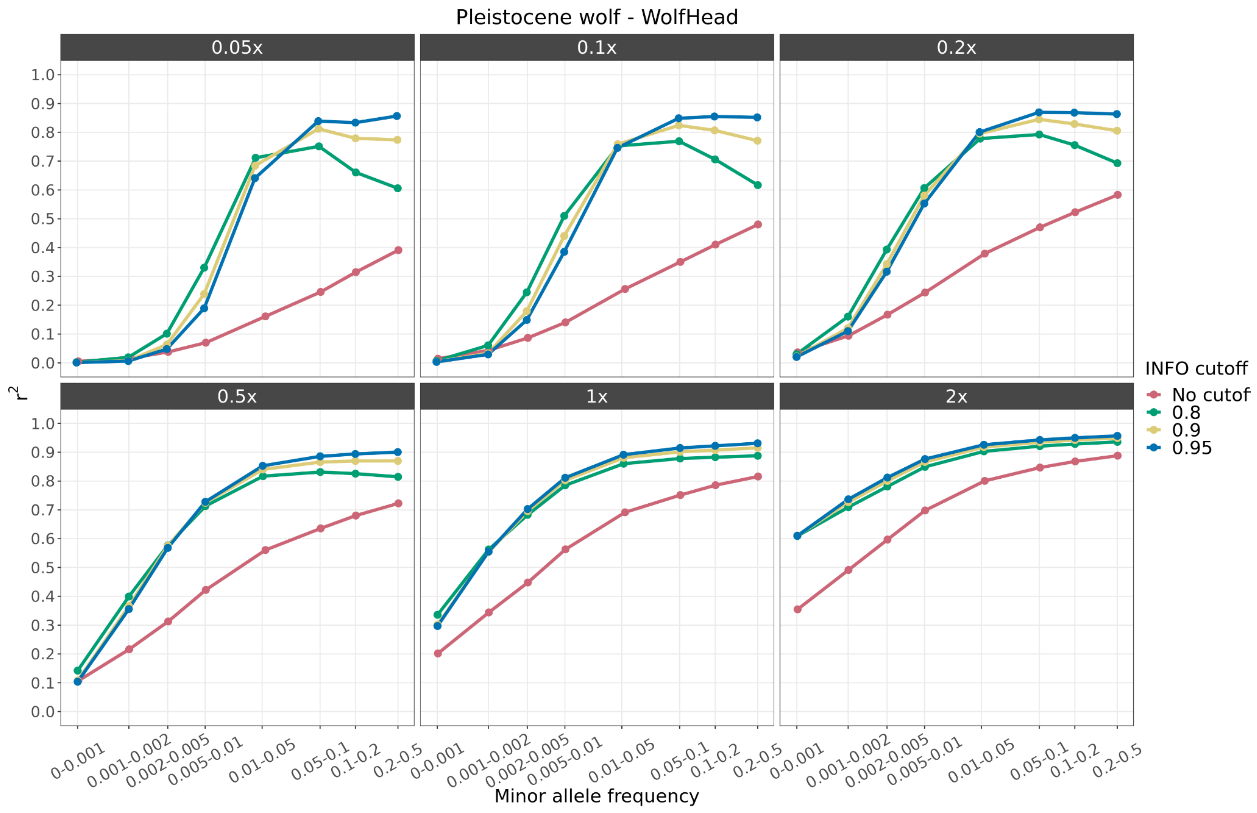


**Fig. S13:** *Squared correlation between imputed genotypes by GLIMPSE and highly confident called genotypes for the WolfHead Pleistocene wolf, downsampled to six coverage values (0.01x, 0.05, 0.1x, 0.5x, 1x and 2x) and across different MAF bins. All target samples were imputed using the reference panel containing all canids. Each colour depicts the accuracy for a given INFO score cutoff. Red: no cut-off, Green: 0.8, Yellow: 0.9 and Blue: 0.95.*

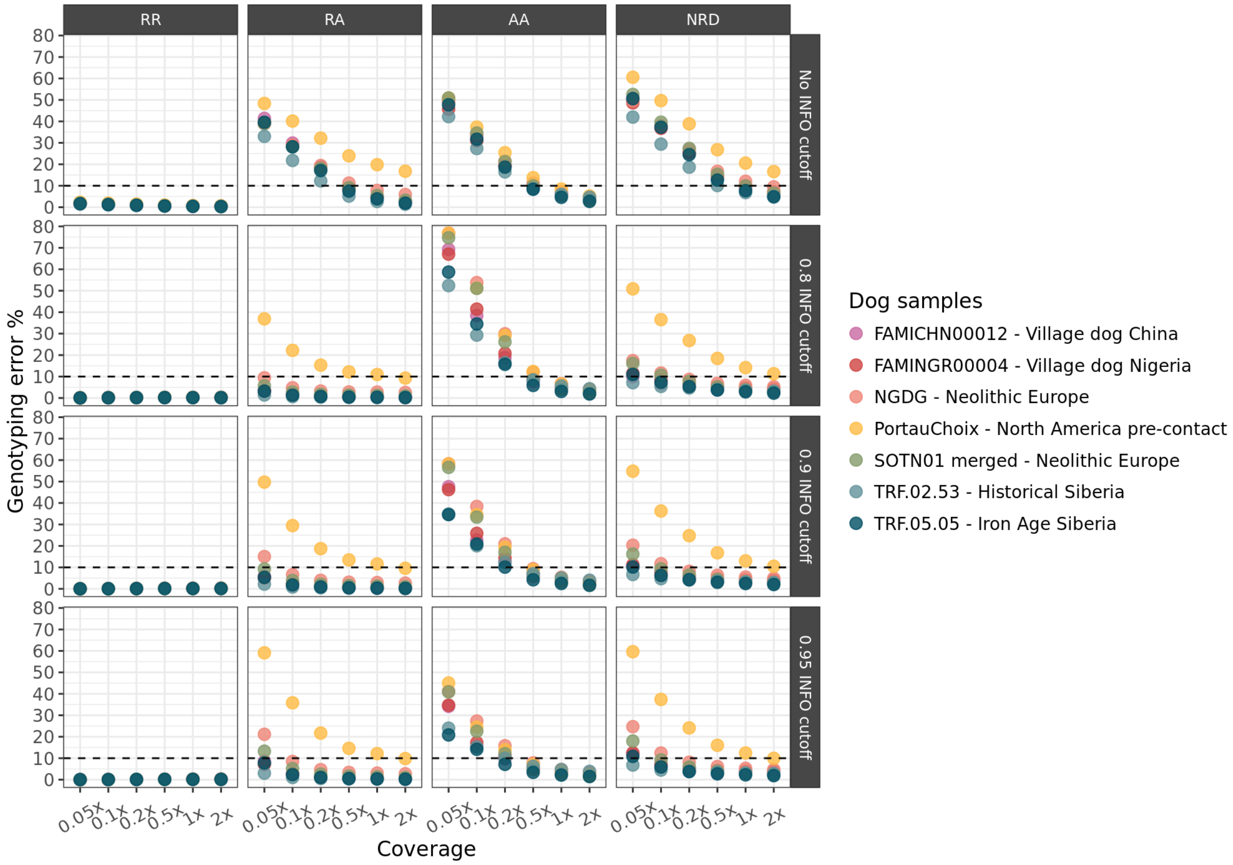


**Fig. S14:** *Genotyping error between imputed downsampled and the high coverage target dog samples for homozygous alternative (AA), heterozygous (RA) and homozygous reference alleles (RR), and the non-reference discordance (NRD) metric estimated by GLIMPSE concordance for all autosomes. Comparisons are shown for different INFO score cutoffs (no cutoff, 0.8, 0.9 and 0.95).*

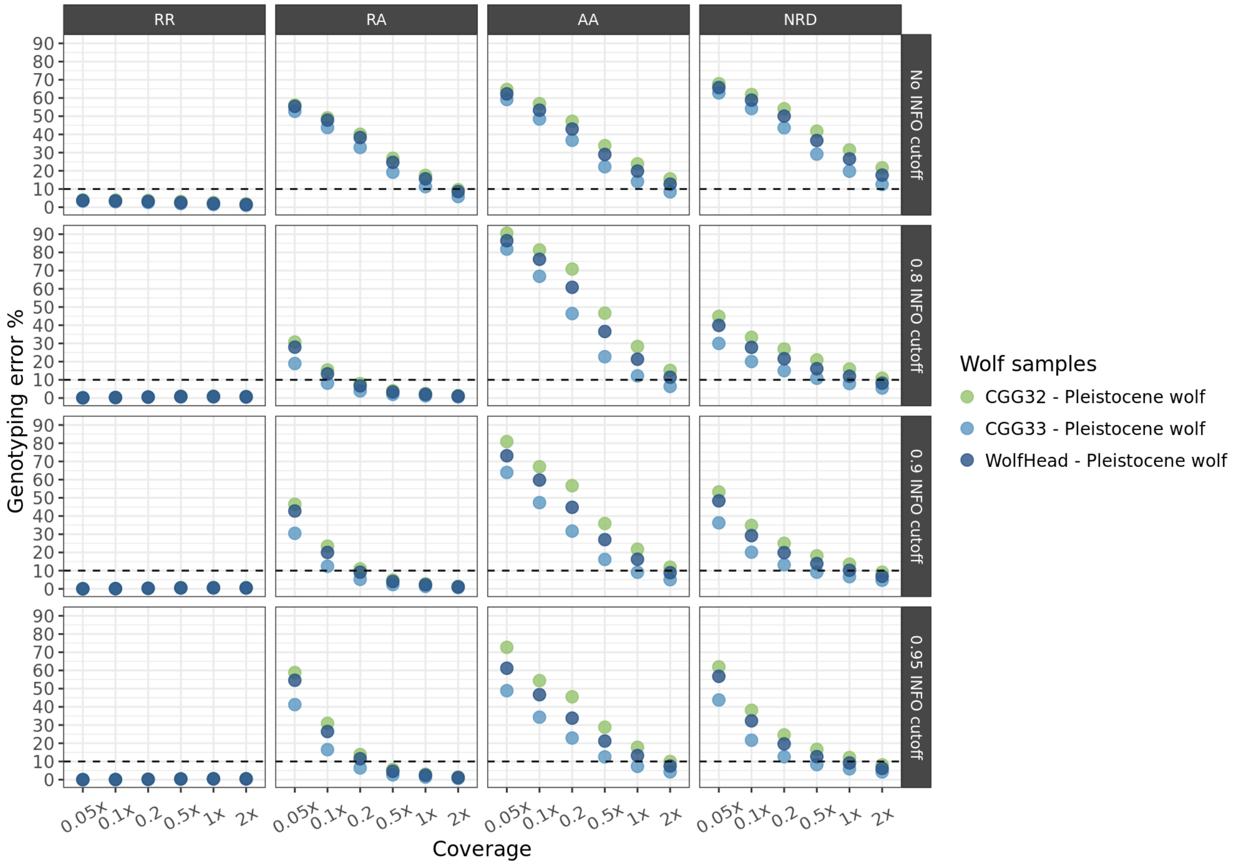


**Fig. S15:** *Genotyping error between imputed downsampled and the high coverage target Pleistocene wolf samples for homozygous alternative (AA), heterozygous (RA) and homozygous reference alleles (RR), and the non-reference discordance (NRD) metric estimated by GLIMPSE concordance for all autosomes. Comparisons are shown for different INFO score cutoffs (no cutoff, 0.8, 0.9 and 0.95).*

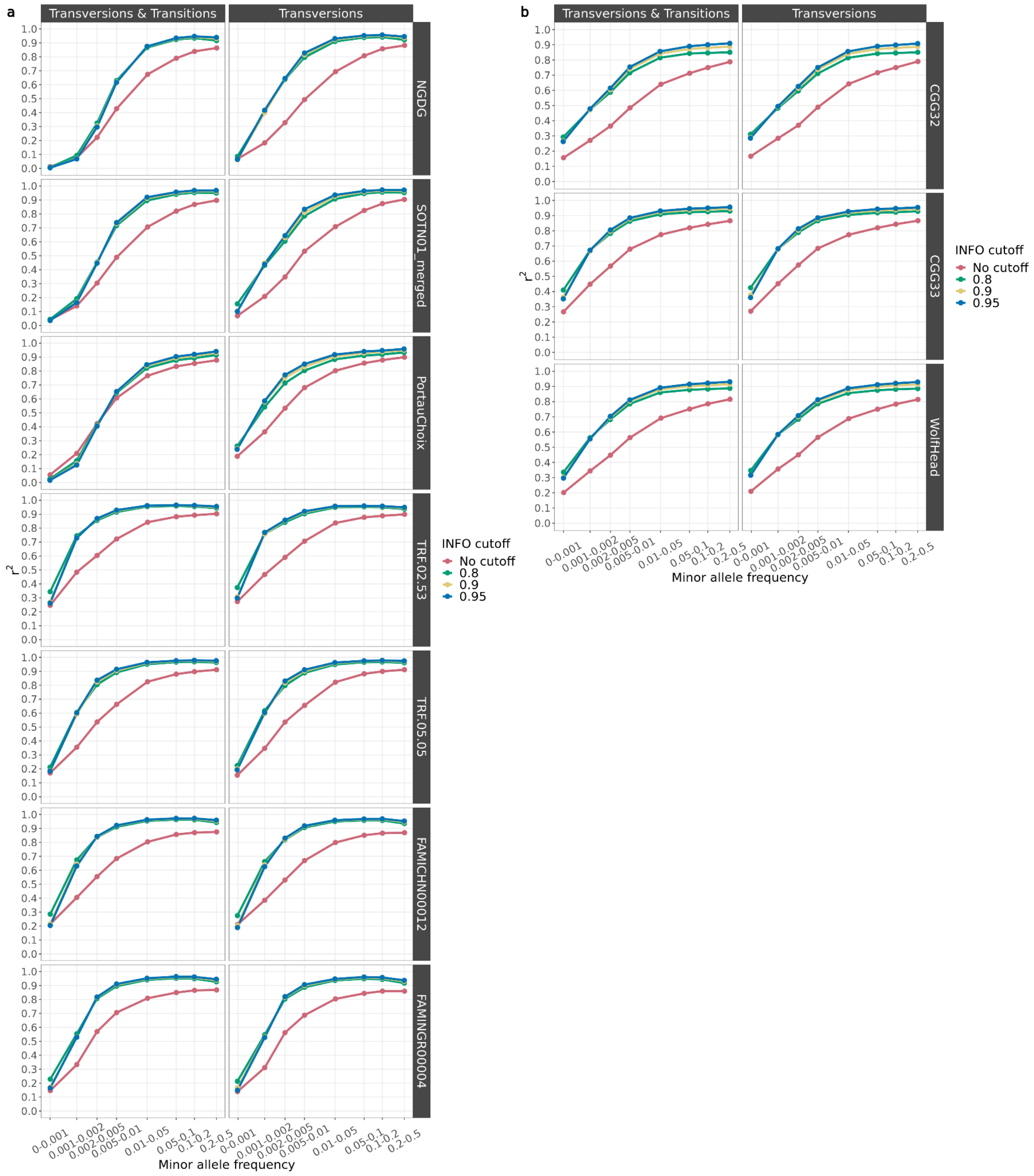


**Fig. S16:** *Squared correlation between imputed genotypes by GLIMPSE and highly confident called genotypes for all 10 target samples, downsampled to a) 0.5x for dogs and b) 1x for wolves and across different MAF bins using either transversions and transitions or only transversions. Each colour depicts the accuracy for a given INFO score cutoff. Red: no cut-off, Green: 0.8, Yellow: 0.9 and Blue: 0.95.*

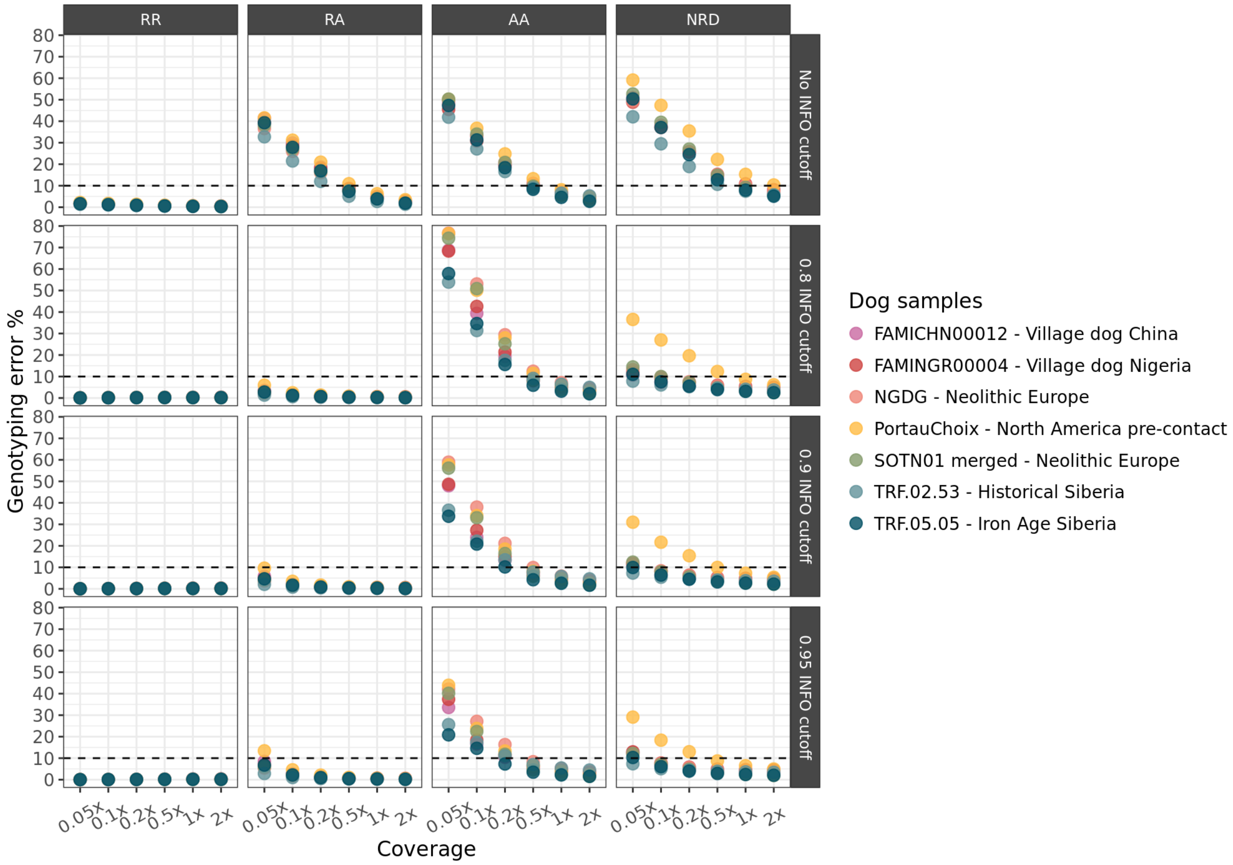


**Fig. S17:** *Genotyping error between imputed downsampled and the high coverage target dog samples using only transversions for homozygous alternative (AA), heterozygous (RA) and homozygous reference alleles (RR), and the non-reference discordance (NRD) metric estimated by GLIMPSE concordance for all autosomes. Comparisons are shown for different INFO score cutoffs (no cutoff, 0.8, 0.9 and 0.95).*

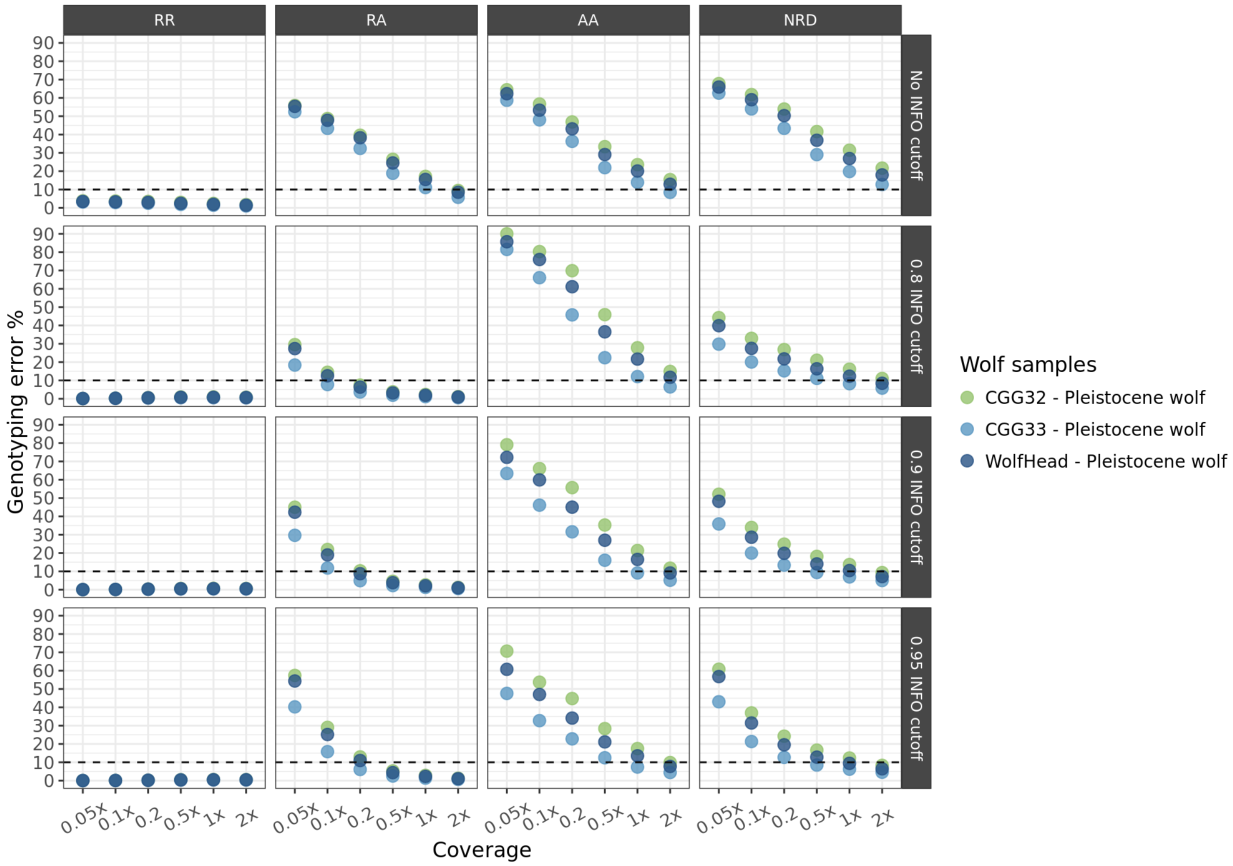


**Fig. S18:** *Genotyping error between imputed downsampled and the high coverage target Pleistocene wolf samples using only transversions for homozygous alternative (AA), heterozygous (RA) and homozygous reference alleles (RR), and the non-reference discordance (NRD) metric estimated by GLIMPSE concordance for all autosomes. Comparisons are shown for different INFO score cutoffs (no cutoff, 0.8, 0.9 and 0.95).*

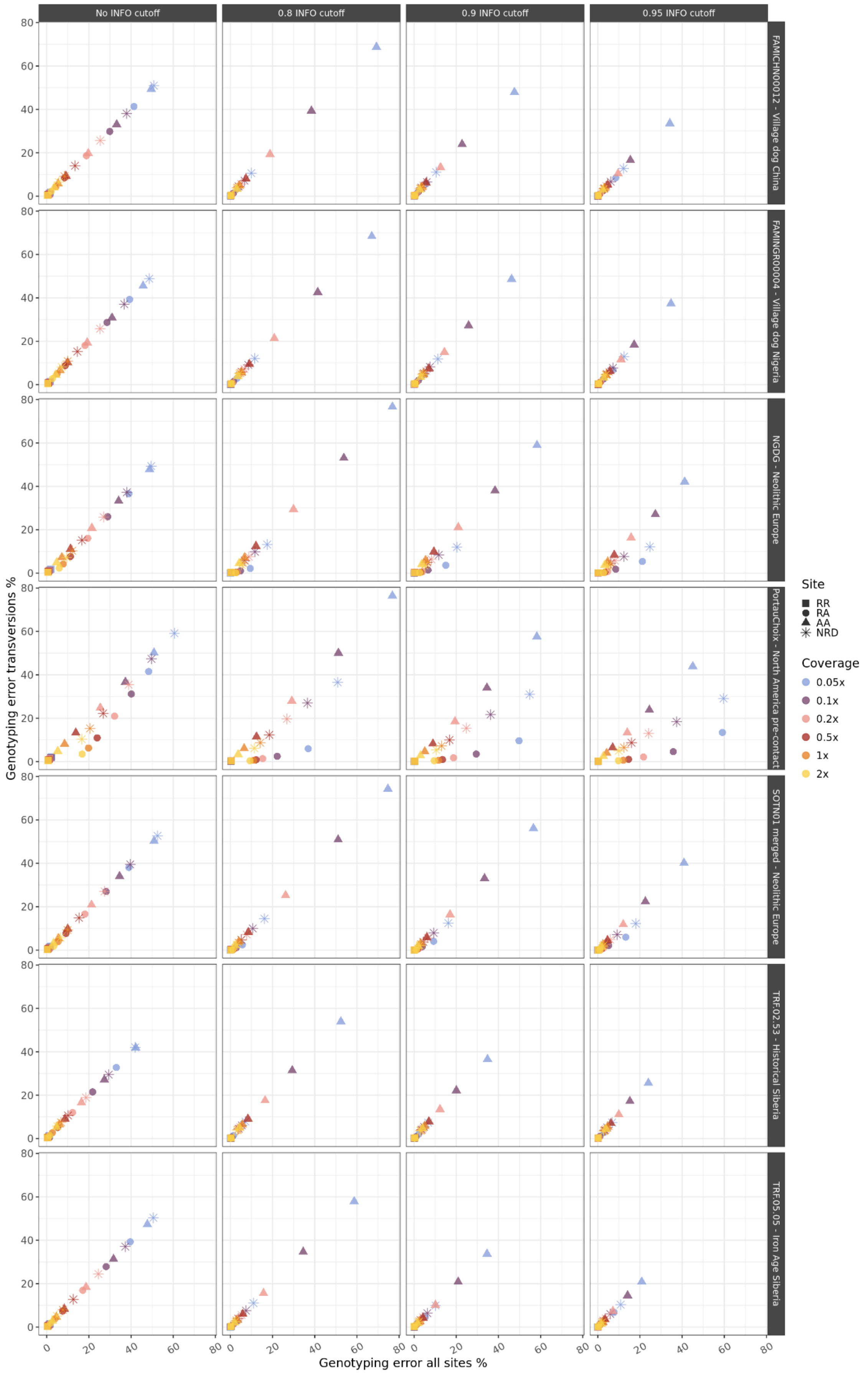
No INFO cutoff
0.8 INFO cutoff
0.9 INFO cutoff
0.95 INFO cutoff
FAMICHN00012 - Village dog China
FAMINGR00004 - Village dog Nigeria
NGDG - Neolithic Europe
PortauChoix - North America pre-contact
SOTN01 merged - Neolithic Europe
TRF.02.53 - Historical Siberia
TRF.05.05 - Iron Age Siberia
Genotyping error transversions %
Genotyping error all sites %
Site
RR
RA
AA
NRD
Coverage
0.05x
0.1x
0.2x
0.5x
1x
2x

**Fig. S19:** *Genotyping error of transversions and transitions (all sites) against genotyping error of transversions for the seven imputed target dog samples (rows) applying different INFO score cutoffs (columns). Estimates are shown for genotyping errors of homozygous reference (RR), heterozygous (RA), homozygous alternative (AA) alleles and the non-reference discordance (NRD) metric. Colours represent different coverages.*

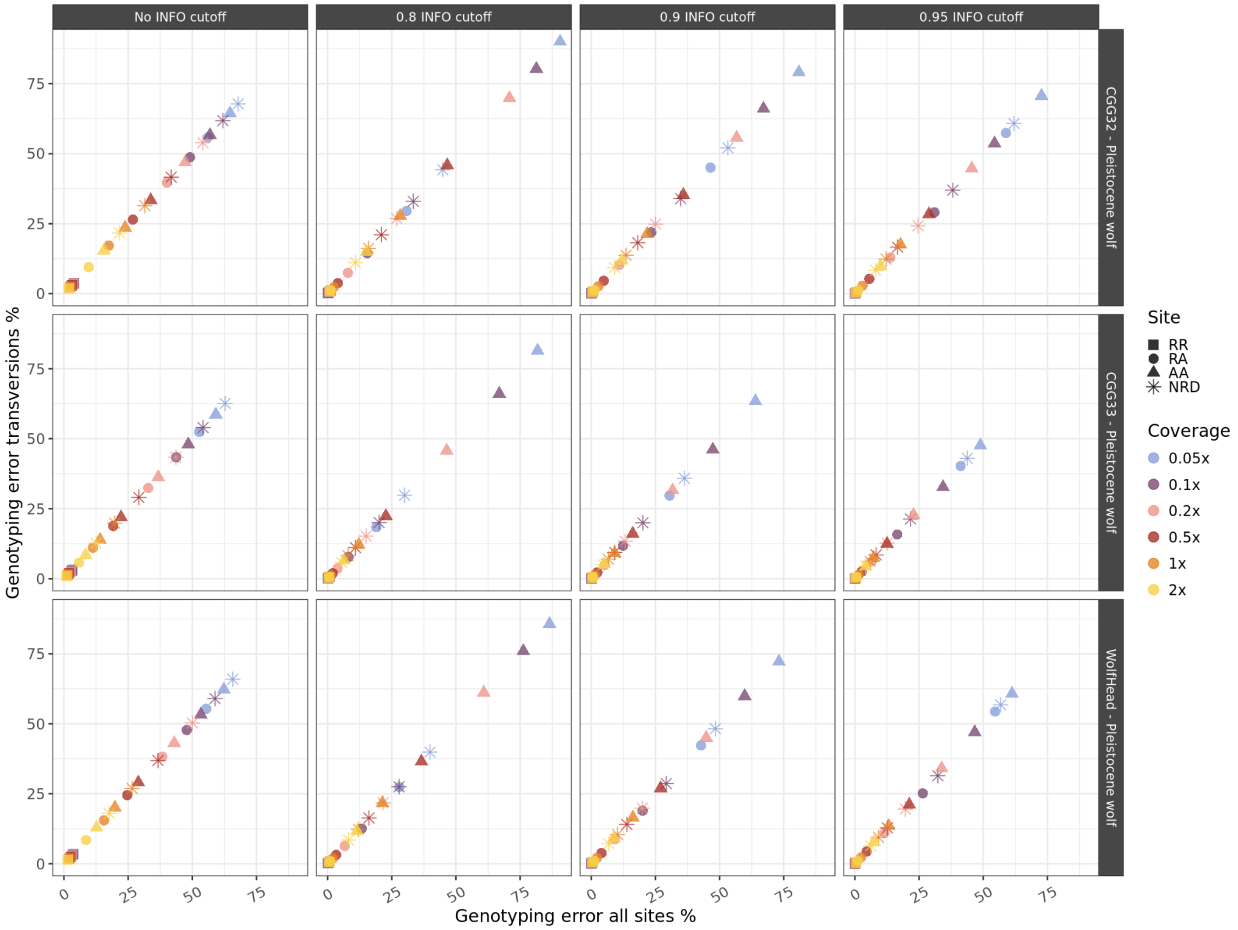


**Fig. S20:** *Genotyping error of transversions and transitions (all sites) against genotyping error of transversions for the three imputed target wolf samples (rows) applying different INFO score cutoffs (columns). Estimates are shown for genotyping errors of homozygous reference (RR), heterozygous (RA), homozygous alternative (AA) alleles and the non-reference discordance (NRD) metric. Colours represent different coverages.*

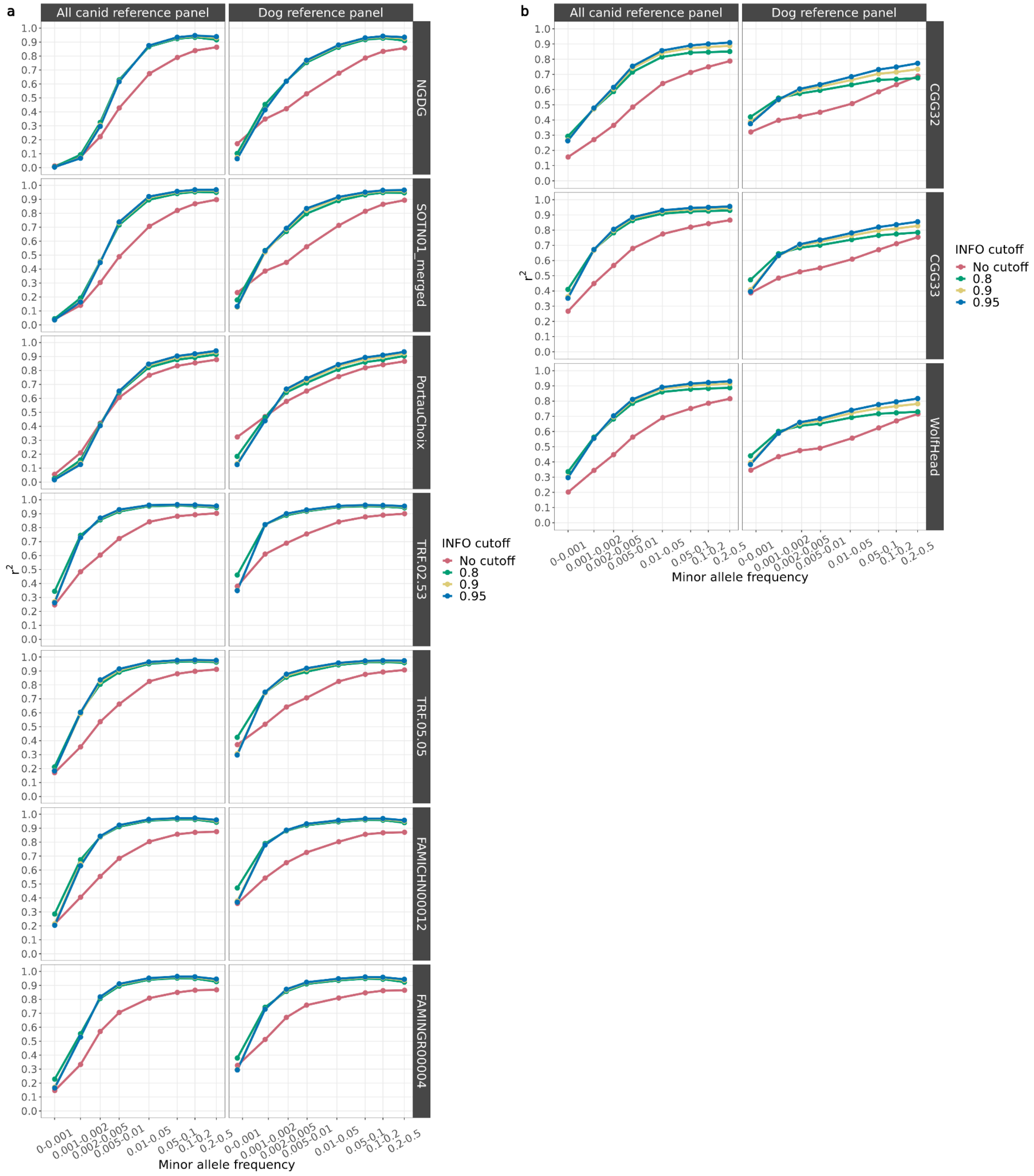


**Fig. S21:** *Squared correlation between imputed genotypes by GLIMPSE and highly confident called genotypes for all 10 target samples, downsampled to a) 0.5x for dogs and b) 1x for wolves and across different MAF bins using either the all canid reference panel or the reference panel containing only dogs. Each colour depicts the accuracy for a given INFO score cutoff. Red: no cut-off, Green: 0.8, Yellow: 0.9 and Blue: 0.95.*

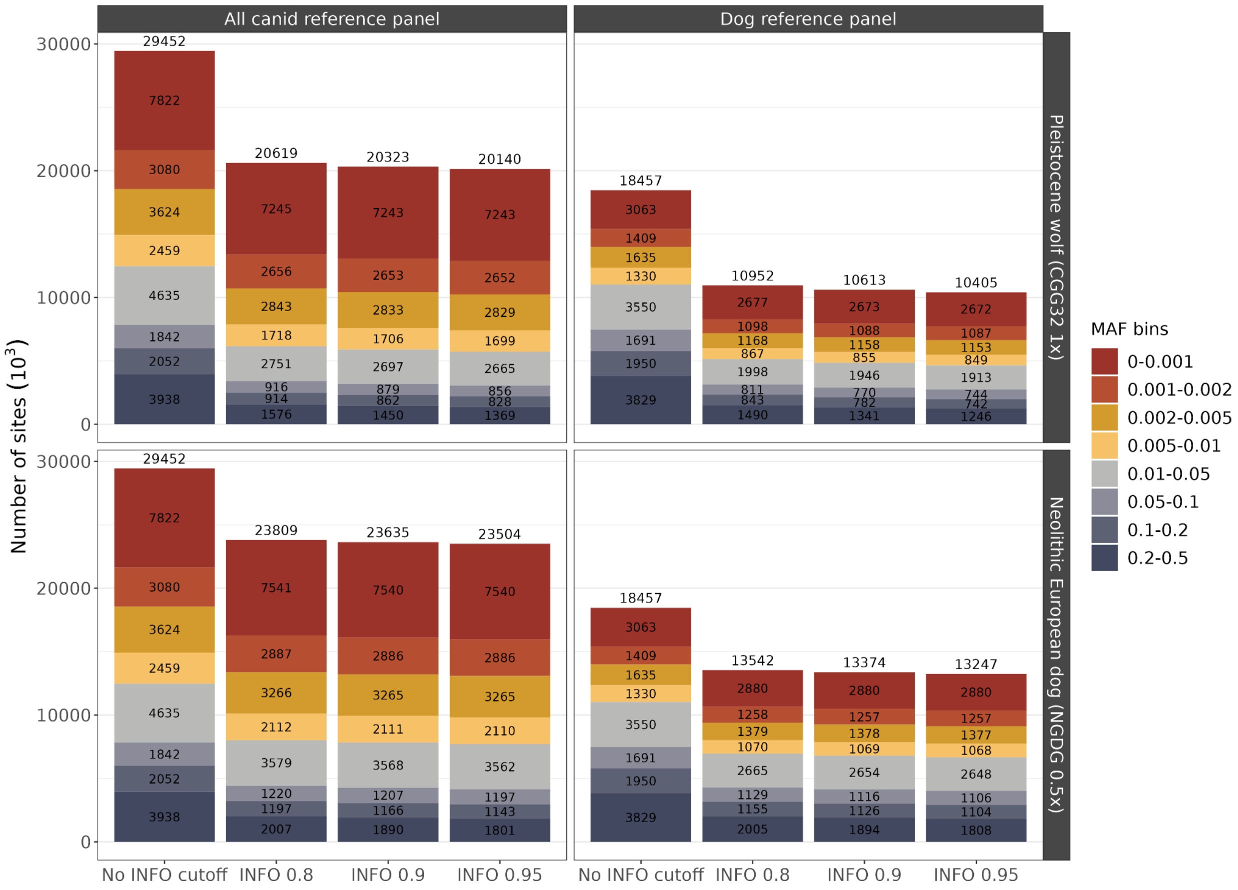


**Fig. S22:** *Number of imputed sites retained after applying INFO score cutoffs (no cutoff, >0.8, >0.9 and >0.95) across different minor allele frequency (MAF) bins for two imputed target samples (Dog: NGDG 0.5x, Wolf: CGG32 1x) using an all canid and a dog only reference panel. Numbers in each subgroup correspond to the number of sites ($10^3$) within that MAF bin. Numbers on top of each bar are the total number of sites ($10^3$) across all MAF bins.*

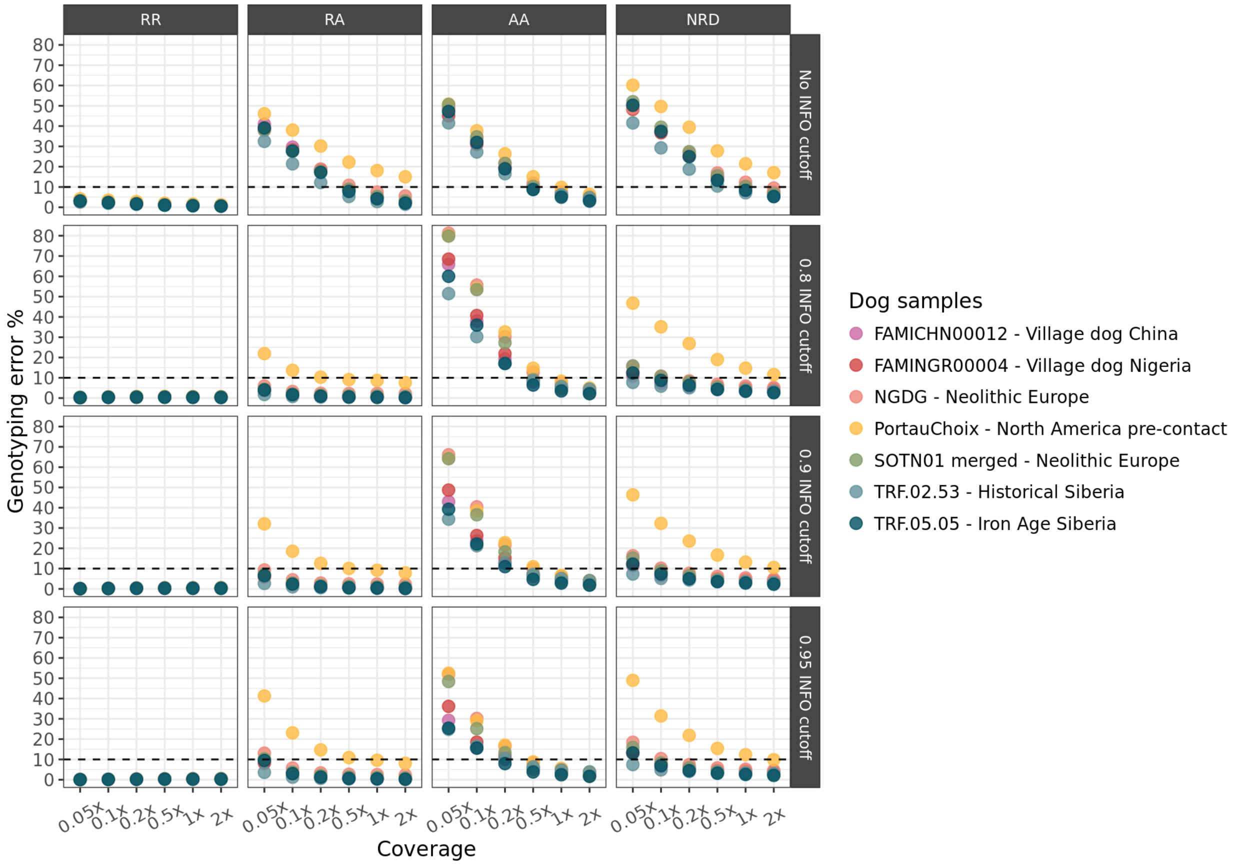


**Fig. S23:** *Genotyping error between imputed downsampled and the high coverage target dog samples imputed using a dog reference panel for homozygous alternative (AA), heterozygous (RA) and homozygous reference alleles (RR), and the non-reference discordance (NRD) metric estimated by GLIMPSE concordance for all autosomes. Comparisons are shown for different INFO score cutoffs (no cutoff, 0.8, 0.9 and 0.95).*

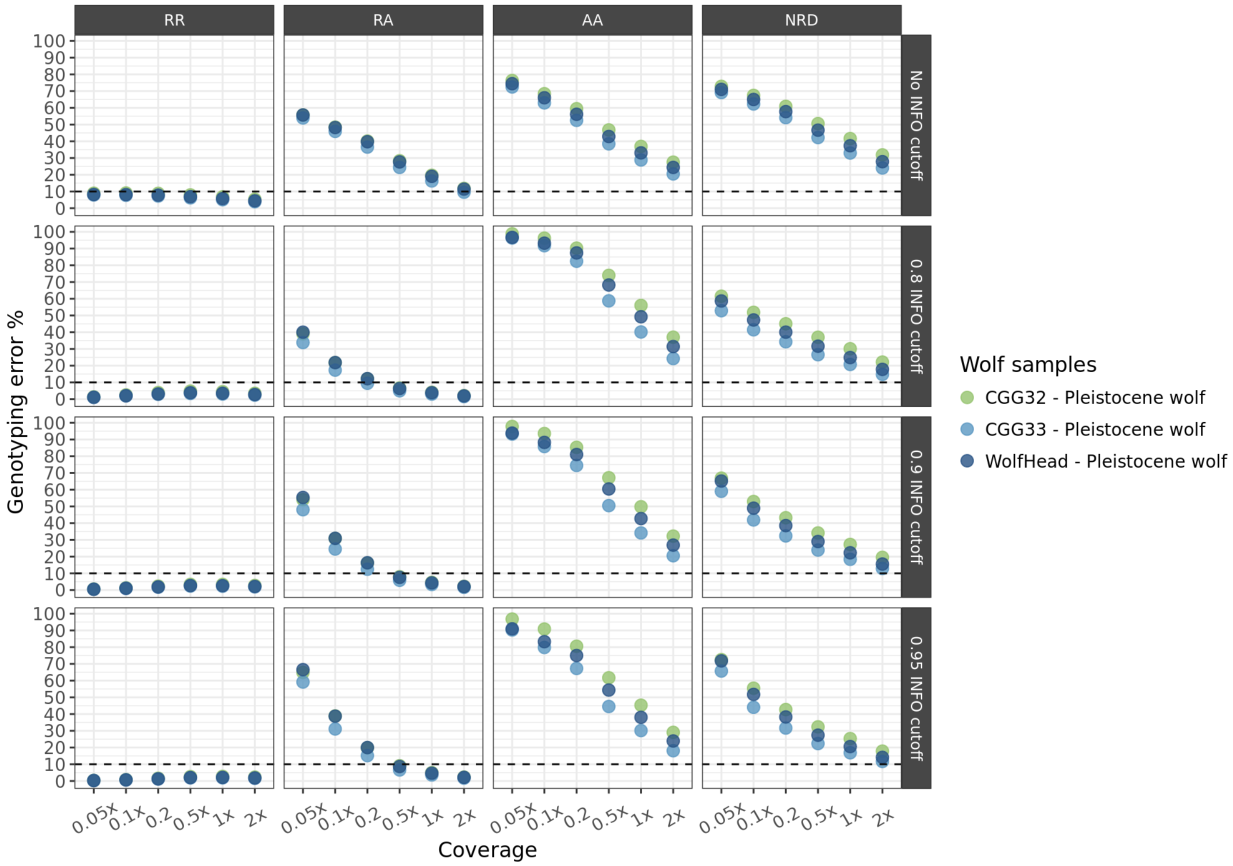


**Fig. S24:** *Genotyping error between imputed downsampled and the high coverage target Pleistocene wolf samples using a dog reference panel for homozygous alternative (AA), heterozygous (RA) and homozygous reference alleles (RR), and the non-reference discordance (NRD) metric estimated by GLIMPSE concordance for all autosomes. Comparisons are shown for different INFO score cutoffs (no cutoff, 0.8, 0.9 and 0.95).*

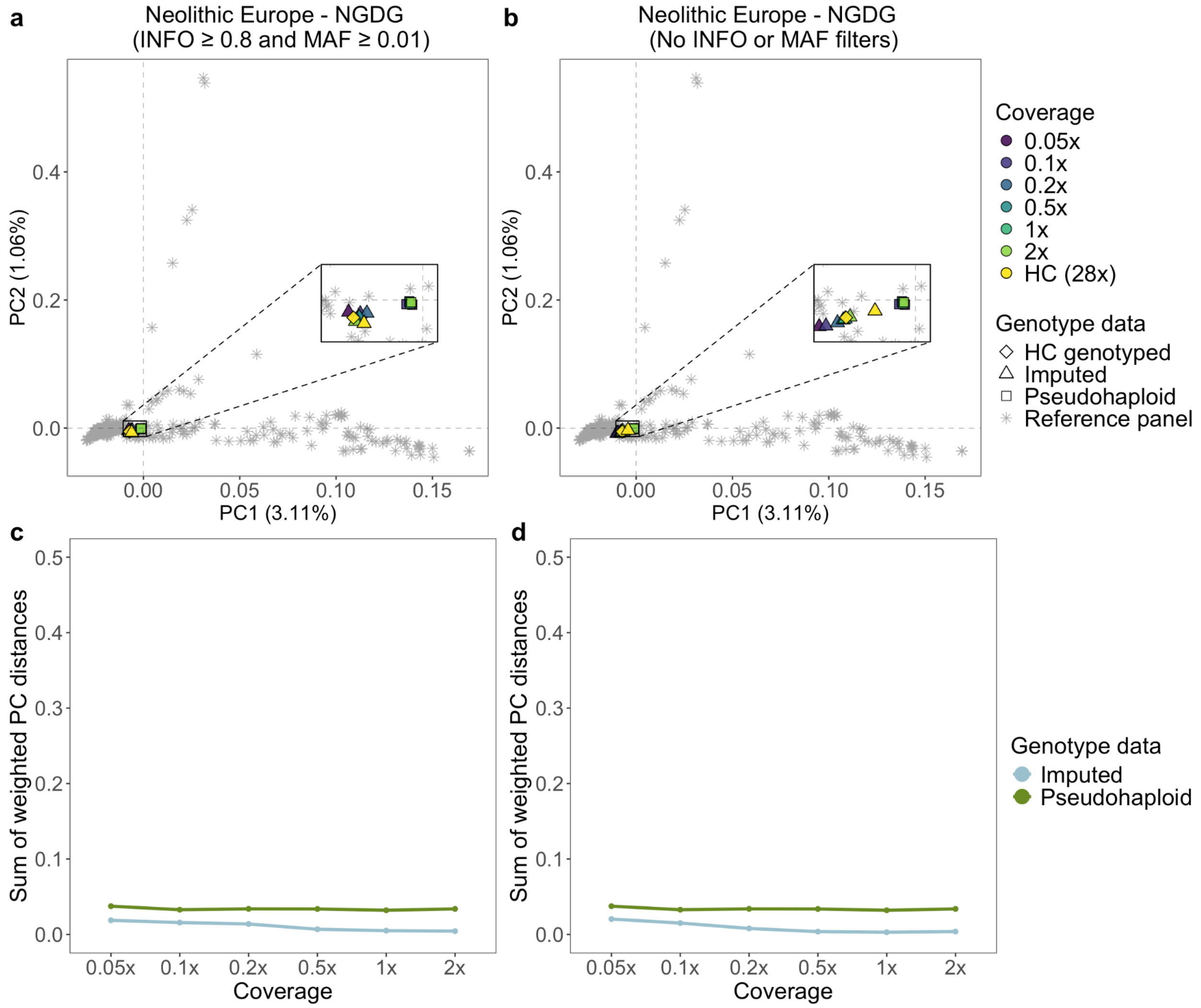


**Fig. S25:** *a, b) Principal component analysis showcasing the imputation accuracy for the Newgrange Neolithic European dog against its corresponding downsampled pseudohaploid counterpart. The PCs were created using modern dog samples from the reference panel, and then the imputed, pseudohaploid and high coverage genotyped replicas were projected onto them. c, d) Sum of weighted PC distances across all 10 PCs of the imputed and pseudohaploid downsampled individual from its high coverage genotyped version. The left plots (a,c) show the PCA results when applying INFO score and MAF cutoffs on the imputed samples, whereas the right ones (b,d) show when no post-imputation filter is applied. HC: High coverage.*

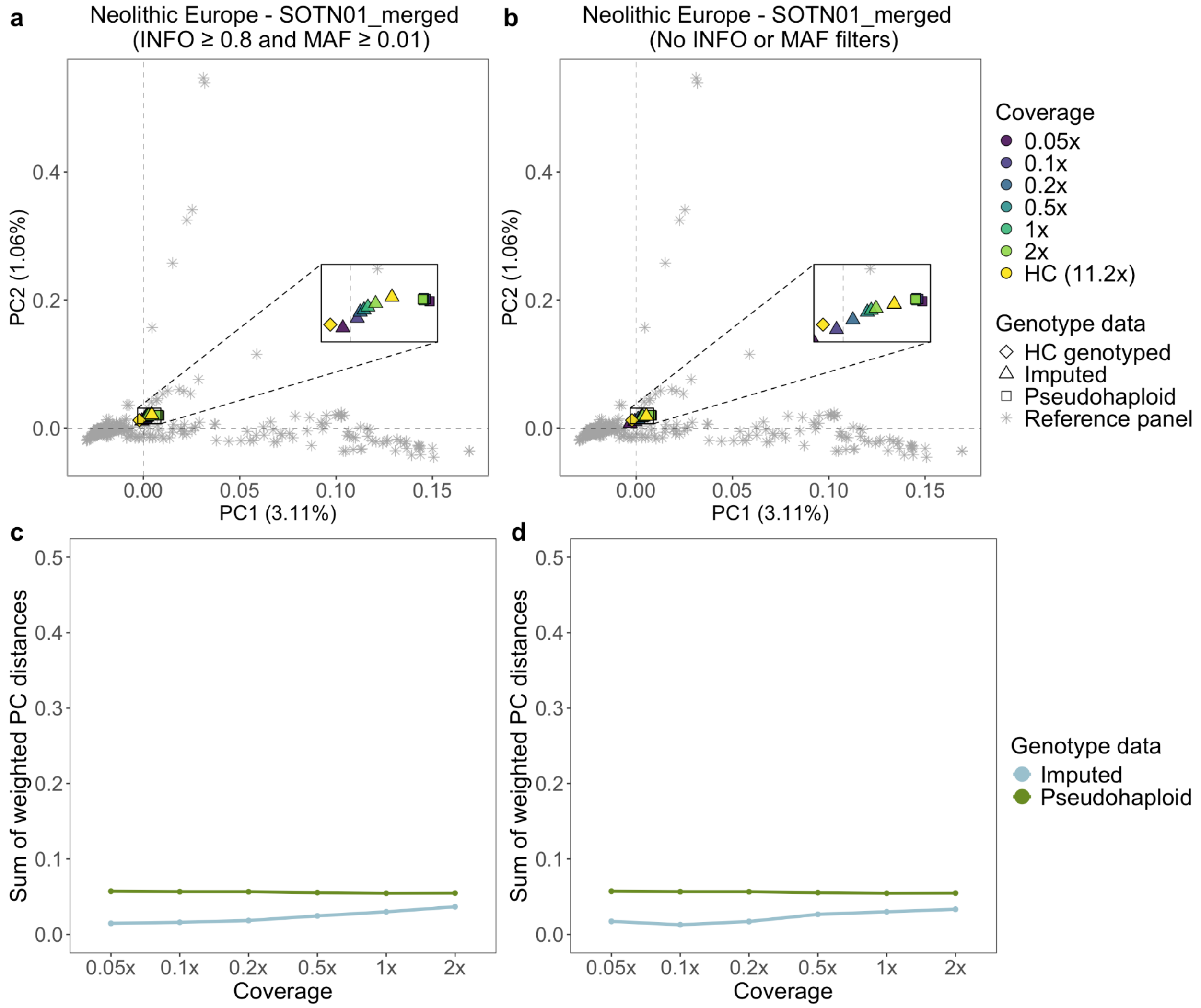


**Fig. S26:** *a, b) Principal component analysis showcasing the imputation accuracy for the SOTN01 Neolithic European dog against its corresponding downsampled pseudohaploid counterpart. The PCs were created using modern dog samples from the reference panel, and then the imputed, pseudohaploid and high coverage genotyped replicas were projected onto them. c, d) Sum of weighted PC distances across all 10 PCs of the imputed and pseudohaploid downsampled individual from its high coverage genotyped version. The left plots (a,c) show the PCA results when applying INFO score and MAF cutoffs on the imputed samples, whereas the right ones (b,d) show when no post-imputation filter is applied. HC: High coverage.*

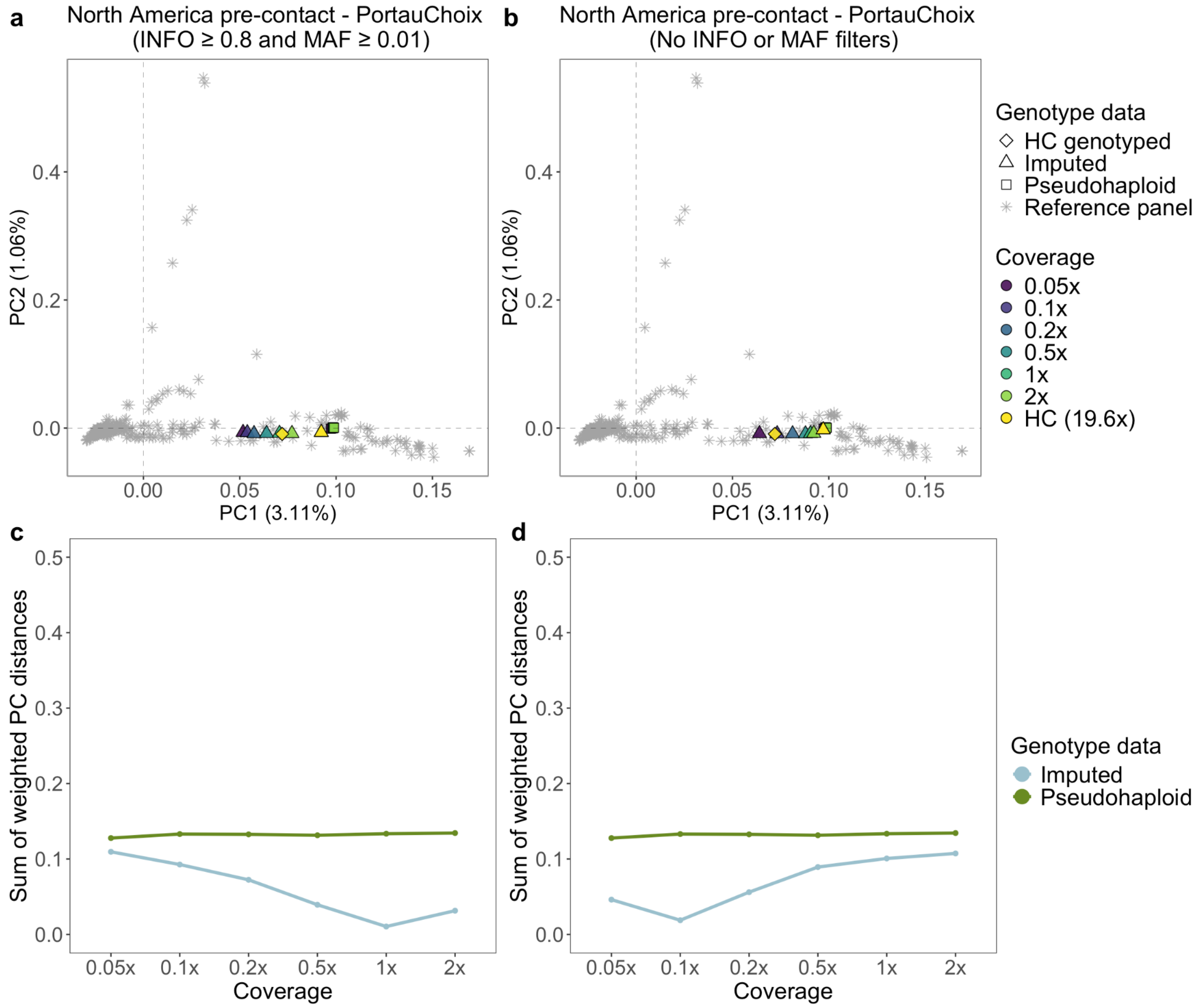


**Fig. S27:** *a, b) Principal component analysis showcasing the imputation accuracy for the Port au Choix North American pre-contact dog against its corresponding downsampled pseudohaploid counterpart. The PCs were created using modern dog samples from the reference panel, and then the imputed, pseudohaploid and high coverage genotyped replicas were projected onto them. c, d) Sum of weighted PC distances across all 10 PCs of the imputed and pseudohaploid downsampled individual from its high coverage genotyped version. The left plots (a,c) show the PCA results when applying INFO score and MAF cutoffs on the imputed samples, whereas the right ones (b,d) show when no post-imputation filter is applied. HC: High coverage.*

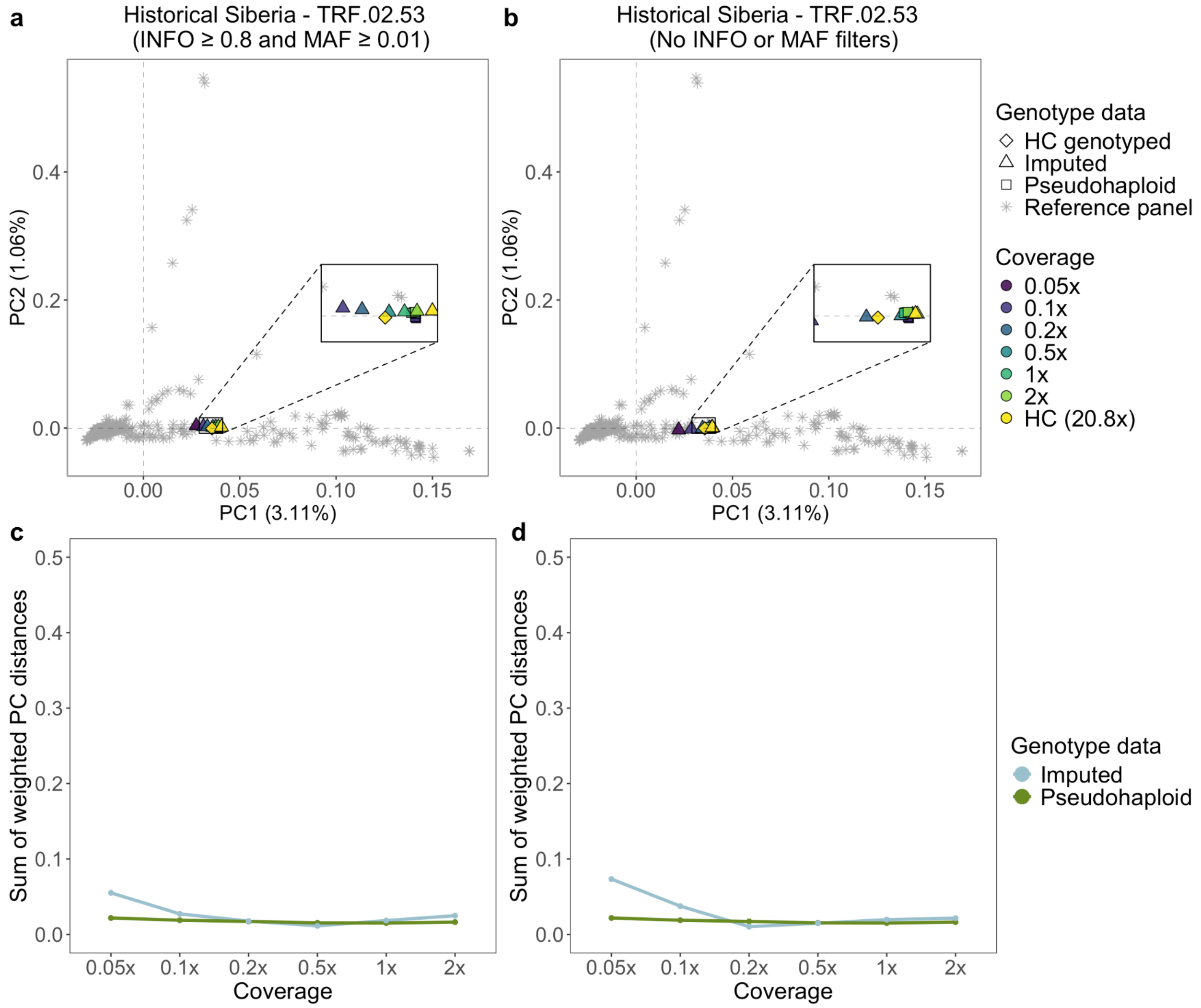


**Fig. S28:** *a, b) Principal component analysis showcasing the imputation accuracy for the TRF.02.53 historical Siberian dog against its corresponding downsampled pseudohaploid counterpart. The PCs were created using modern dog samples from the reference panel, and then the imputed, pseudohaploid and high coverage genotyped replicas were projected onto them. c, d) Sum of weighted PC distances across all 10 PCs of the imputed and pseudohaploid downsampled individual from its high coverage genotyped version. The left plots (a,c) show the PCA results when applying INFO score and MAF cutoffs on the imputed samples, whereas the right ones (b,d) show when no post-imputation filter is applied. HC: High coverage.*

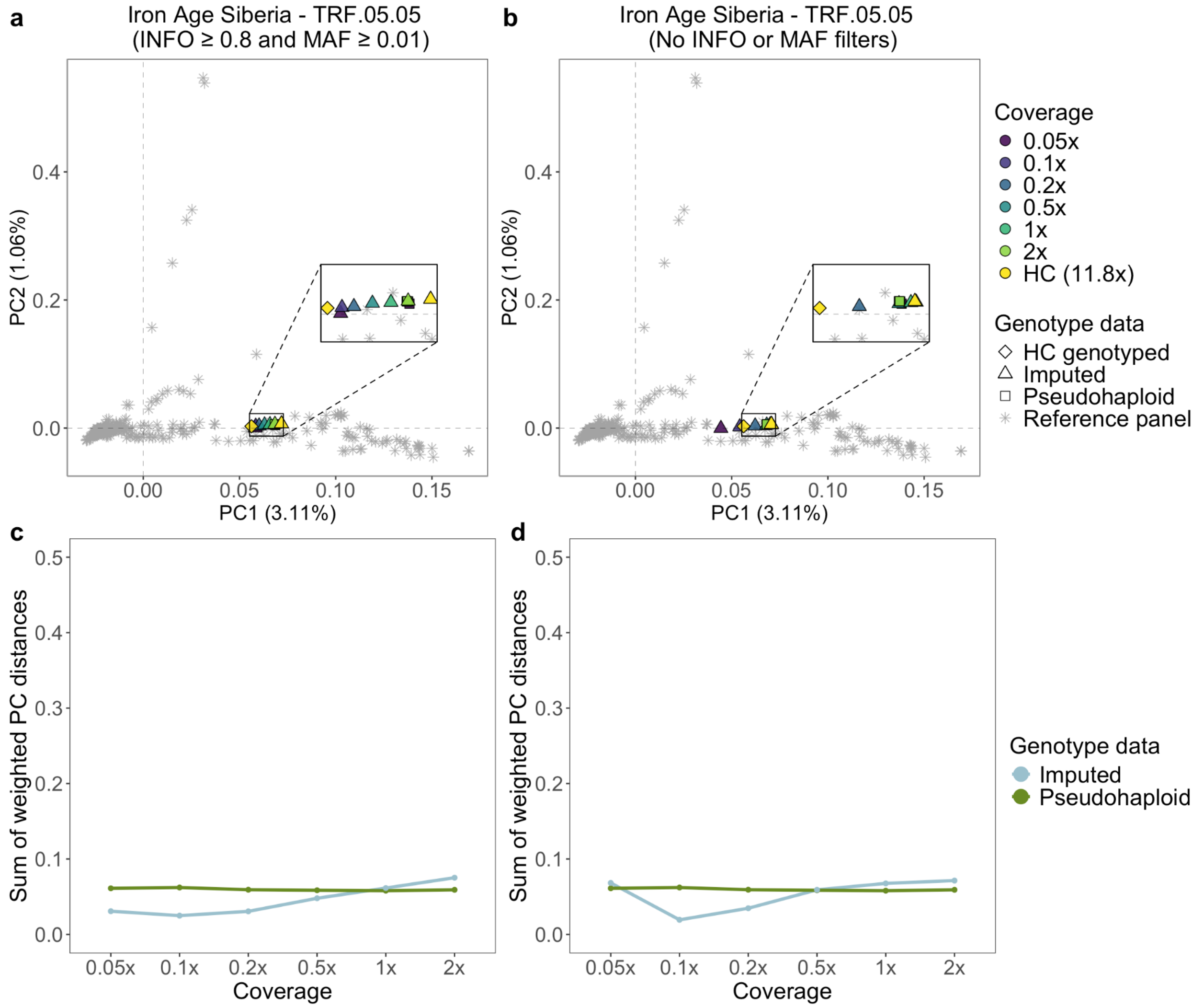


**Fig. S29:** *a, b) Principal component analysis showcasing the imputation accuracy for the TRF.05.05 Iron Age Siberian dog against its corresponding downsampled pseudohaploid counterpart. The PCs were created using modern dog samples from the reference panel, and then the imputed, pseudohaploid and high coverage genotyped replicas were projected onto them. c, d) Sum of weighted PC distances across all 10 PCs of the imputed and pseudohaploid downsampled individual from its high coverage genotyped version. The left plots (a,c) show the PCA results when applying INFO score and MAF cutoffs on the imputed samples, whereas the right ones (b,d) show when no post-imputation filter is applied. HC: High coverage.*

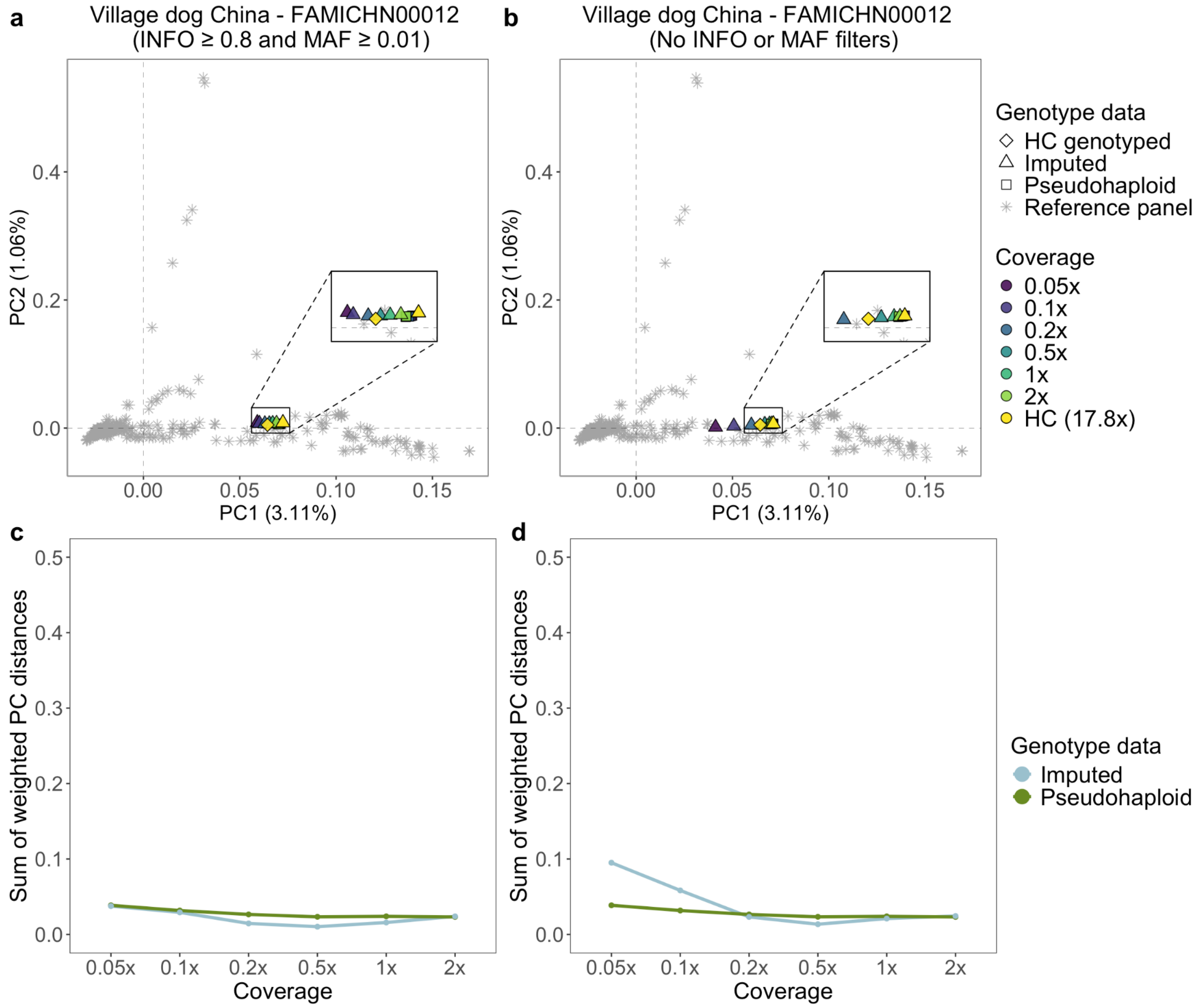


**Fig. S30:** *a, b) Principal component analysis showcasing the imputation accuracy for the FAMICHN00012 Chinese village dog against its corresponding downsampled pseudohaploid counterpart. The PCs were created using modern dog samples from the reference panel, and then the imputed, pseudohaploid and high coverage genotyped replicas were projected onto them. c, d) Sum of weighted PC distances across all 10 PCs of the imputed and pseudohaploid downsampled individual from its high coverage genotyped version. The left plots (a,c) show the PCA results when applying INFO score and MAF cutoffs on the imputed samples, whereas the right ones (b,d) show when no post-imputation filter is applied. HC: High coverage.*

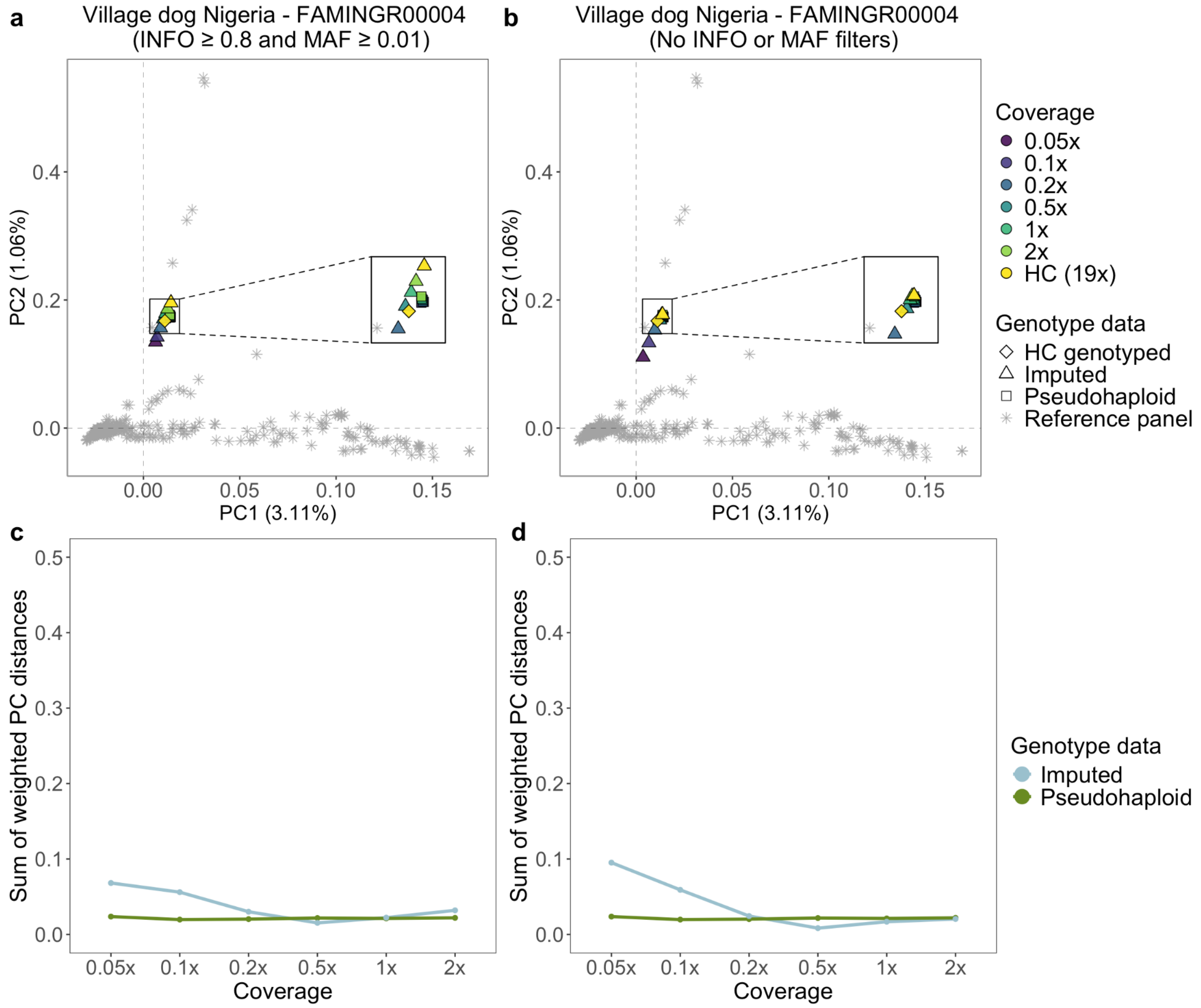


**Fig. S31:** *a, b) Principal component analysis showcasing the imputation accuracy for the FAMINGR00004 Nigerian village dog against its corresponding downsampled pseudohaploid counterpart. The PCs were created using modern dog samples from the reference panel, and then the imputed, pseudohaploid and high coverage genotyped replicas were projected onto them. c, d) Sum of weighted PC distances across all 10 PCs of the imputed and pseudohaploid downsampled individual from its high coverage genotyped version. The left plots (a,c) show the PCA results when applying INFO score and MAF cutoffs on the imputed samples, whereas the right ones (b,d) show when no post-imputation filter is applied. HC: High coverage.*

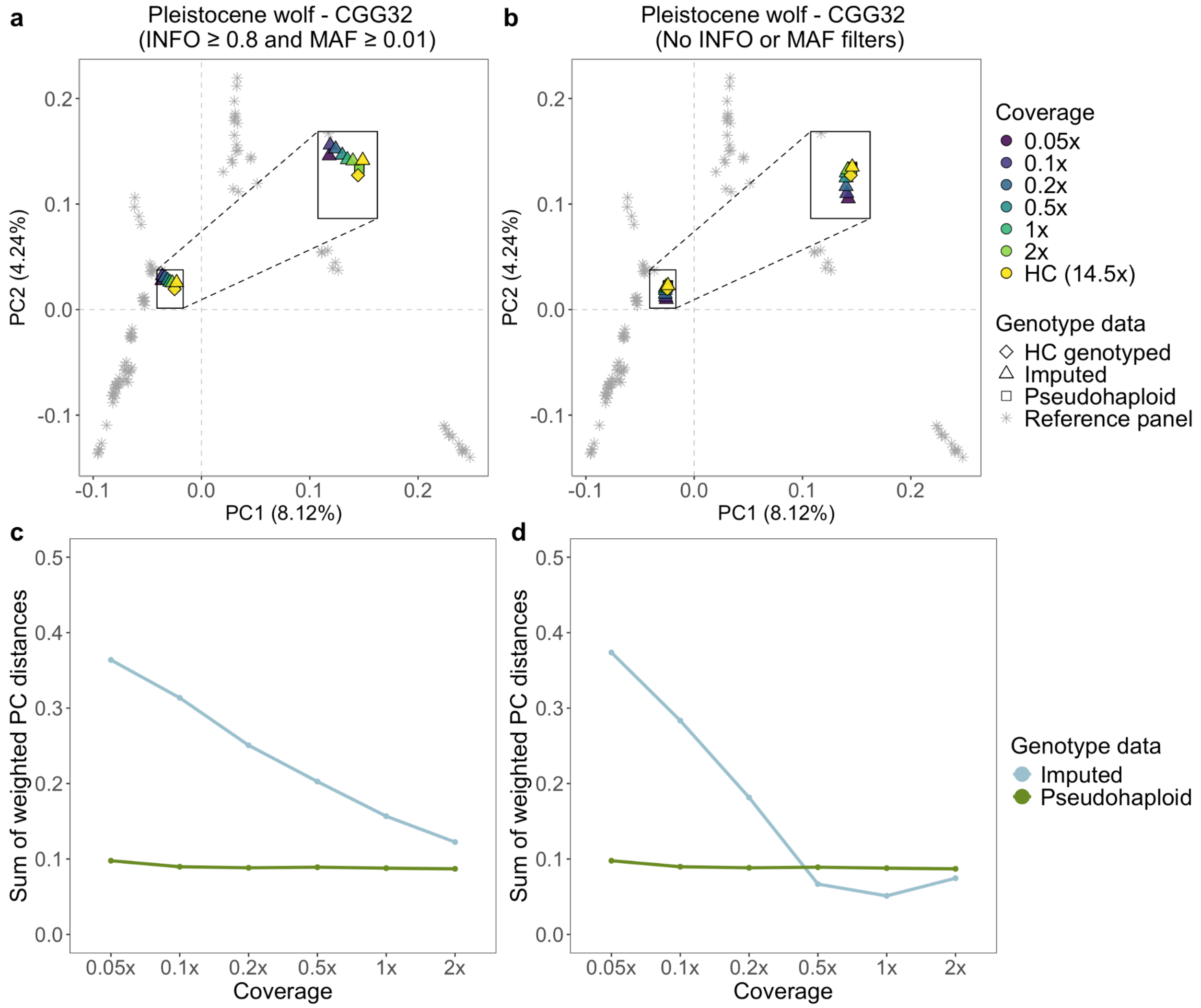


**Fig. S32:** *a, b) Principal component analysis showcasing the imputation accuracy for the CGG32 Pleistocene wolf against its corresponding downsampled pseudohaploid counterpart. The PCs were created using modern wolf samples from the reference panel, and then the imputed, pseudohaploid and high coverage genotyped replicas were projected onto them. c, d) Sum of weighted PC distances across all 10 PCs of the imputed and pseudohaploid downsampled individual from its high coverage genotyped version. The left plots (a,c) show the PCA results when applying INFO score and MAF cutoffs on the imputed samples, whereas the right ones (b,d) show when no post-imputation filter is applied. HC: High coverage.*

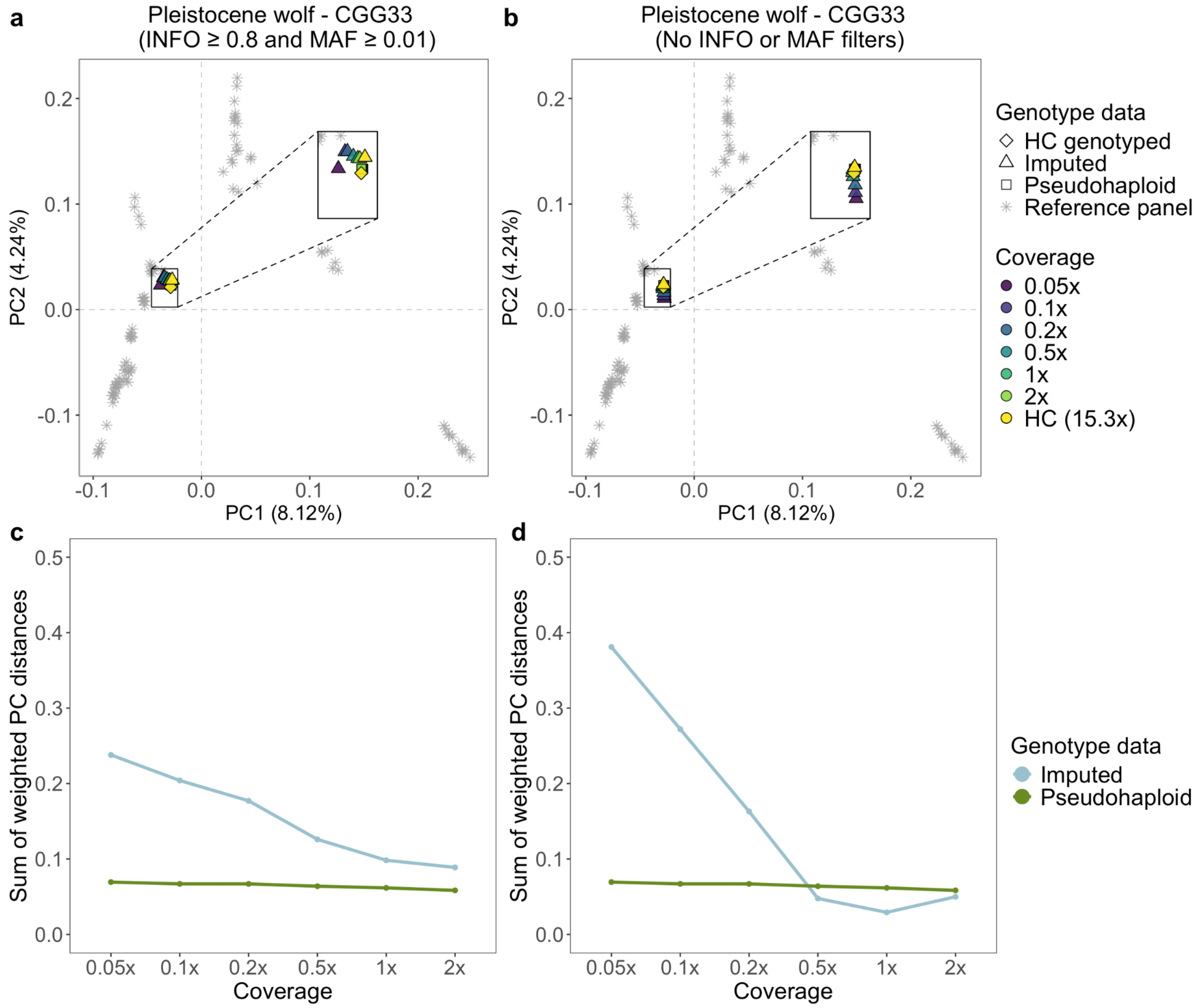


**Fig. S33:** *a, b) Principal component analysis showcasing the imputation accuracy for the CGG33 Pleistocene wolf against its corresponding downsampled pseudohaploid counterpart. The PCs were created using modern wolf samples from the reference panel, and then the imputed, pseudohaploid and high coverage genotyped replicas were projected onto them. c, d) Sum of weighted PC distances across all 10 PCs of the imputed and pseudohaploid downsampled individual from its high coverage genotyped version. The left plots (a,c) show the PCA results when applying INFO score and MAF cutoffs on the imputed samples, whereas the right ones (b,d) show when no post-imputation filter is applied. HC: High coverage.*

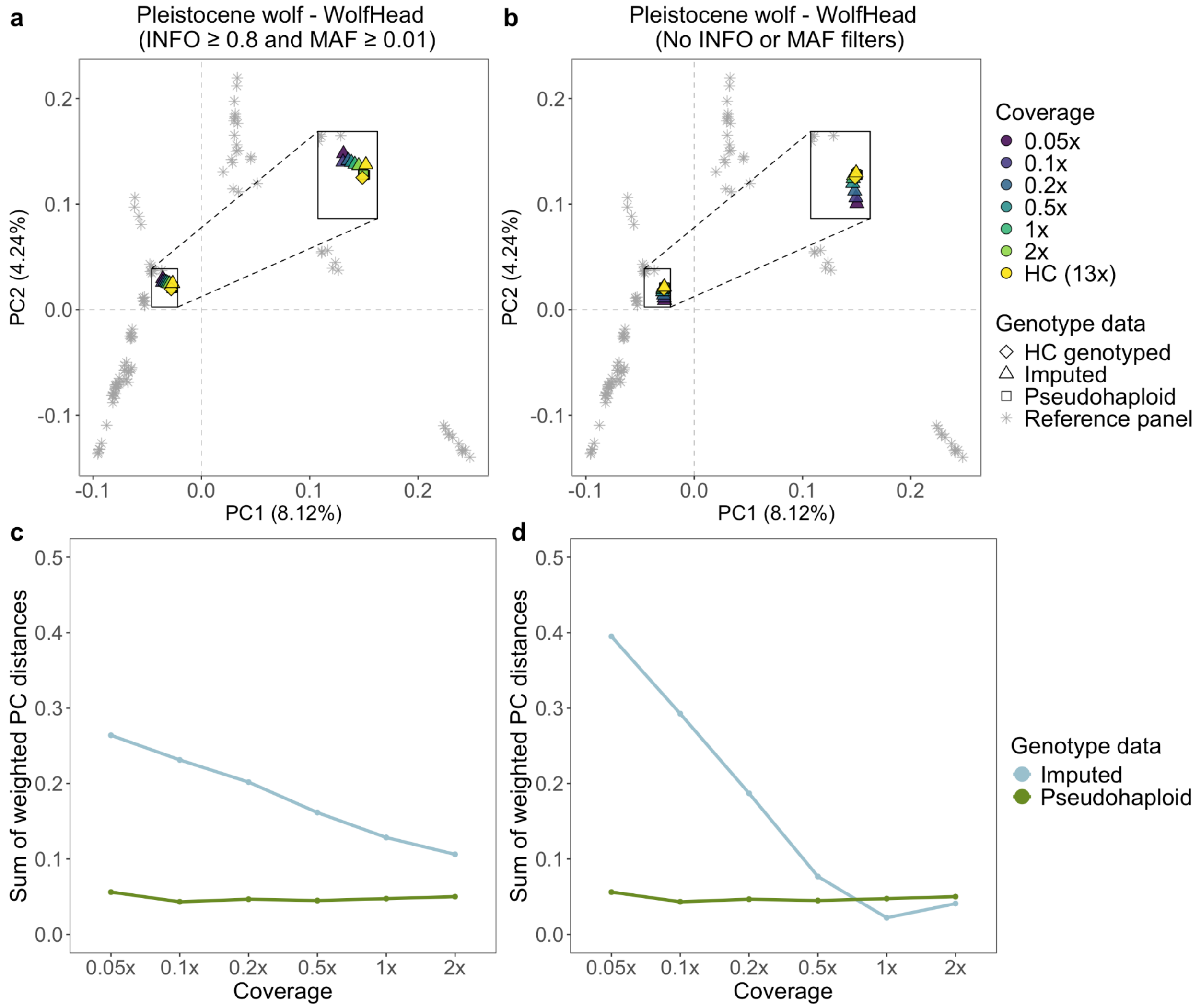


**Fig. S34:** *a, b) Principal component analysis showcasing the imputation accuracy for the WolfHead Pleistocene wolf against its corresponding downsampled pseudohaploid counterpart. The PCs were created using modern wolf samples from the reference panel, and then the imputed, pseudohaploid and high coverage genotyped replicas were projected onto them. c, d) Sum of weighted PC distances across all 10 PCs of the imputed and pseudohaploid downsampled individual from its high coverage genotyped version. The left plots (a,c) show the PCA results when applying INFO score and MAF cutoffs on the imputed samples, whereas the right ones (b,d) show when no post-imputation filter is applied. HC: High coverage.*

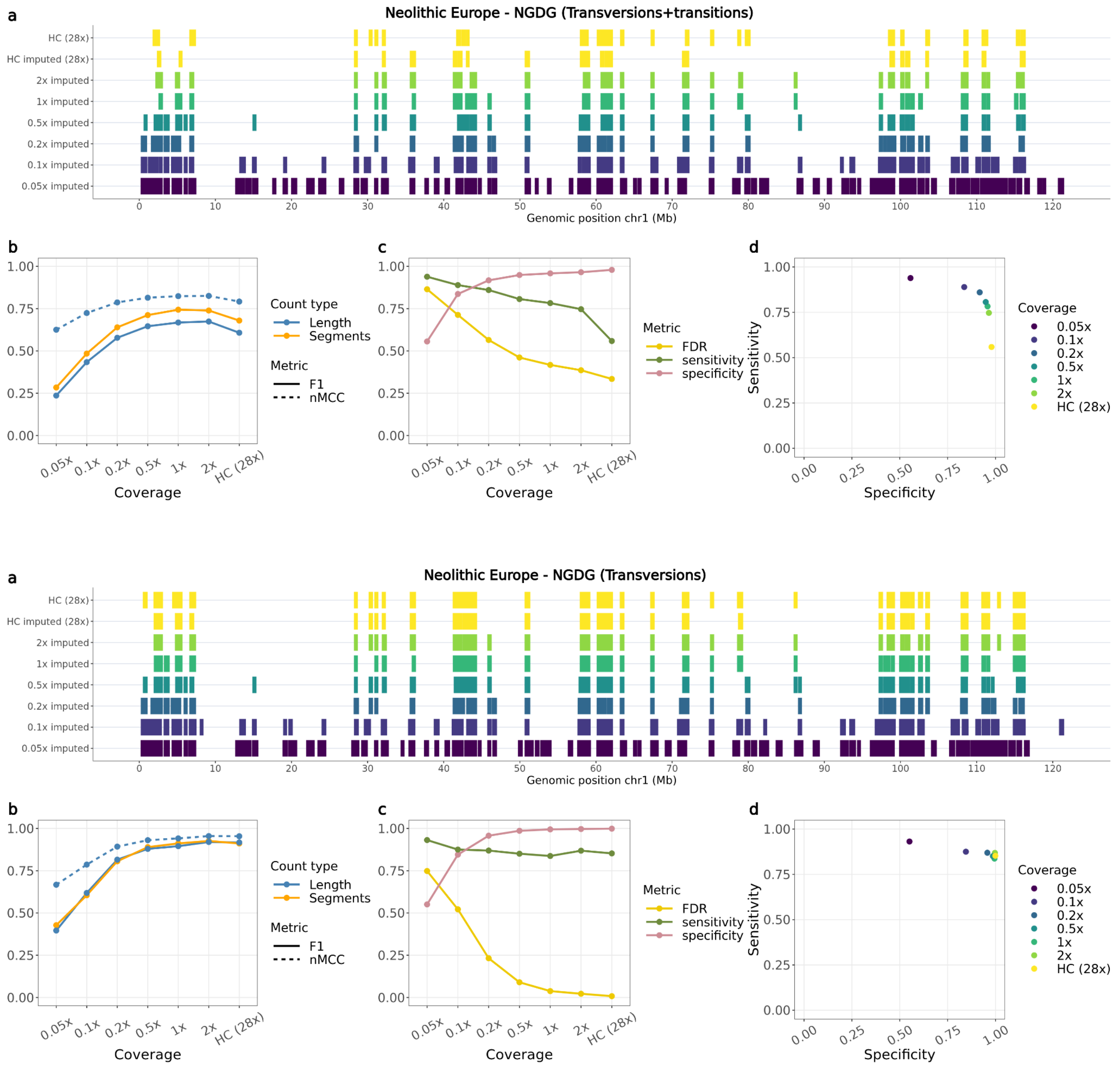


**Fig. S35:** *Overlap of ROH called from the Newgrange Neolithic European dog for each imputed downsampled replicate using ROH estimates from the ground truth, using transversions and transitions (top panel) or only transversions (bottom panel). a) ROH called across the six tested coverages and the high coverage imputed and genotyped (ground truth) sample on chromosome one. b) Accuracy of recovering ROH across all tested coverages based on total length in bp (blue lines) and total number of segments (orange line) using the F1-score (solid line) and normalised Matthew correlation coefficient (nMCC) (dotted line). c) FDR, sensitivity and specificity measurements based on the total length of recovered ROH per coverage. d) Sensitivity plotted against specificity estimated based on the total length of recovered ROH across all tested coverages. HC: High coverage.*

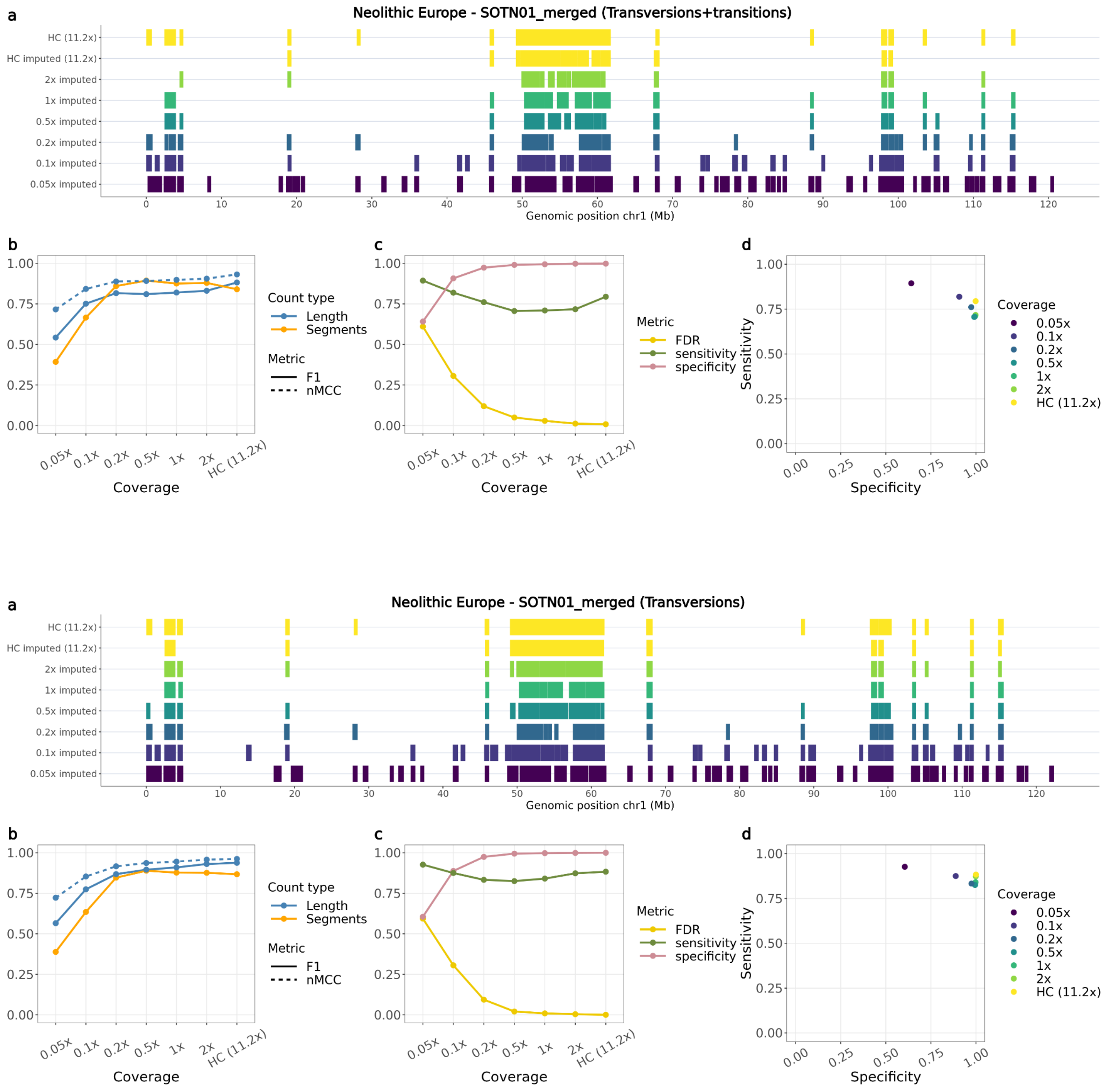


**Fig. S36:** *Overlap of ROH called from the SOTN01 Neolithic European dog for each imputed downsampled replicate using ROH estimates from the ground truth, using transversions and transitions (top panel) or only transversions (bottom panel). a) ROH called across the six tested coverages and the high coverage imputed and genotyped (ground truth) sample on chromosome one. b) Accuracy of recovering ROH across all tested coverages based on total length in bp (blue lines) and total number of segments (orange line) using the F1-score (solid line) and normalised Matthew correlation coefficient (nMCC) (dotted line). c) FDR, sensitivity and specificity measurements based on the total length of recovered ROH per coverage. d) Sensitivity plotted against specificity estimated based on the total length of recovered ROH across all tested coverages. HC: High coverage.*

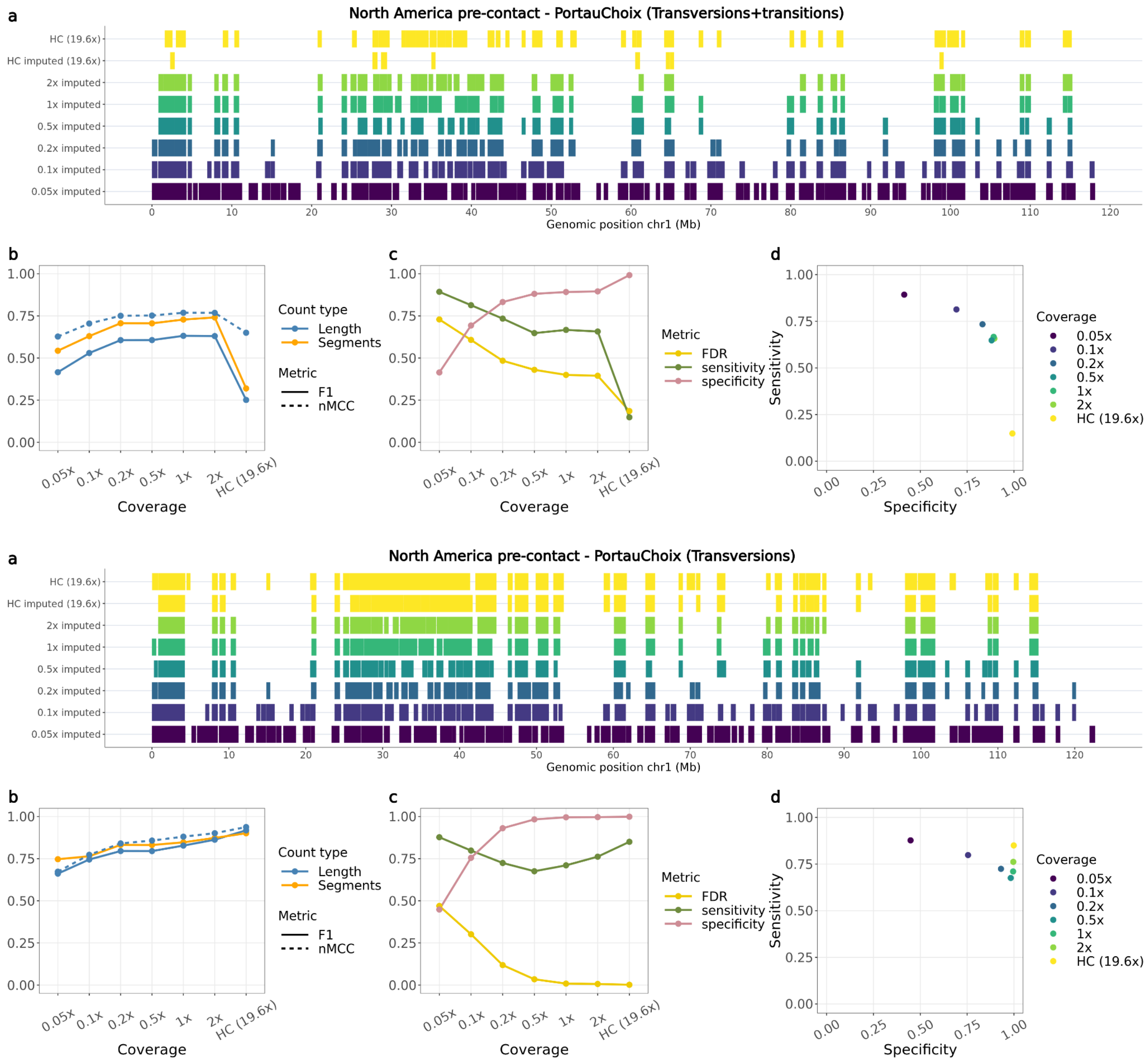


**Fig. S37:** *Overlap of ROH called from the Port au Choix North American pre-contact dog for each imputed downsampled replicate using ROH estimates from the ground truth, using transversions and transitions (top panel) or only transversions (bottom panel). a) ROH called across the six tested coverages and the high coverage imputed and genotyped (ground truth) sample on chromosome one. b) Accuracy of recovering ROH across all tested coverages based on total length in bp (blue lines) and total number of segments (orange line) using the F1-score (solid line) and normalised Matthew correlation coefficient (nMCC) (dotted line). c) FDR, sensitivity and specificity measurements based on the total length of recovered ROH per coverage. d) Sensitivity plotted against specificity estimated based on the total length of recovered ROH across all tested coverages. HC: High coverage.*

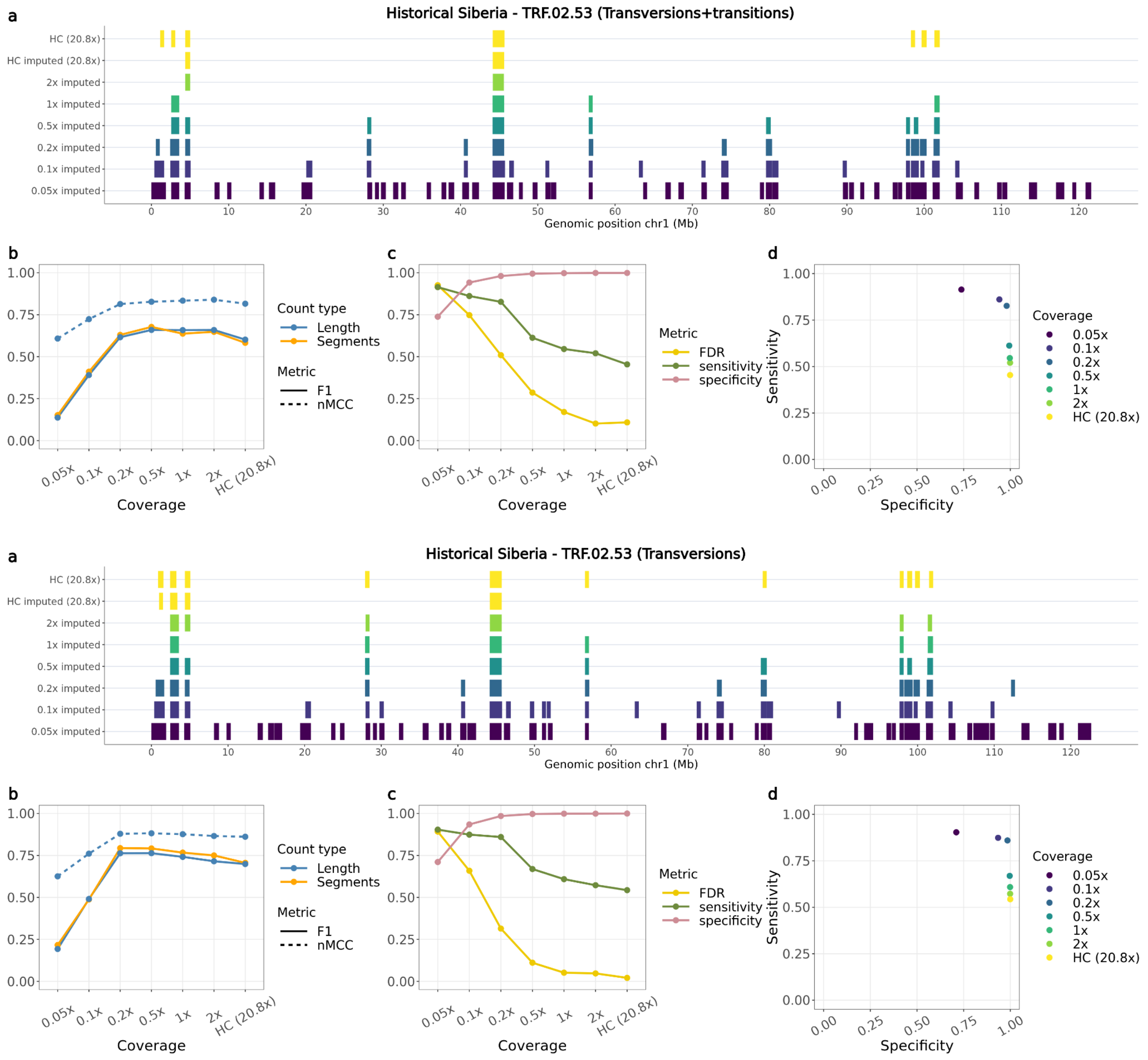


**Fig. S38:** *Overlap of ROH called from the TRF.02.53 historical Siberian dog for each imputed downsampled replicate using ROH estimates from the ground truth, using transversions and transitions (top panel) or only transversions (bottom panel). a) ROH called across the six tested coverages and the high coverage imputed and genotyped (ground truth) sample on chromosome one. b) Accuracy of recovering ROH across all tested coverages based on total length in bp (blue lines) and total number of segments (orange line) using the F1-score (solid line) and normalised Matthew correlation coefficient (nMCC) (dotted line). c) FDR, sensitivity and specificity measurements based on the total length of recovered ROH per coverage. d) Sensitivity plotted against specificity estimated based on the total length of recovered ROH across all tested coverages. HC: High coverage.*

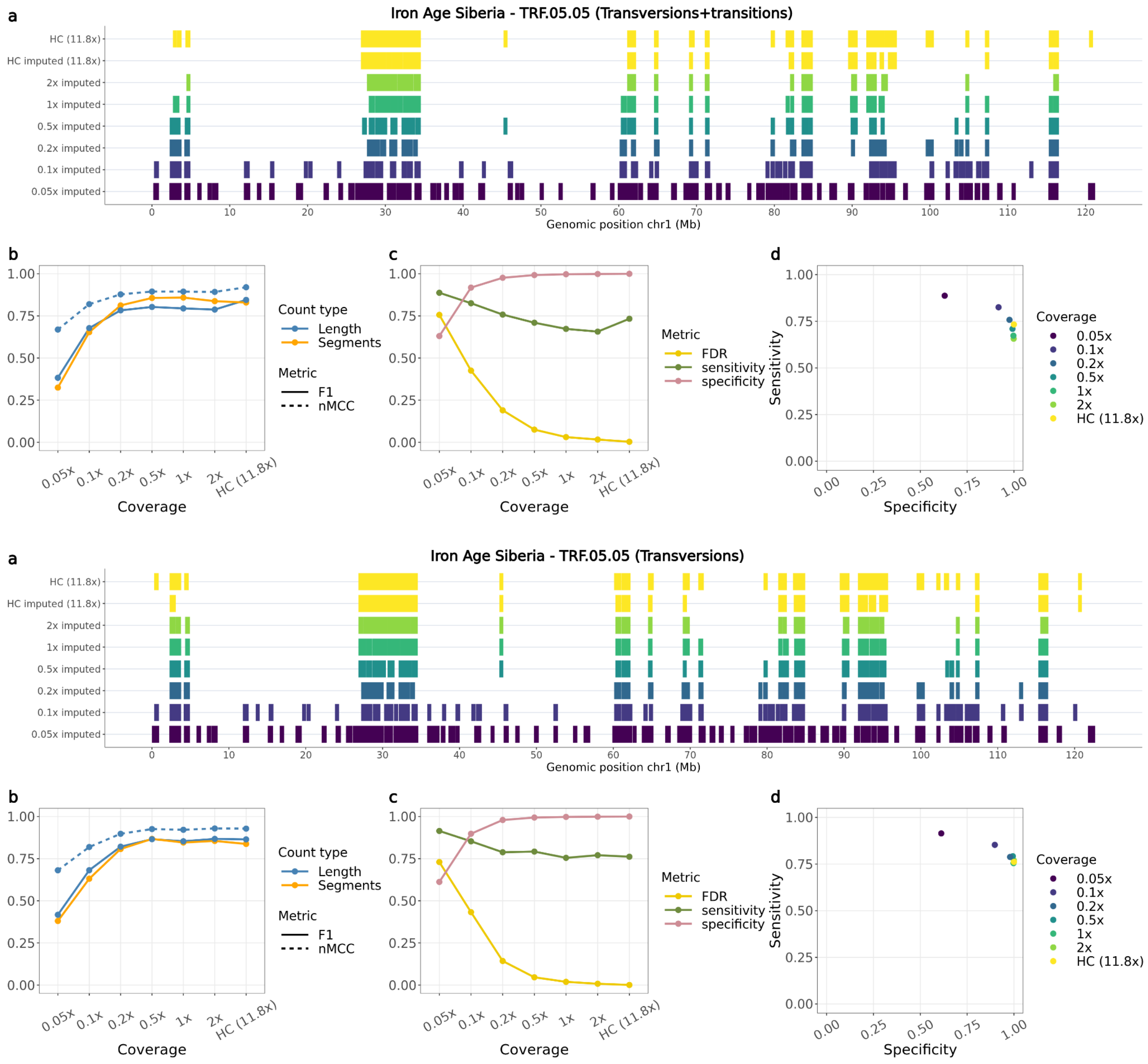


**Fig. S39:** *Overlap of ROH called from the TRF.05.05 Iron Age Siberian dog for each imputed downsampled replicate using ROH estimates from the ground truth, using transversions and transitions (top panel) or only transversions (bottom panel). a) ROH called across the six tested coverages and the high coverage imputed and genotyped (ground truth) sample on chromosome one. b) Accuracy of recovering ROH across all tested coverages based on total length in bp (blue lines) and total number of segments (orange line) using the F1-score (solid line) and normalised Matthew correlation coefficient (nMCC) (dotted line). c) FDR, sensitivity and specificity measurements based on the total length of recovered ROH per coverage. d) Sensitivity plotted against specificity estimated based on the total length of recovered ROH across all tested coverages. HC: High coverage.*

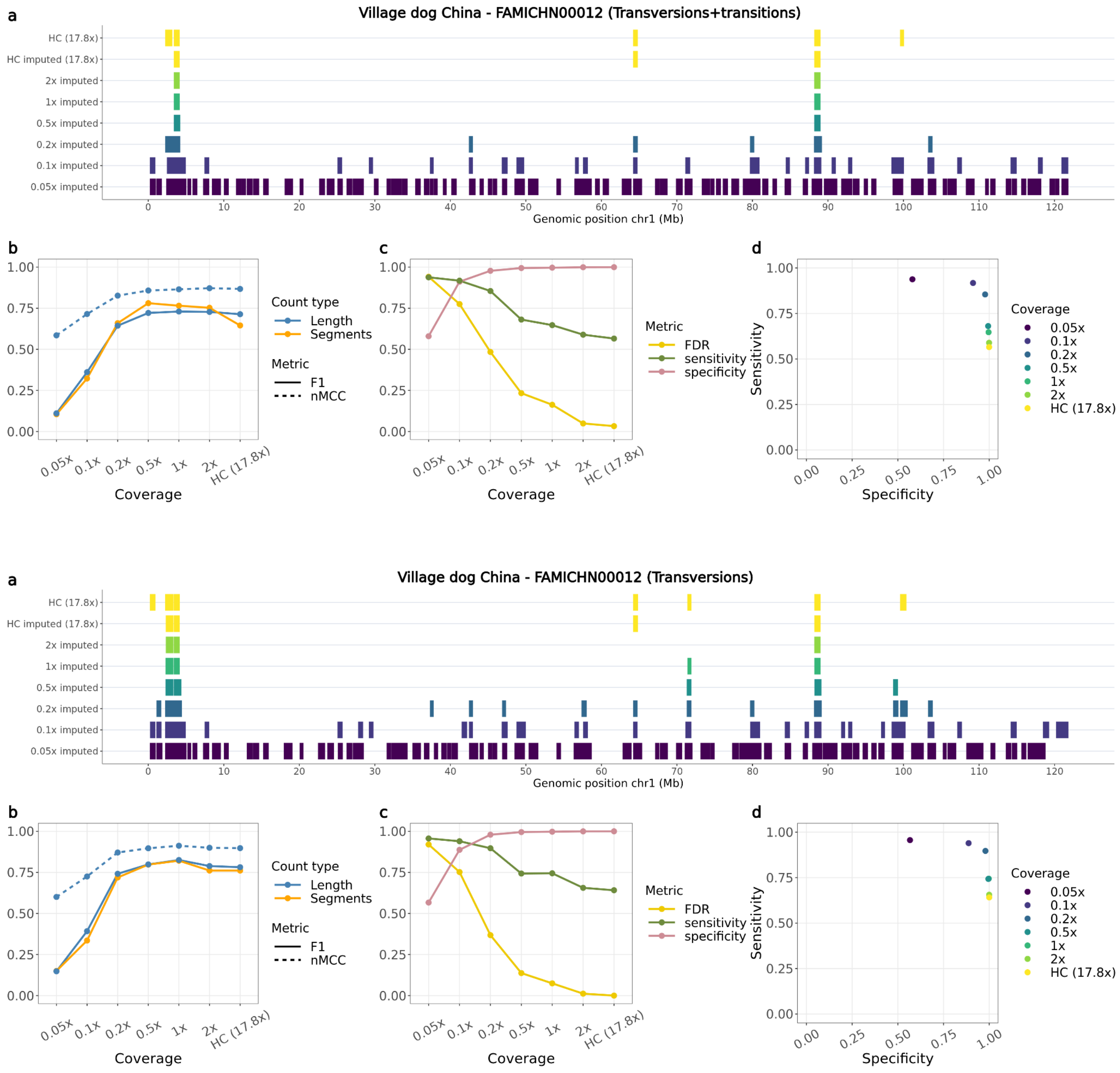


**Fig. S40:** *Overlap of ROH called from the FAMICHN00012 Chinese Village dog for each imputed downsampled replicate using ROH estimates from the ground truth, using transversions and transitions (top panel) or only transversions (bottom panel). a) ROH called across the six tested coverages and the high coverage imputed and genotyped (ground truth) sample on chromosome one. b) Accuracy of recovering ROH across all tested coverages based on total length in bp (blue lines) and total number of segments (orange line) using the F1-score (solid line) and normalised Matthew correlation coefficient (nMCC) (dotted line). c) FDR, sensitivity and specificity measurements based on the total length of recovered ROH per coverage. d) Sensitivity plotted against specificity estimated based on the total length of recovered ROH across all tested coverages. HC: High coverage.*

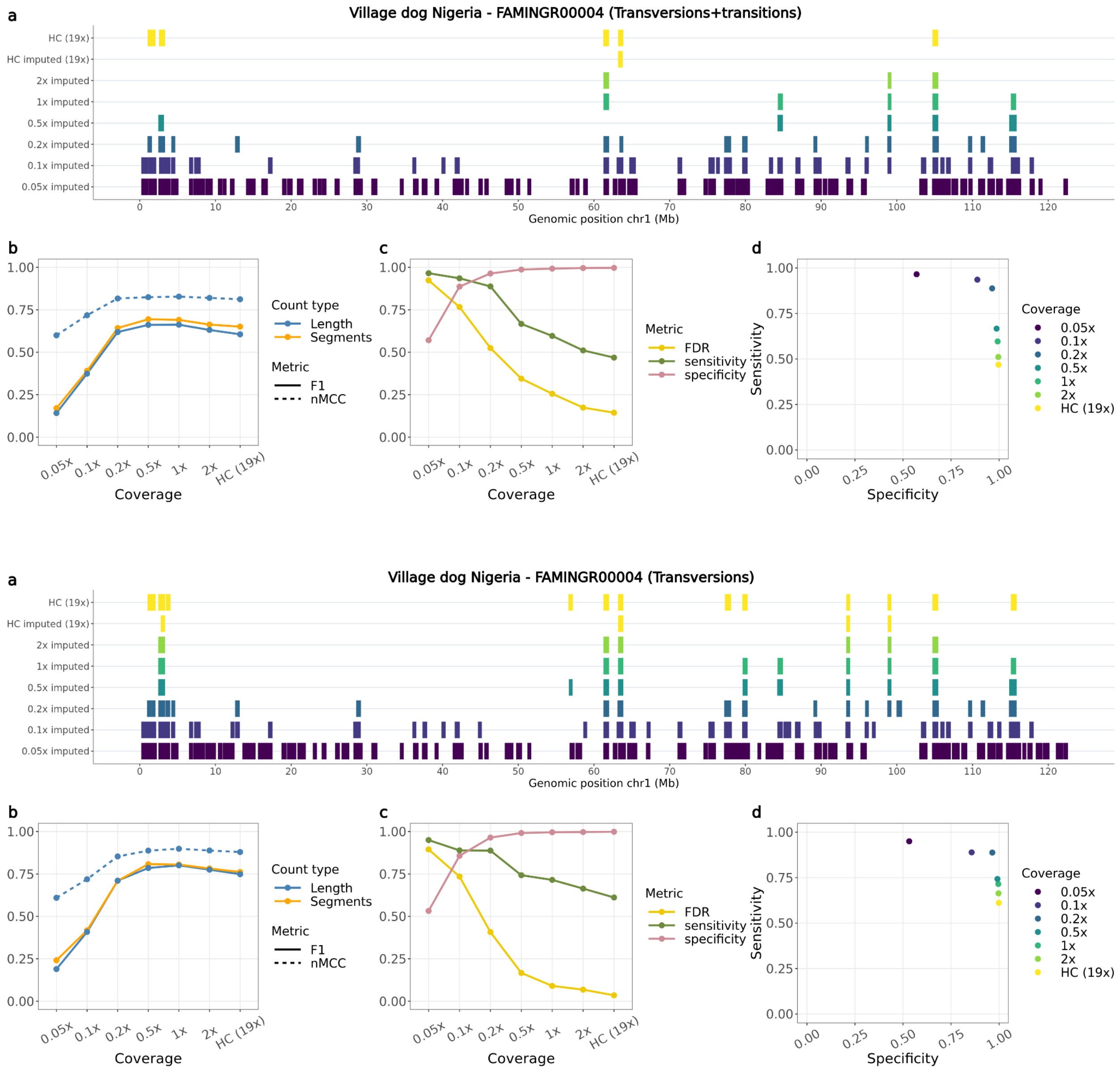


**Fig. S41:** *Overlap of ROH called from the FAMINGR00004 Nigerian Village dog for each imputed downsampled replicate using ROH estimates from the ground truth, using transversions and transitions (top panel) or only transversions (bottom panel). a) ROH called across the six tested coverages and the high coverage imputed and genotyped (ground truth) sample on chromosome one. b) Accuracy of recovering ROH across all tested coverages based on total length in bp (blue lines) and total number of segments (orange line) using the F1-score (solid line) and normalised Matthew correlation coefficient (nMCC) (dotted line). c) FDR, sensitivity and specificity measurements based on the total length of recovered ROH per coverage. d) Sensitivity plotted against specificity estimated based on the total length of recovered ROH across all tested coverages. HC: High coverage.*

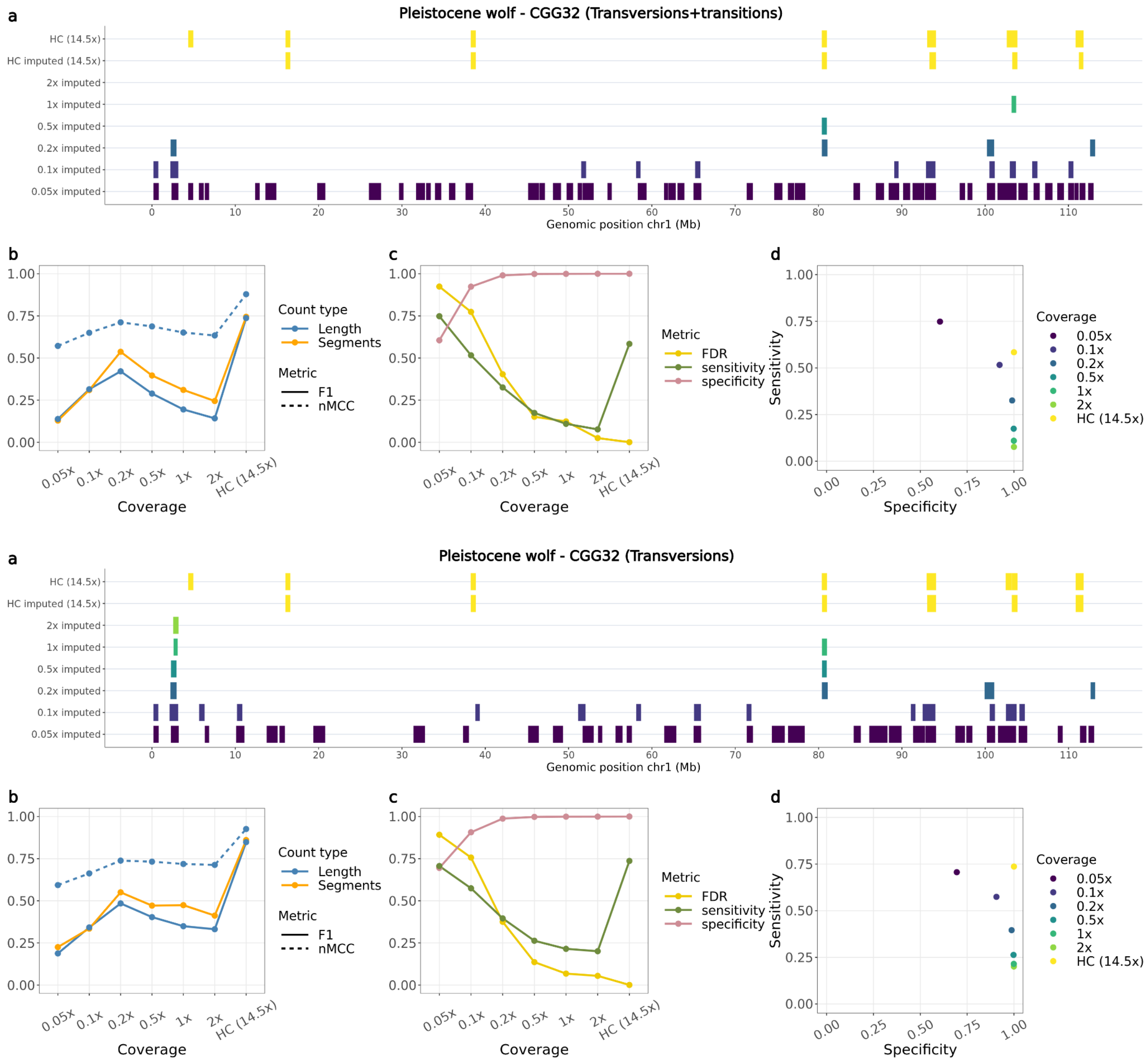


**Fig. S42:** *Overlap of ROH called from the CGG32 Pleistocene wolf for each imputed downsampled replicate using ROH estimates from the ground truth, using transversions and transitions (top panel) or only transversions (bottom panel). a) ROH called across the six tested coverages and the high coverage imputed and genotyped (ground truth) sample on chromosome one. b) Accuracy of recovering ROH across all tested coverages based on total length in bp (blue lines) and total number of segments (orange line) using the F1-score (solid line) and normalised Matthew correlation coefficient (nMCC) (dotted line). c) FDR, sensitivity and specificity measurements based on the total length of recovered ROH per coverage. d) Sensitivity plotted against specificity estimated based on the total length of recovered ROH across all tested coverages. HC: High coverage.*

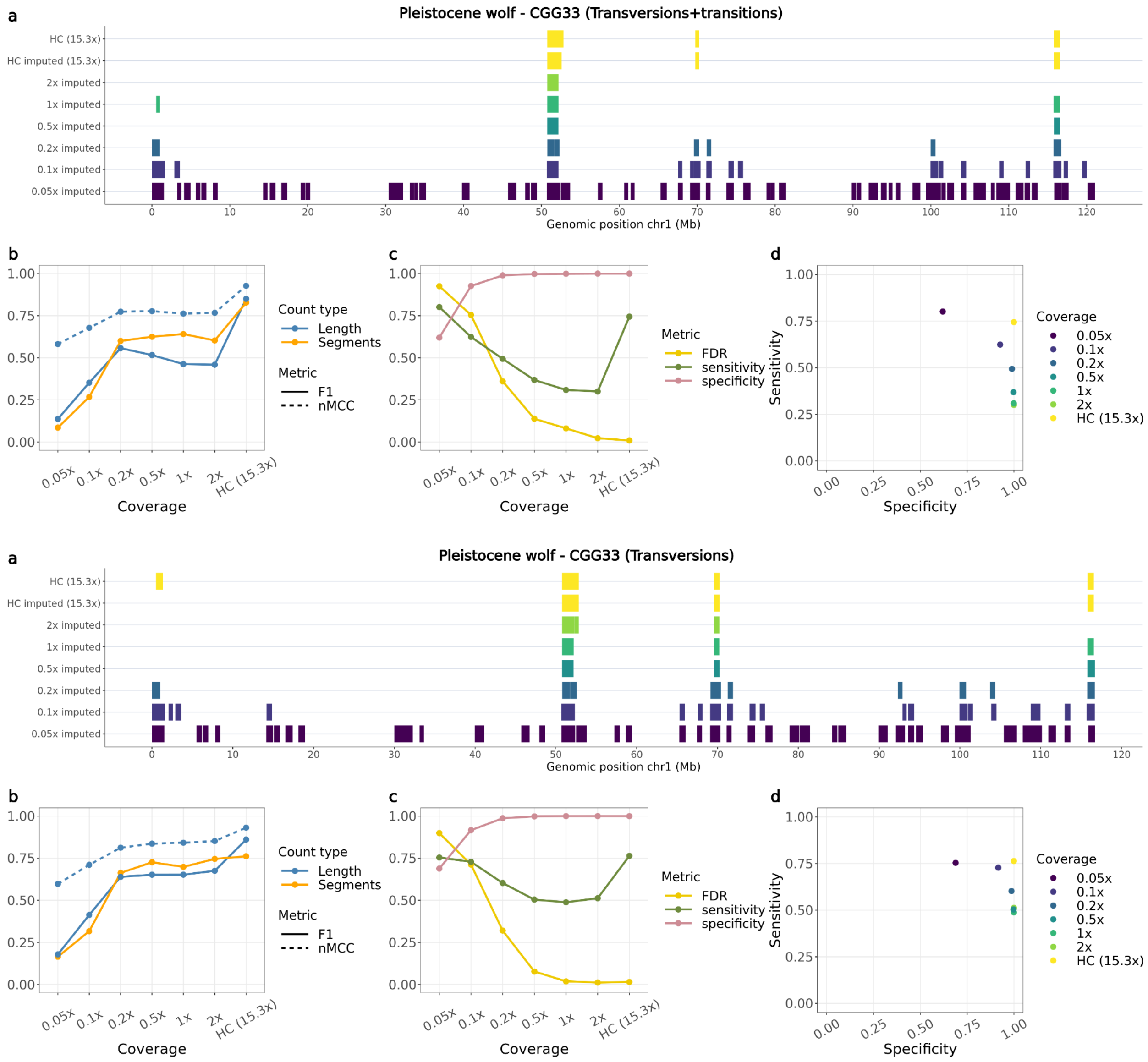


**Fig. S43:** *Overlap of ROH called from the CGG33 Pleistocene wolf for each imputed downsampled replicate using ROH estimates from the ground truth, using transversions and transitions (top panel) or only transversions (bottom panel). a) ROH called across the six tested coverages and the high coverage imputed and genotyped (ground truth) sample on chromosome one. b) Accuracy of recovering ROH across all tested coverages based on total length in bp (blue lines) and total number of segments (orange line) using the F1-score (solid line) and normalised Matthew correlation coefficient (nMCC) (dotted line). c) FDR, sensitivity and specificity measurements based on the total length of recovered ROH per coverage. d) Sensitivity plotted against specificity estimated based on the total length of recovered ROH across all tested coverages. HC: High coverage.*

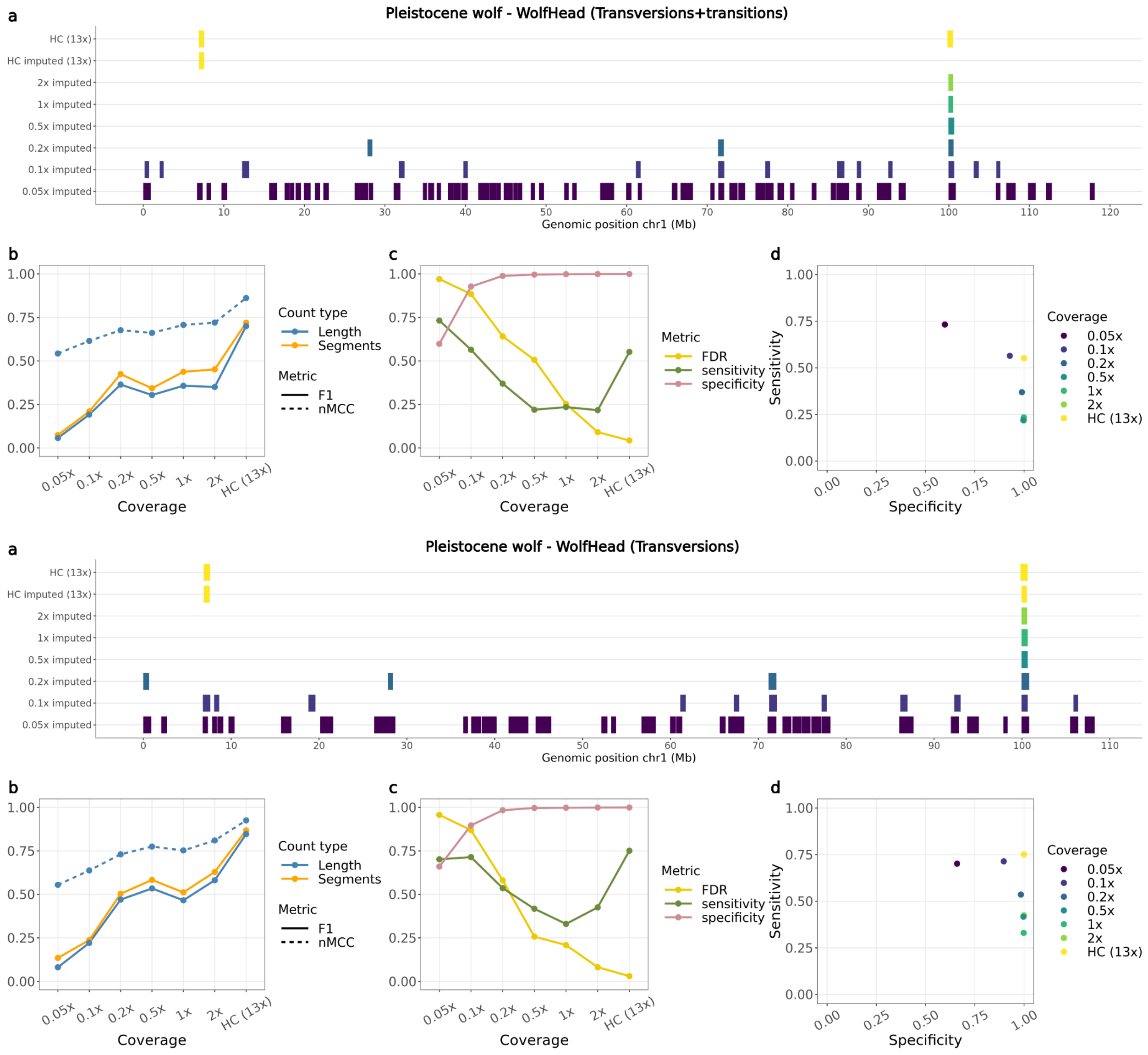


**Fig. S44:** *Overlap of ROH called from the WolfHead Pleistocene wolf for each imputed downsampled replicate using ROH estimates from the ground truth, using transversions and transitions (top panel) or only transversions (bottom panel). a) ROH called across the six tested coverages and the high coverage imputed and genotyped (ground truth) sample on chromosome one. b) Accuracy of recovering ROH across all tested coverages based on total length in bp (blue lines) and total number of segments (orange line) using the F1-score (solid line) and normalised Matthew correlation coefficient (nMCC) (dotted line). c) FDR, sensitivity and specificity measurements based on the total length of recovered ROH per coverage. d) Sensitivity plotted against specificity estimated based on the total length of recovered ROH across all tested coverages. HC: High coverage.*

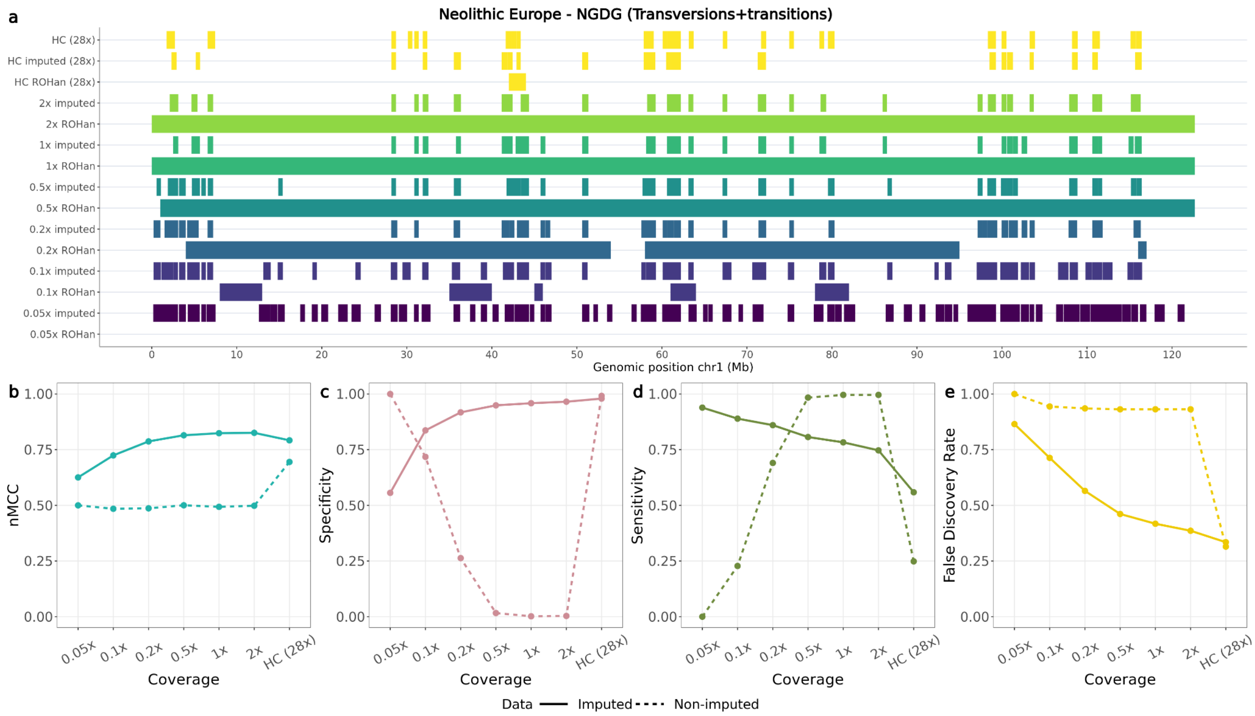


**Fig. S45:** *a) ROH called for the Newgrange Neolithic European dog across the high coverage and downsampled coverages (imputed and non-imputed) on chromosome one. b) Normalised Matthew correlation coefficient (nMCC), c) specificity, d) sensitivity and e) false discovery rate estimates based on the ROH inferred using the imputed (PLINK) or non-imputed (ROHan) samples, across all chromosomes and tested coverages. HC: High coverage.*

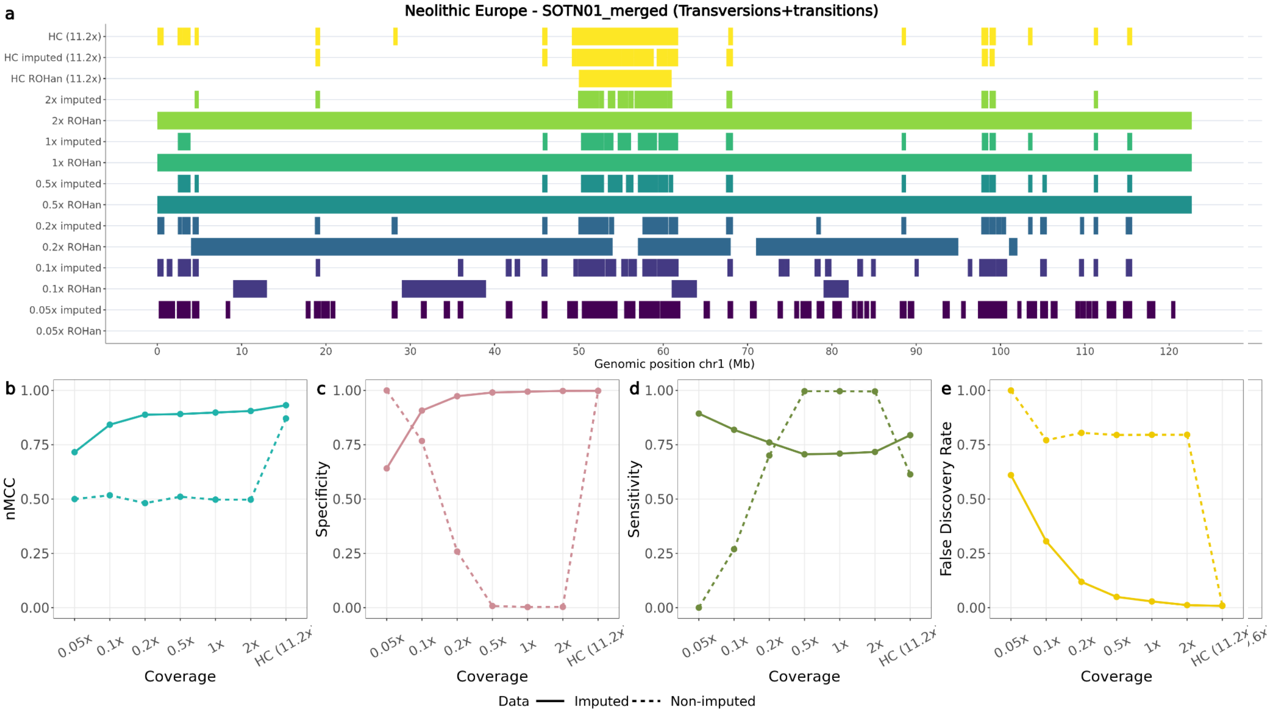


**Fig. S46:** *a) ROH called for the SOTN01 Neolithic European dog across the high coverage and downsampled coverages (imputed and non-imputed) on chromosome one. b) Normalised Matthew correlation coefficient (nMCC), c) specificity, d) sensitivity and e) false discovery rate estimates based on the ROH inferred using the imputed (PLINK) or non-imputed (ROHan) samples, across all chromosomes and tested coverages. HC: High coverage*

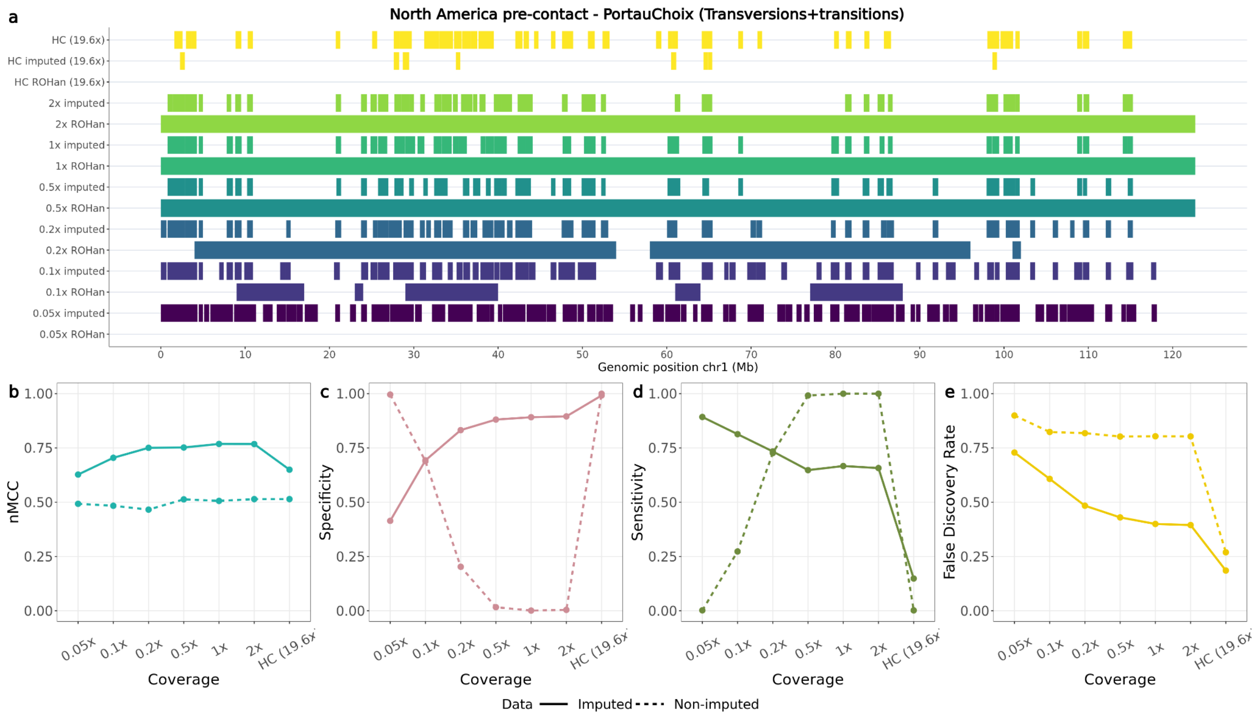


**Fig. S47:** *a) ROH called for the Port au Choix North American pre-contact dog across the high coverage and downsampled coverages (imputed and non-imputed) on chromosome one. b) Normalised Matthew correlation coefficient (nMCC), c) specificity, d) sensitivity and e) false discovery rate estimates based on the ROH inferred using the imputed (PLINK) or non-imputed (ROHan) samples, across all chromosomes and tested coverages. HC: High coverage*

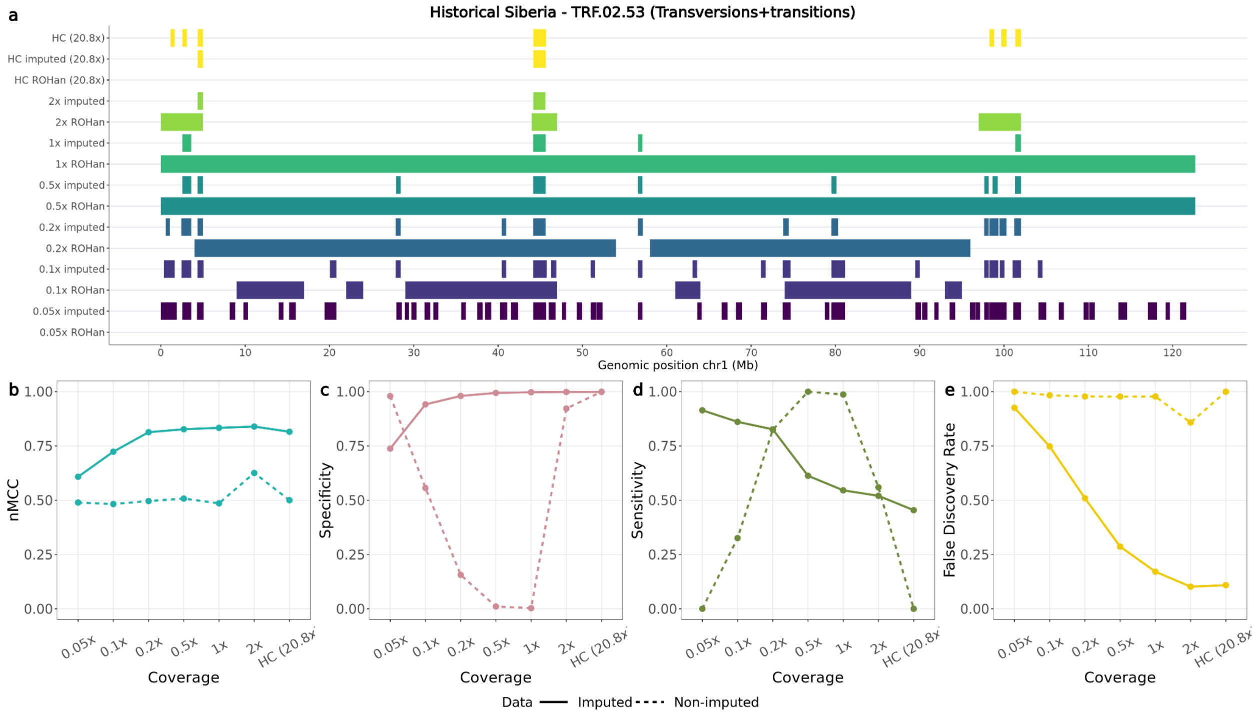


**Fig. S48:** *a) ROH called for the TRF.02.53 historical Siberian dog across the high coverage and downsampled coverages (imputed and non-imputed) on chromosome one. b) Normalised Matthew correlation coefficient (nMCC), c) specificity, d) sensitivity and e) false discovery rate estimates based on the ROH inferred using the imputed (PLINK) or non-imputed (ROHan) samples, across all chromosomes and tested coverages. HC: High coverage*

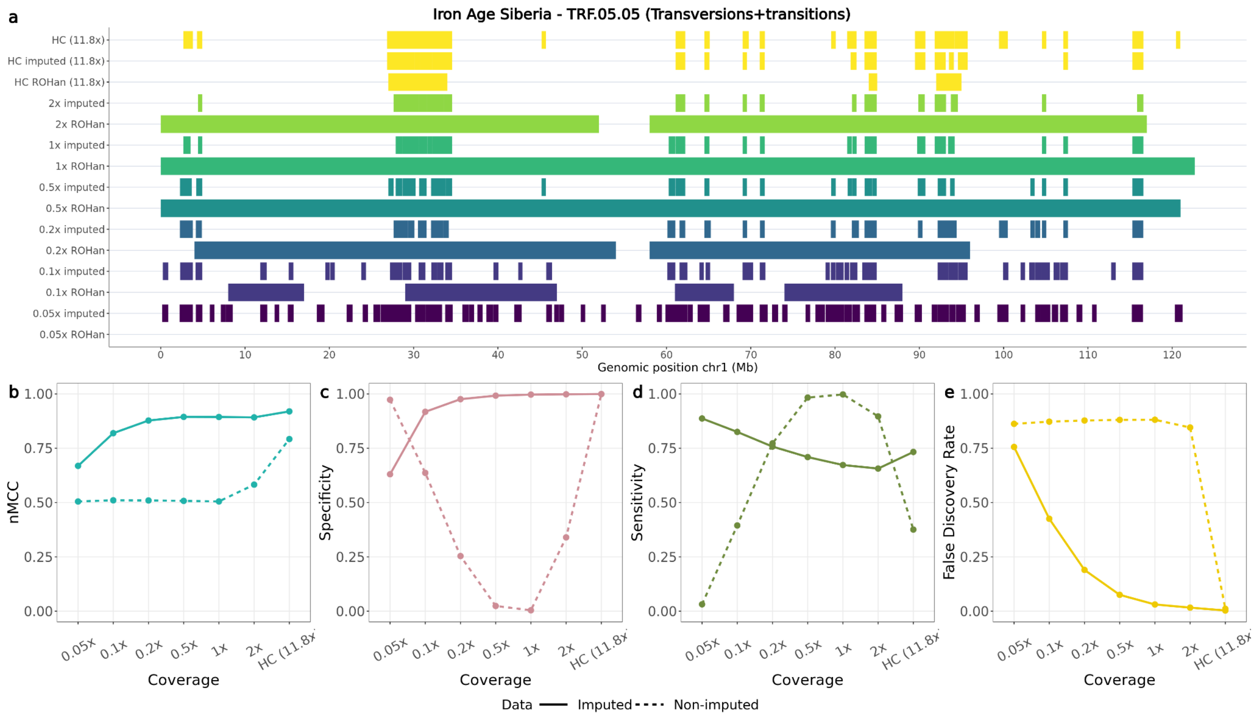


**Fig. S49:** *a) ROH called for the TRF.05.05 Iron Age Siberian dog across the high coverage and downsampled coverages (imputed and non-imputed) on chromosome one. b) Normalised Matthew correlation coefficient (nMCC), c) specificity, d) sensitivity and e) false discovery rate estimates based on the ROH inferred using the imputed (PLINK) or non-imputed (ROHan) samples, across all chromosomes and tested coverages. HC: High coverage*

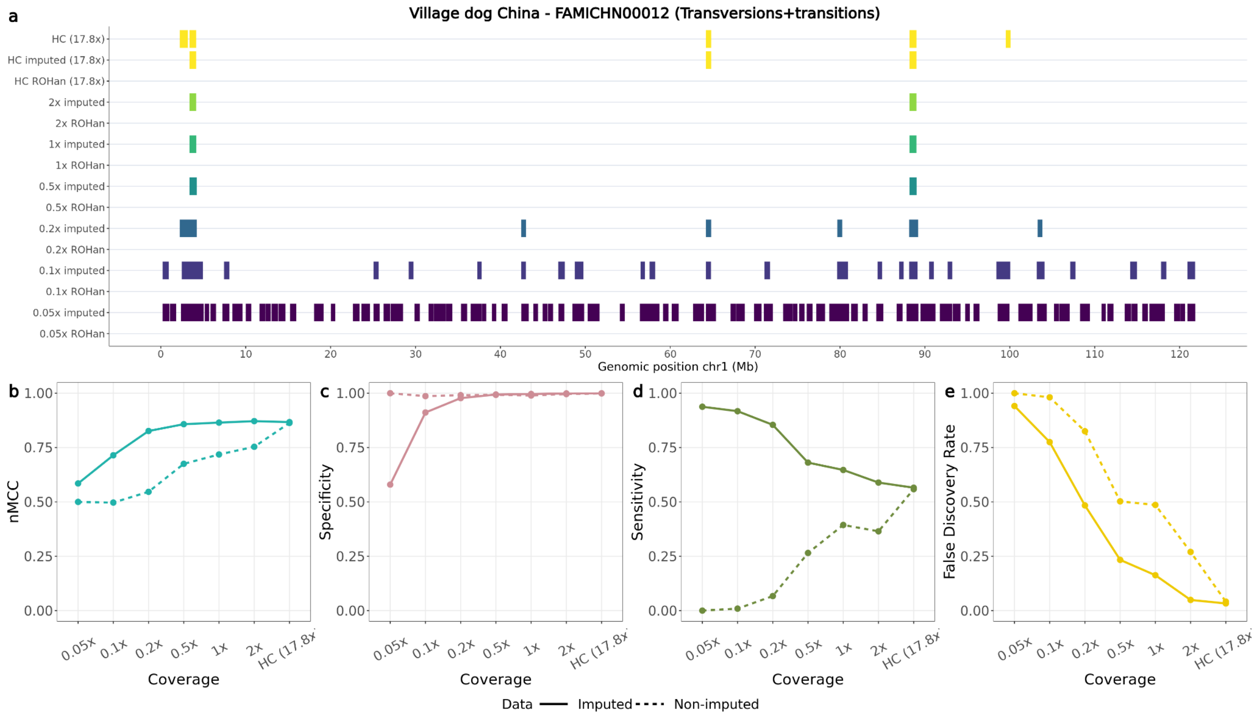


**Fig. S50:** *a) ROH called for the FAMICHN00012 Chinese Village dog across the high coverage and downsampled coverages (imputed and non-imputed) on chromosome one. b) Normalised Matthew correlation coefficient (nMCC), c) specificity, d) sensitivity and e) false discovery rate estimates based on the ROH inferred using the imputed (PLINK) or non-imputed (ROHan) samples, across all chromosomes and tested coverages. HC: High coverage*

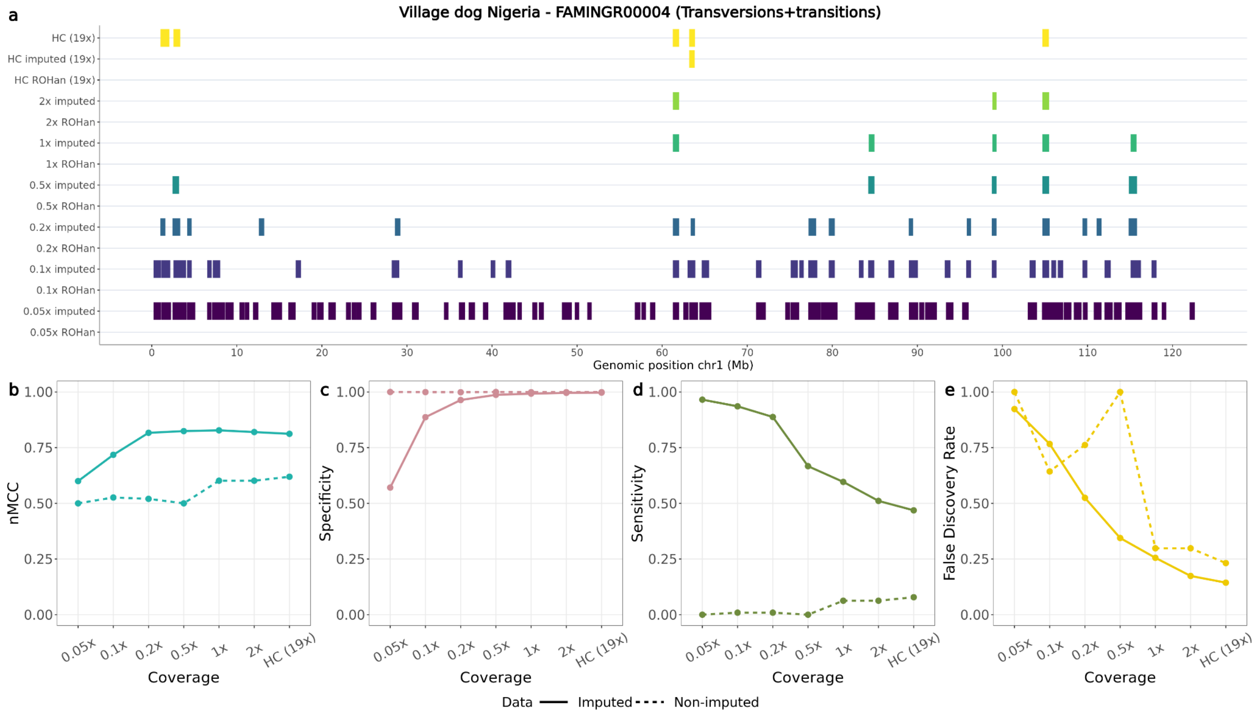


**Fig. S51:** *a) ROH called for the FAMINGR00004 Nigerian Village dog across the high coverage and downsampled coverages (imputed and non-imputed) on chromosome one. b) Normalised Matthew correlation coefficient (nMCC), c) specificity, d) sensitivity and e) false discovery rate estimates based on the ROH inferred using the imputed (PLINK) or non-imputed (ROHan) samples, across all chromosomes and tested coverages. HC: High coverage*

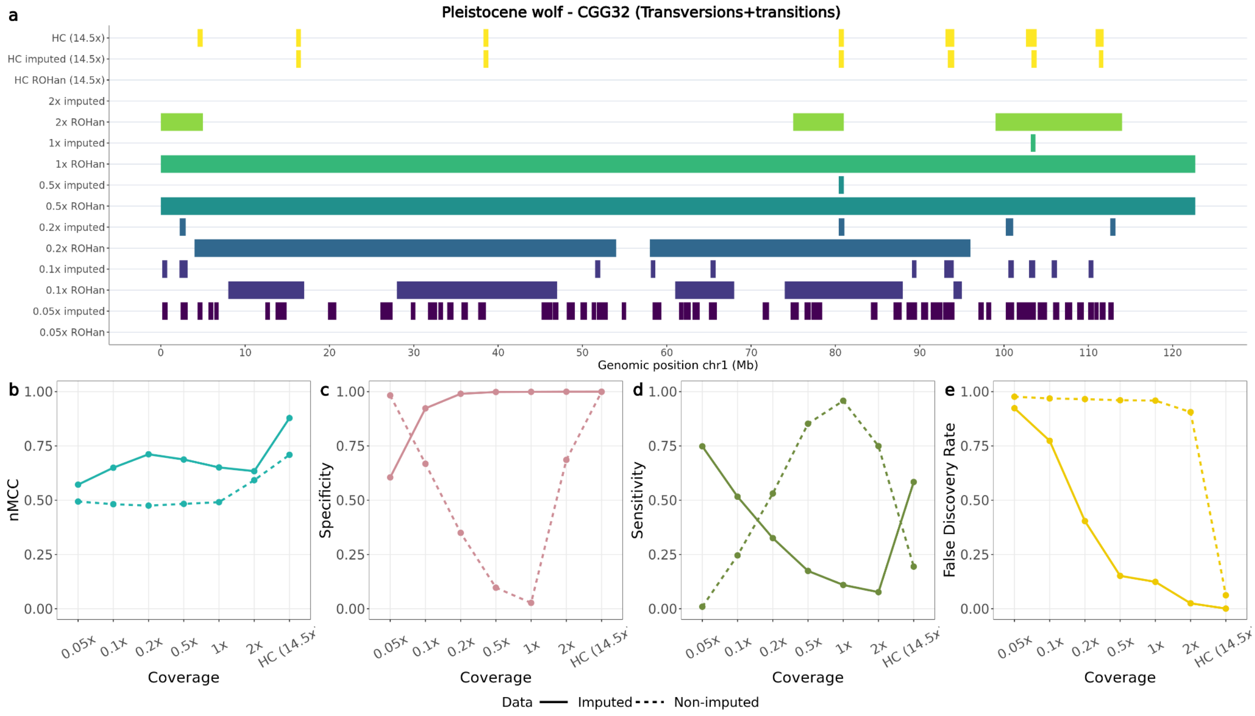


**Fig. S52:** *a) ROH called for the CGG32 Pleistocene wolf across the high coverage and downsampled coverages (imputed and non-imputed) on chromosome one. b) Normalised Matthew correlation coefficient (nMCC), c) specificity, d) sensitivity and e) false discovery rate estimates based on the ROH inferred using the imputed (PLINK) or non-imputed (ROHan) samples, across all chromosomes and tested coverages. HC: High coverage*

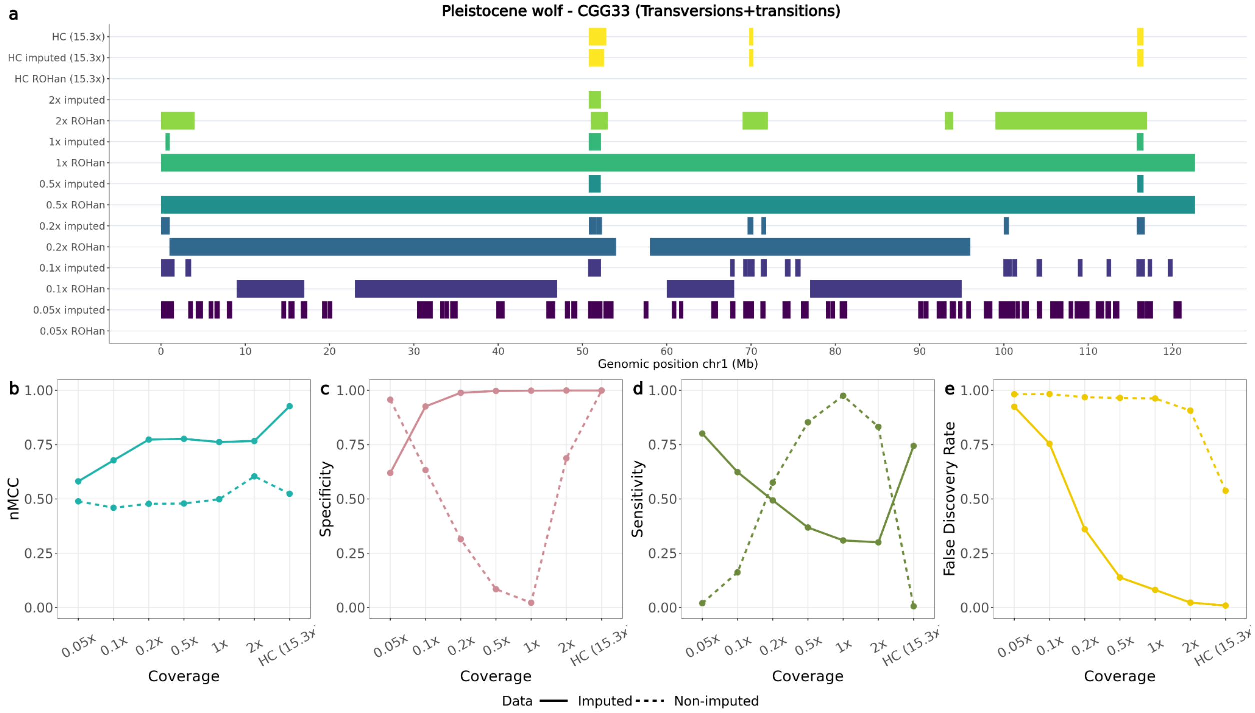


**Fig. S53:** *a) ROH called for the CGG33 Pleistocene wolf across the high coverage and downsampled coverages (imputed and non-imputed) on chromosome one. b) Normalised Matthew correlation coefficient (nMCC), c) specificity, d) sensitivity and e) false discovery rate estimates based on the ROH inferred using the imputed (PLINK) or non-imputed (ROHan) samples, across all chromosomes and tested coverages. HC: High coverage*

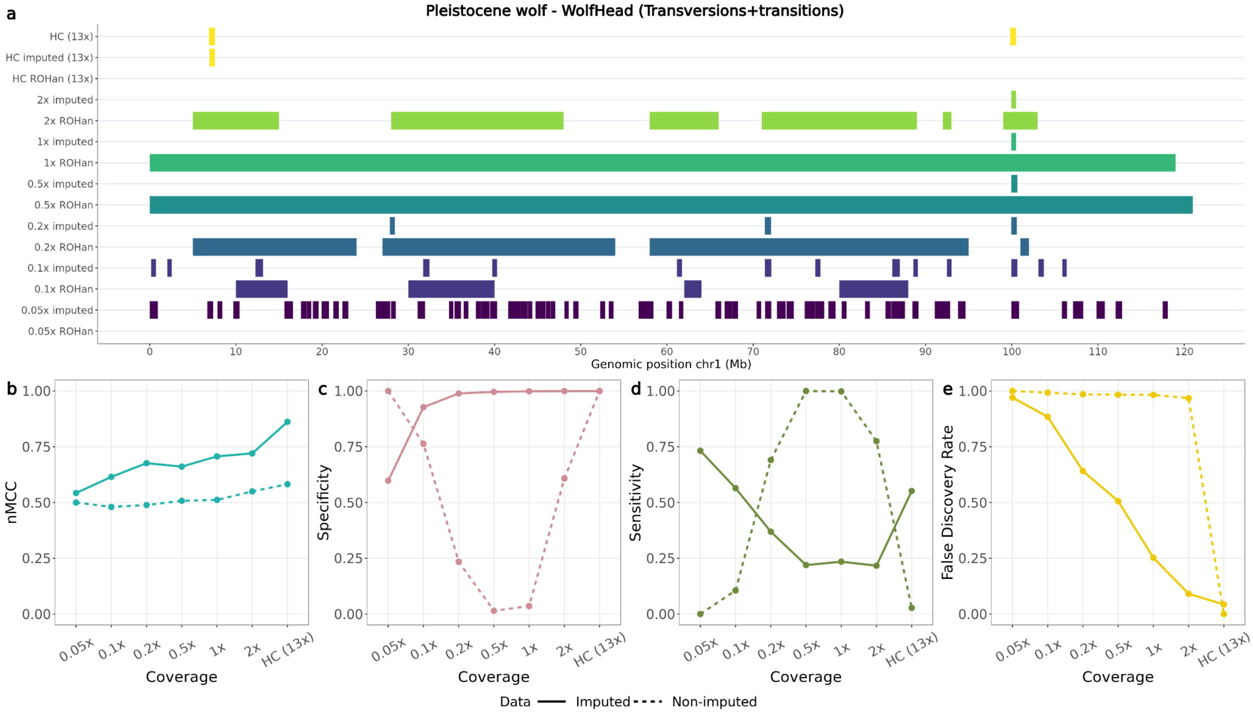


**Fig. S54:** *a) ROH called for the WolfHead Pleistocene wolf across the high coverage and downsampled coverages (imputed and non-imputed) on chromosome one. b) Normalised Matthew correlation coefficient (nMCC), c) specificity, d) sensitivity and e) false discovery rate estimates based on the ROH inferred using the imputed (PLINK) or non-imputed (ROHan) samples, across all chromosomes and tested coverages. HC: High coverage*

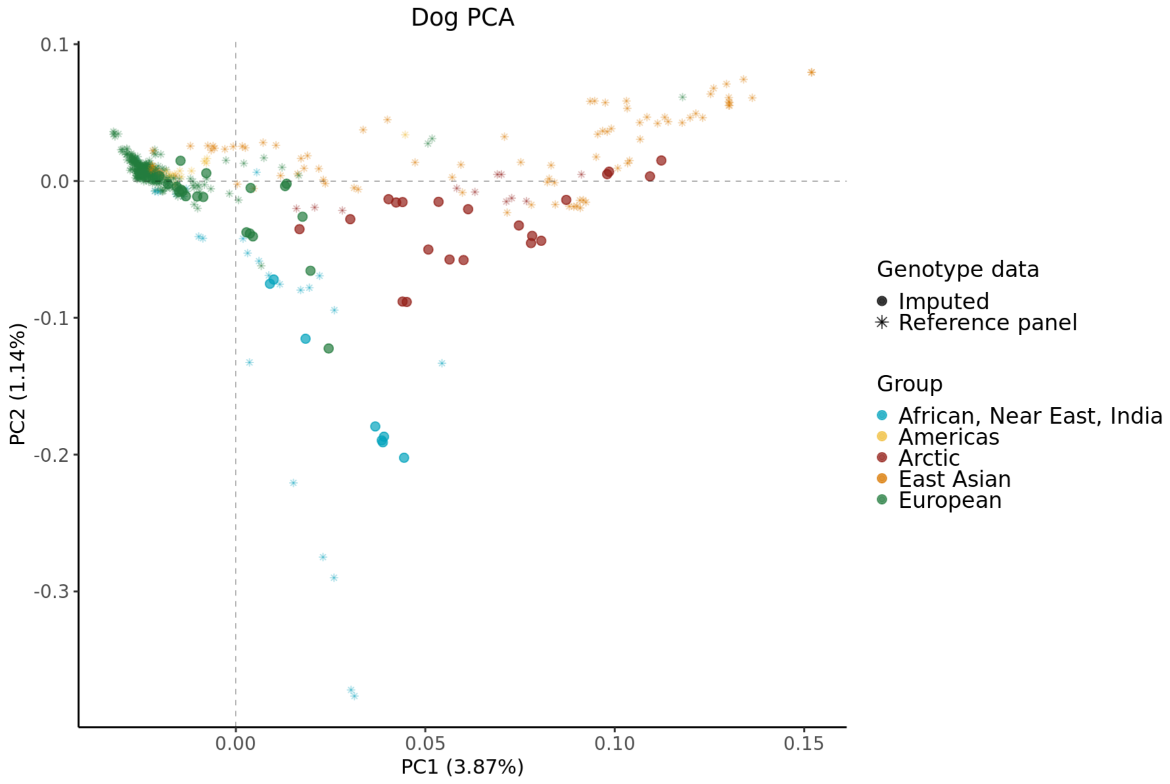
Dog PCA
PC2 (1.14%)
0.1
0.0
-0.1
-0.2
-0.3
0.00
0.05
0.10
0.15
PC1 (3.87%)
Genotype data
Imputed
Reference panel
Group
African, Near East, India
Americas
Arctic
East Asian
European

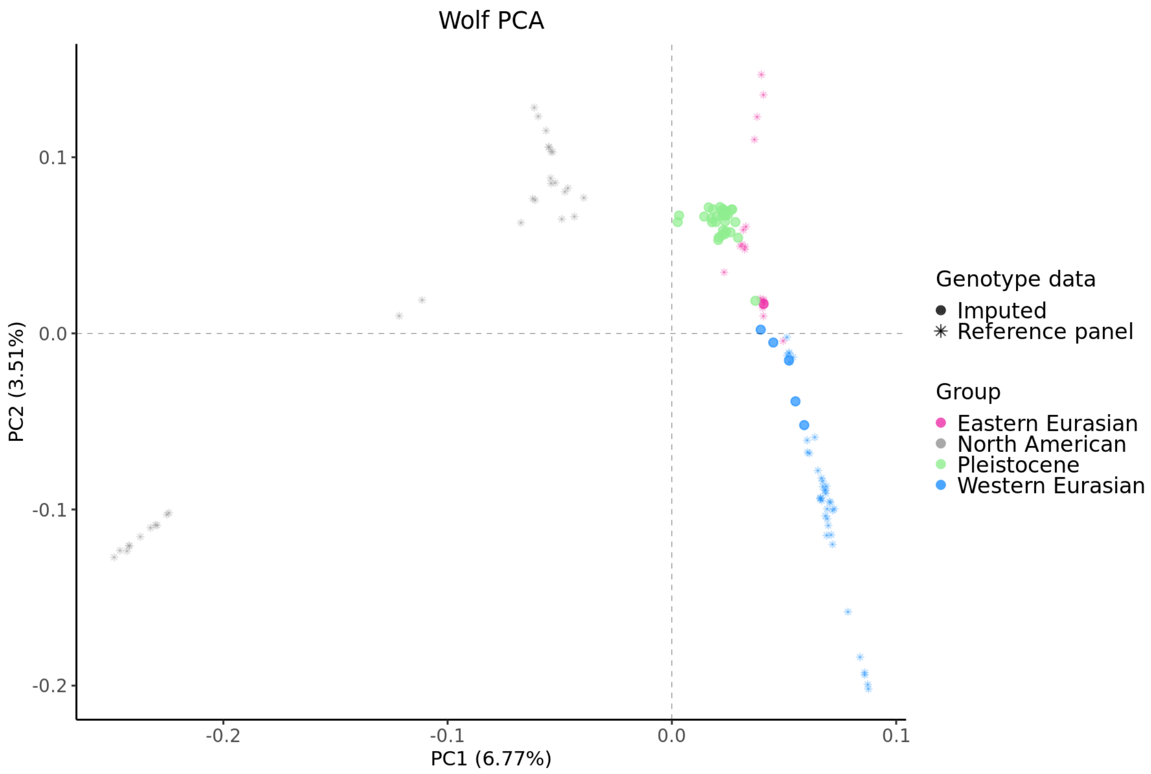
Wolf PCA
PC2 (3.51%)
0.1
0.0
-0.1
-0.2
-0.2
-0.1
0.0
0.1
PC1 (6.77%)
Genotype data
Imputed
Reference panel
Group
Eastern Eurasian
North American
Pleistocene
Western Eurasian

**Fig. S55:** *PCA of the imputed dogs (top plot) and wolves (bottom plot), alongside the dog or wolf samples from the reference panel. Both imputed and modern samples were used to create the PCs.*

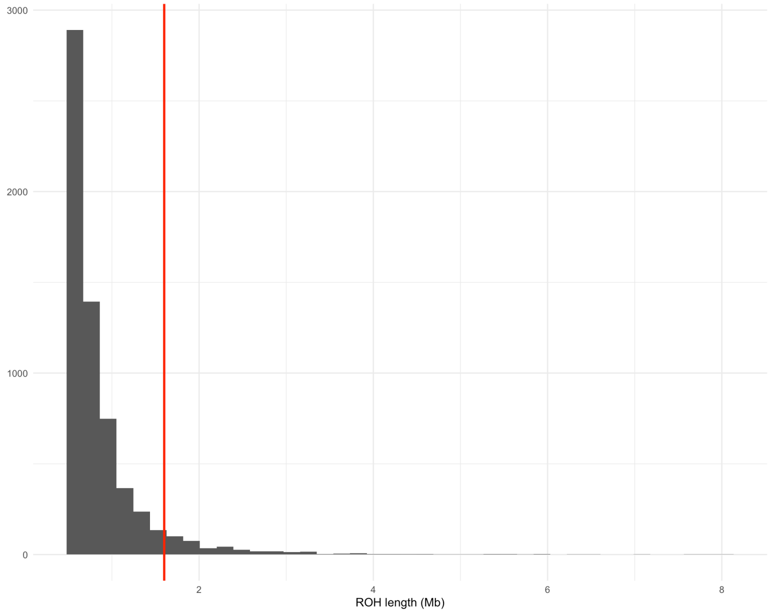


**Fig. S56***: Histogram of ROH lengths of imputed ancient dog samples. Red line is the 1.6Mb cutoff used to categorise short (<1.6Mb) and long (≥1.6Mb) ROH.*

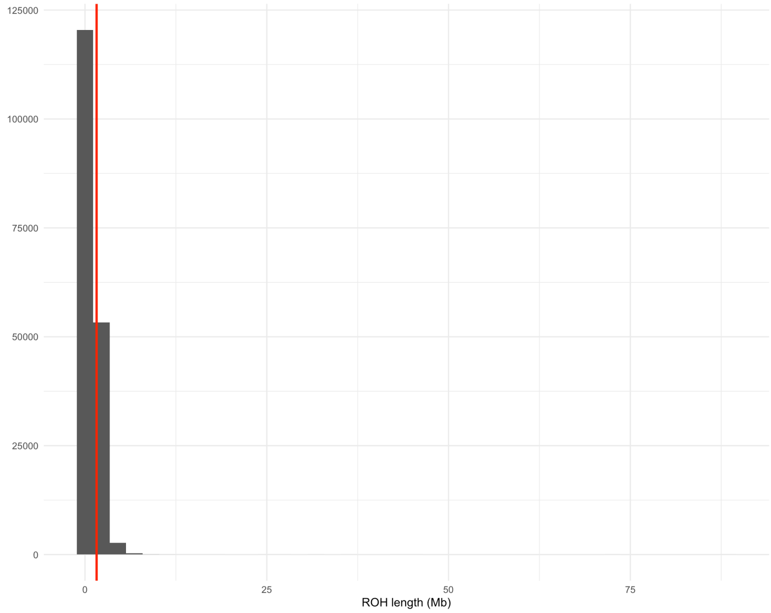


**Fig. S57***: Histogram of ROH lengths of modern dog samples. Red line is the 1.6Mb cutoff used to categorise short (<1.6Mb) and long (≥1.6Mb) ROH.*

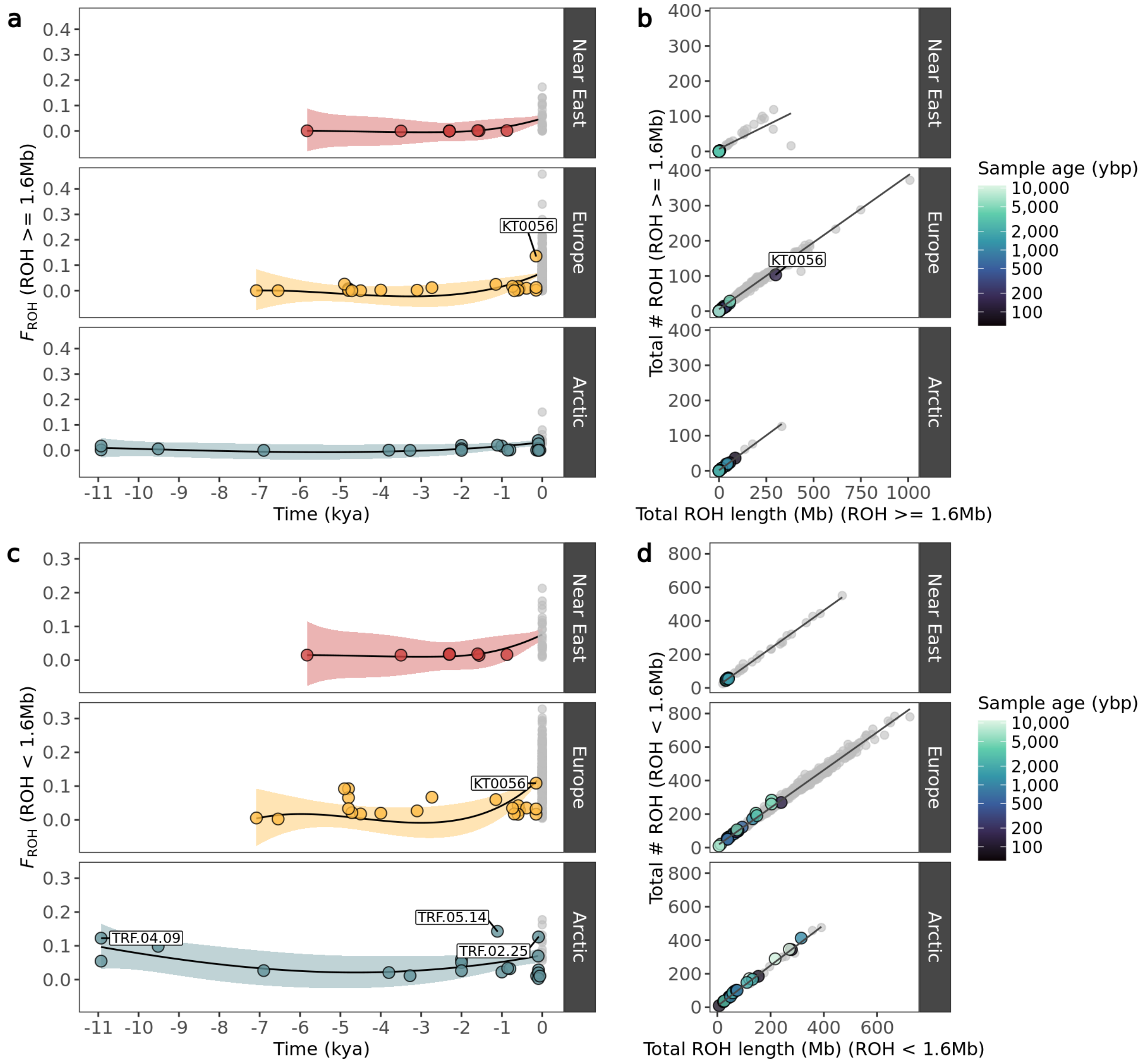


**Fig. S58:** *Left plots: Genomic inbreeding coefficient (FROH) of imputed and modern dogs as a function of time for a) ROH≥1.6Mb and c) ROH<1.6Mb. Imputed samples are coloured based on their geographic grouping, while modern samples are coloured in grey. Samples with $F_{ROH}$ values above 0.1 are indicated. Right plots: Total number of ROH segments plotted against total ROH length for the imputed dogs for b) ROH ≥ 1.6Mb and d) ROH < 1.6Mb. Colours correspond to age of imputed samples in years before present, while modern samples belonging to each dog group are coloured in grey.*

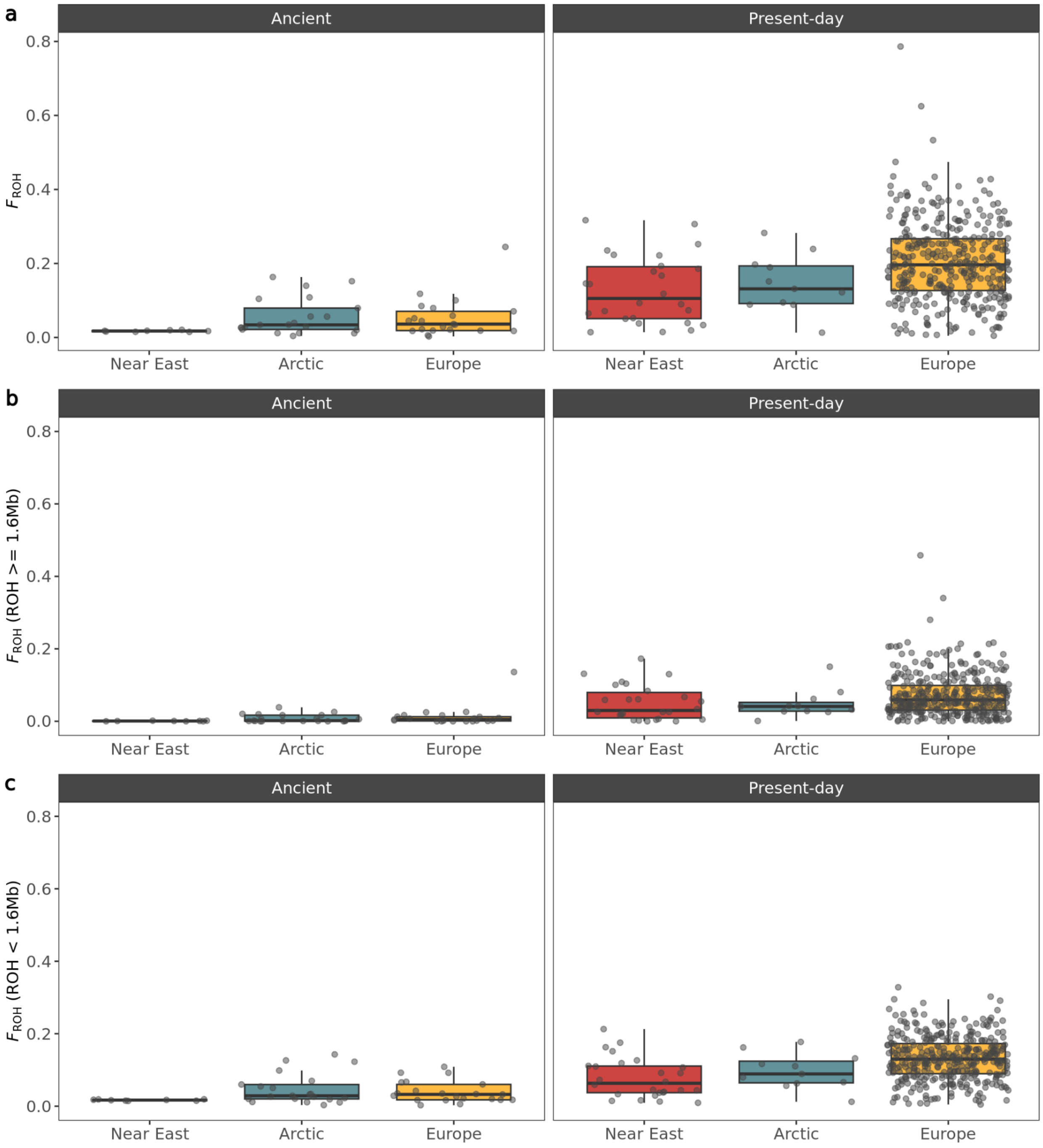


**Fig. S59:** *Boxplots of genomic inbreeding coefficients (FROH) from ancient and modern dogs for three ancestral groups for a) all ROH, b) ROH ≥ 1.6Mb and c) ROH < 1.6Mb. Horizontal lines represent the median.*

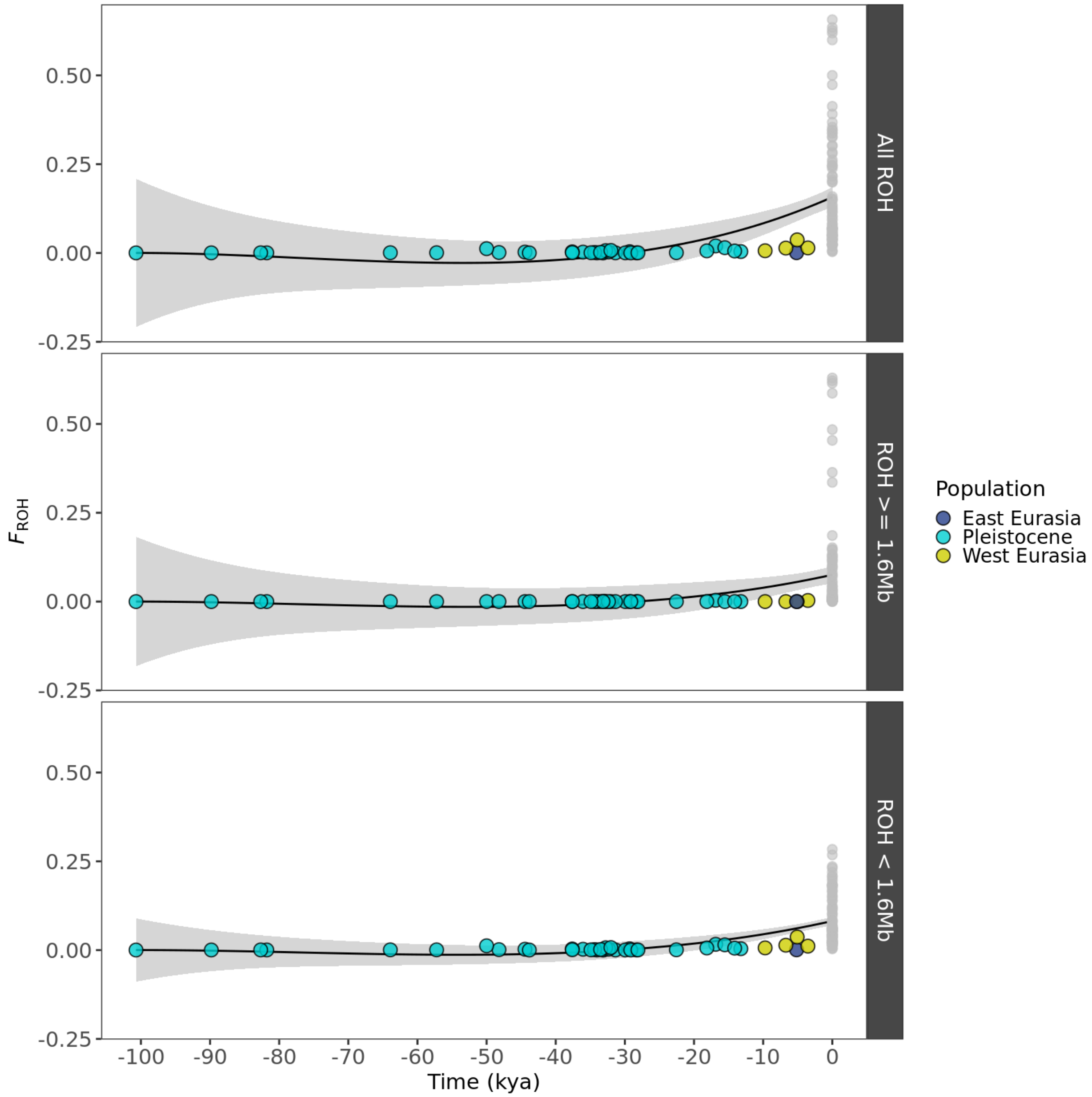


**Fig. S60:** *Genomic inbreeding coefficient (FROH) of imputed and modern wolves plotted as a function of time for all ROH, ROH ≥ 1.6Mb and ROH < 1.6Mb. Imputed samples are coloured based on their geographic grouping, while modern samples are coloured in grey.*

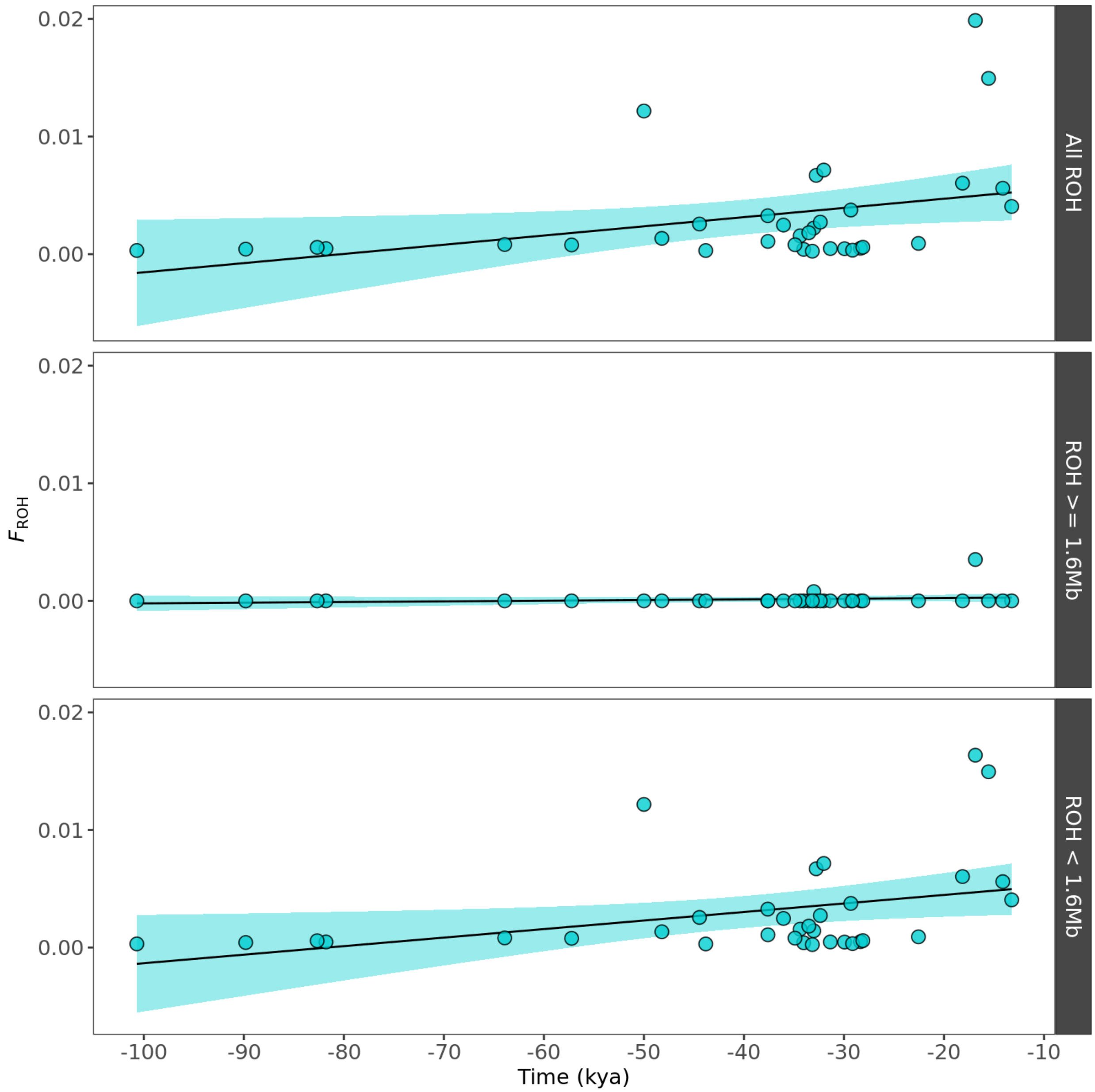


**Fig. S61:** *Genomic inbreeding coefficient (FROH) of imputed Pleistocene wolves plotted as a function of time for i) all ROH, ii) ROH ≥ 1.6Mb and iii) ROH < 1.6Mb.*

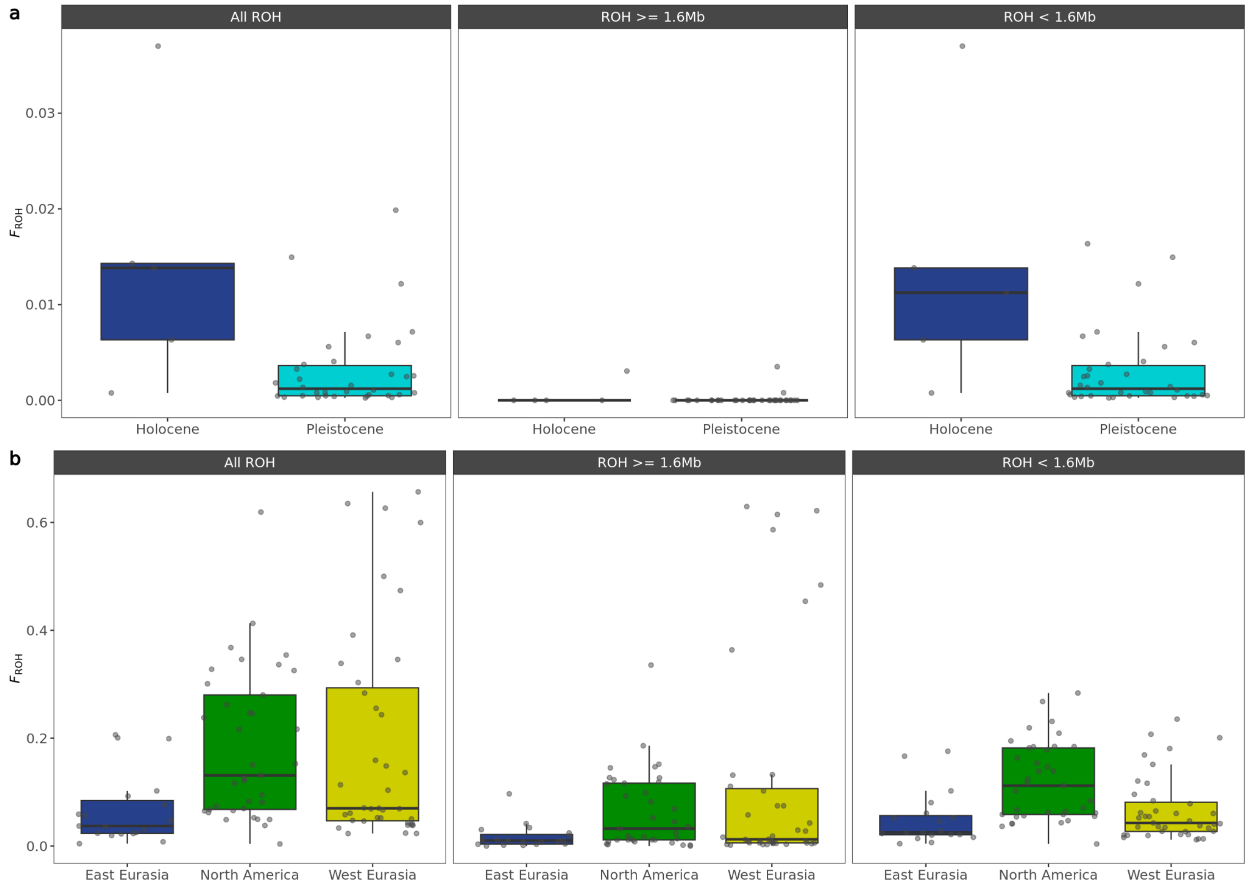


**Fig. S62:** *Boxplots of genomic inbreeding coefficients (FROH) from a) ancient and b) present-day wolves for all ROH, ROH ≥ 1.6Mb and ROH < 1.6Mb. Horizontal lines represent the median.*

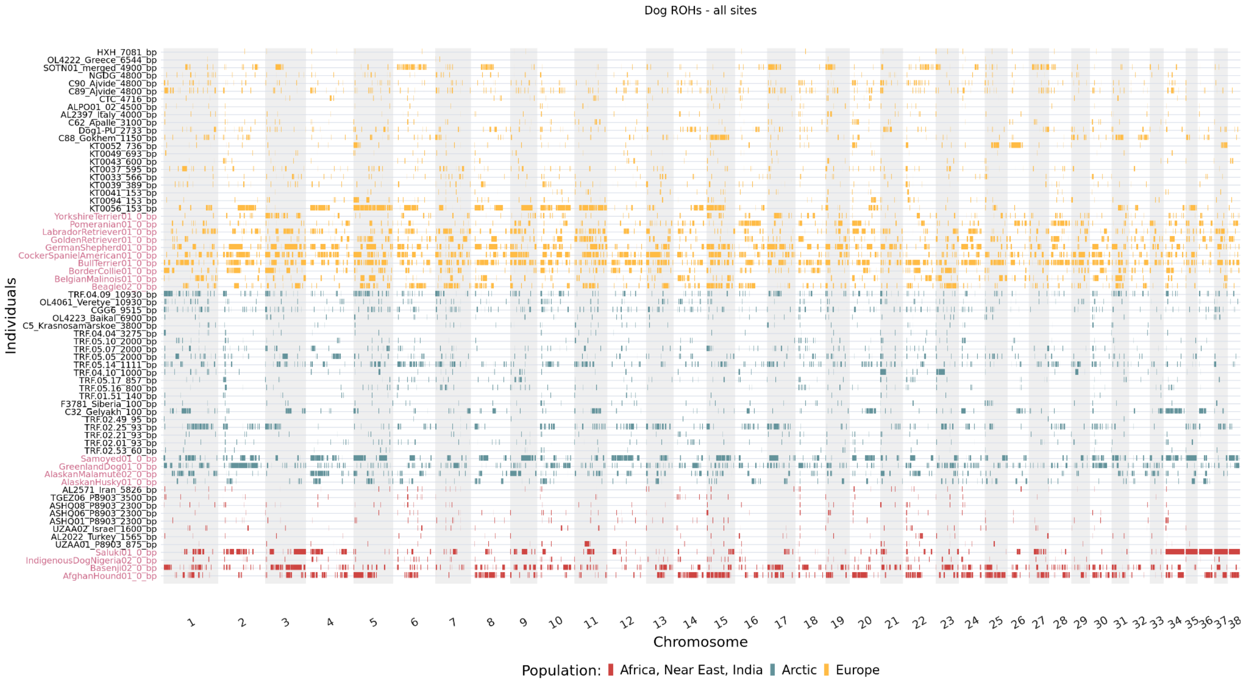


**Fig. S63:** *ROH across the genome for each imputed dog sample estimated based on transversions and transitions. ROH bands are coloured based on geographic origin and ordered based on age within each geographic region. ROH for a subset of modern samples are added for comparison and highlighted with red text.*

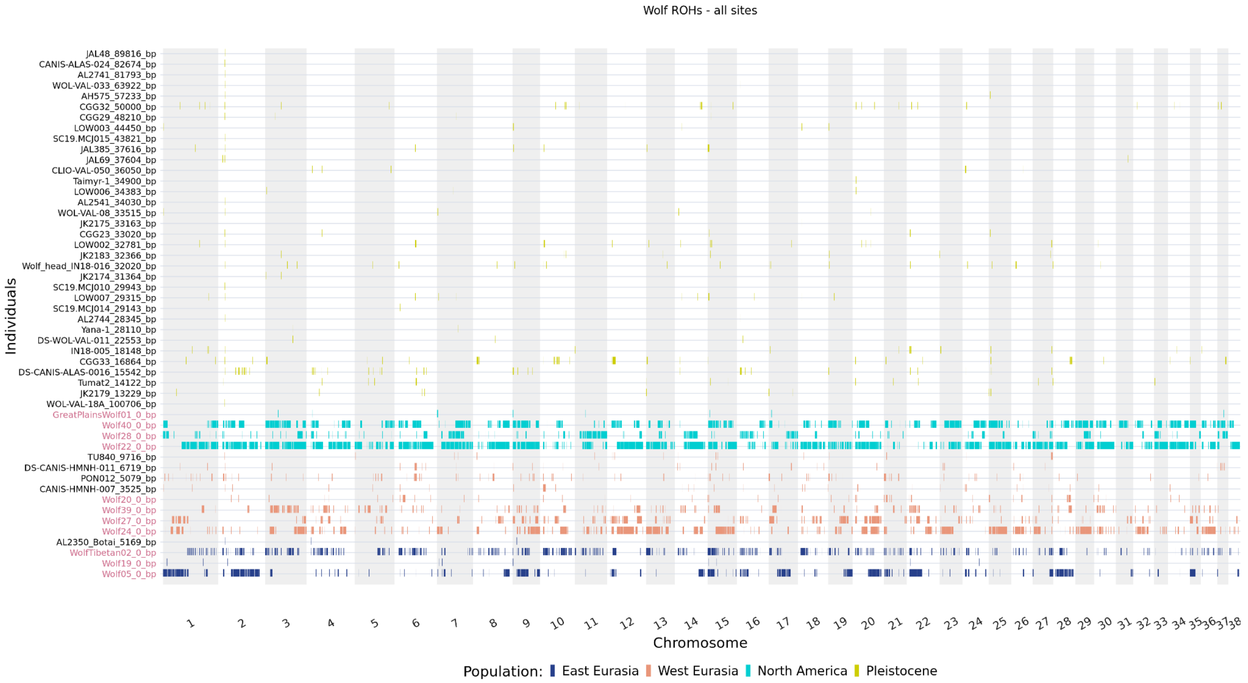


**Fig. S64:** *ROH across the genome for each imputed wolf sample estimated based on transversions and transitions. ROH bands are coloured based on geographic origin and ordered based on age within each*

*geographic region. ROH for a subset of modern samples are also added for comparison and highlighted with red text.*

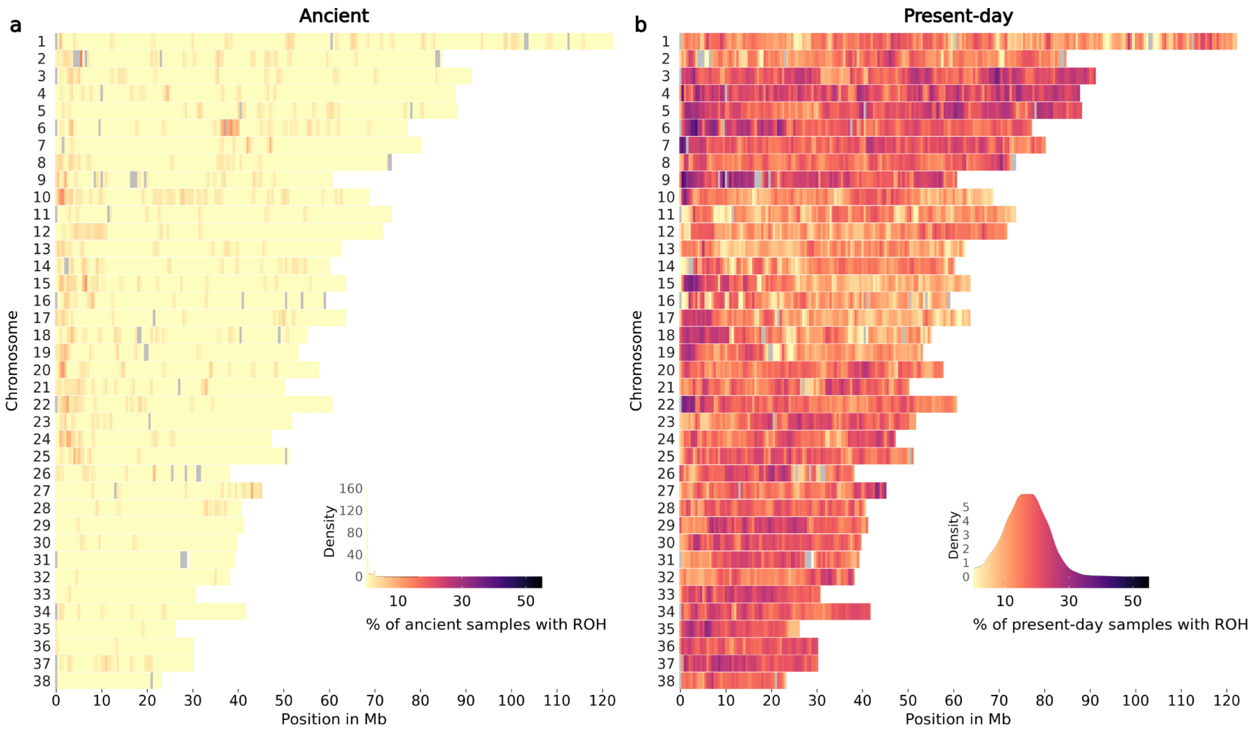


**Fig. S65:** *ROH across all chromosomes of a) ancient wolves and b) present-day wolves. The colour legend represents the % of samples which have an ROH at each genomic position, with more yellow regions representing ROH deserts and more purple regions representing ROH islands. Grey coloured regions indicate windows with an average depth of coverage estimated from all ancient dog samples above or below the mean ± 2*std.*

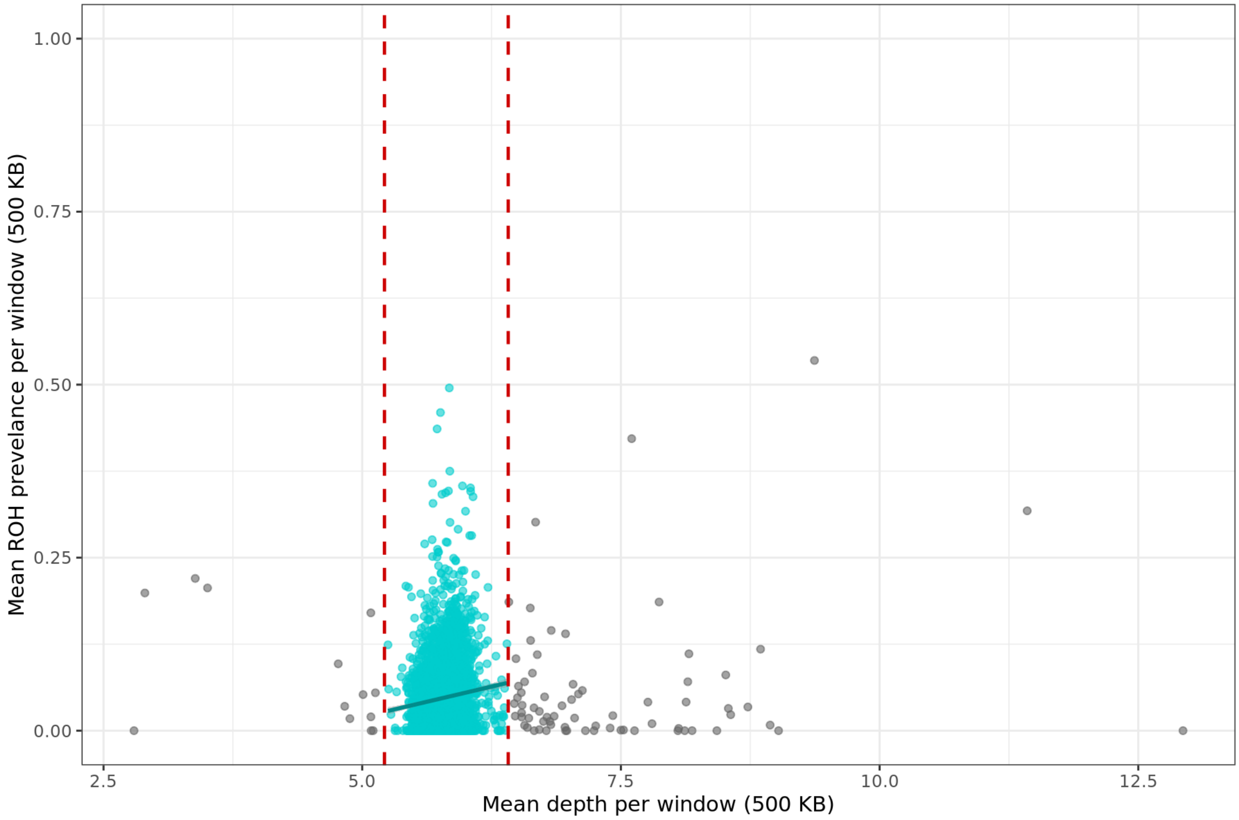


**Fig. S66:** *Mean ROH prevalence per 500Kb genomic window against the mean depth of each window estimated from 50 ancient dogs. Windows with depth above or below the mean ± 2*std were not included in estimating ROH prevalence across the genome of ancient and modern dogs.*

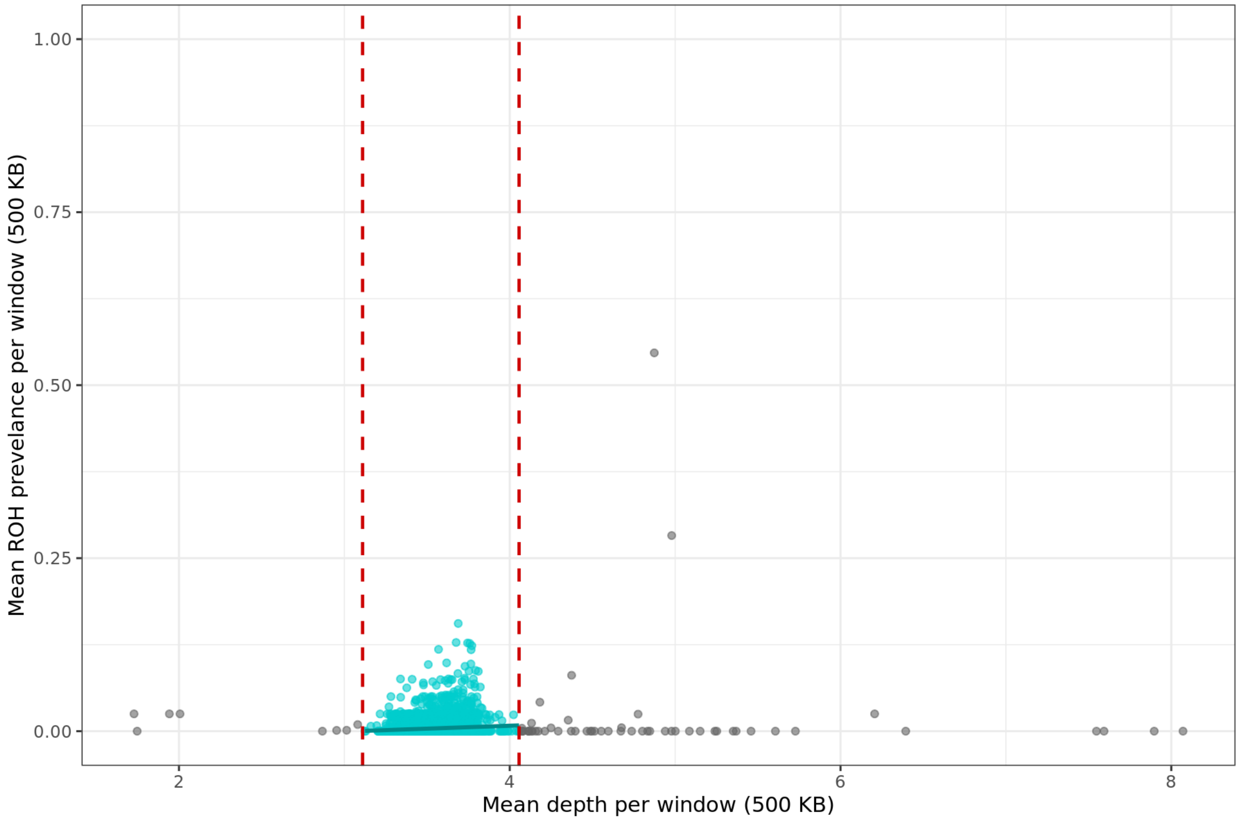


**Fig. S67:** *Mean ROH prevalence per 500Kb genomic window against the mean depth of each window estimated from 40 ancient wolves. Windows with depth above or below the mean ± 2*std were not included in estimating ROH prevalence across the genome of ancient and modern wolves.*
